# Hypothalamic Cell Type Evolution across Vertebrates

**DOI:** 10.64898/2026.08.23.746350

**Authors:** Yunming Wu, Prateek Kalakuntla, Billie C. Goolsby, Yanay Rosen, Alyssa Hayashi, Guang Yang, Chew Chai, Heidi M. Tate, Tom Hindmarsh Sten, Jessica Nowicki, Lindsay L. Sailer, Jun H Song, Jaeyoon Lee, Robert C. Jones, Dong-Wook Kim, David Kabelik, Alexander G. Ophir, Hongkui Zeng, Jure Leskovec, Stephen R. Quake, Lauren A. O’Connell, Bo Wang, Liqun Luo

**Author notes:** These authors contributed equally to this work (Yunming Wu, Prateek Kalakuntla, Billie C. Goolsby). First authors mutually agree to an interchangeable order.

## Abstract

The hypothalamus regulates vital functions and is highly conserved among vertebrates occupying diverse niches, but little is known about the evolution of its cell types. Using comparative single-cell transcriptomics across major vertebrate classes, we identified deeply conserved and clade-specific cell types. While homologous cell types express conserved transcription factors and neuropeptides, their regulatory linkages are often rewired across evolution. The paraventricular nucleus largely retains its neuropeptide expression and responds to dehydration with fine cellular resolution from amphibians to mammals, whereas lamina terminalis neurons are specific to terrestrial vertebrates and exhibit gene expression divergence in cell types activated by dehydration and heat. We propose that a conserved set of keystone cell types sustain essential functions across vertebrates, while flexible gene regulatory programs alongside numerous clade-specific cell types offer evolutionary fluidity.

## Main text

The hypothalamus is an ancient brain structure present in all vertebrates (*1*–*3*). It regulates many vital functions, including fluid intake and osmotic balance (*4*–*7*), feeding and energy balance (*8*, *9*), body temperature (*10*, *11*), social behavior (*12*–*15*), circadian rhythms (*16*), and sleep (*17*, *18*). Yet it remains unclear how the hypothalamic cell types underlying these life-sustaining physiological functions have been conserved or diversified among vertebrates occupying diverse niches.

We compared hypothalamic cell types across select species of mammals, birds, reptiles, amphibians, and teleosts (ray-finned fish) using single-cell transcriptomics. We identified a core set of cell types that are conserved across the five vertebrate classes, including many with well-characterized physiological functions in the mouse such as the dehydration-responsive neurons in the paraventricular hypothalamic nucleus (PVH). Alongside these, we found numerous clade-specific cell types, including the mammalian-specific neurons of the mammillary nuclei known to support spatial memory (*19*), and the lamina terminalis cell types associated with terrestrial vertebrates, which regulate body osmolality and temperature (*6*). Even among deeply conserved cell types, regulatory relationships between transcription factors (TFs) and their targets can undergo substantial turnover. Using spatial transcriptomic profiling in amphibians, birds, and mammals, we showed that conserved cell types in two hypothalamic nuclei retained similar responses to physiological stress even when the canonical marker genes used to define these nuclei diverged. We propose that cell type identity and function are anchored by conserved transcriptional regulators and neurotransmitter effectors, whereas the rewiring of gene regulatory networks, together with the many clade-specific cell types, provides the flexibility for species-specific adaptations.

### High diversity of hypothalamus cell types across vertebrates

To sample hypothalamic cell types across major vertebrate clades (**Fig. 1A**), we performed single-nucleus RNA-sequencing (snRNA-seq) on the hypothalamus of tropical clawed frogs (*Xenopus tropicalis*) representing amphibians, green anoles (*Anolis carolinensis*) representing reptiles, Japanese quail (*Coturnix japonica*) representing birds, and prairie voles (*Microtus ochrogaster*) representing mammals. We chose these species for their well-annotated genomes and relative availability. We cryodissected the hypothalamus and used the same sample preparation protocol to minimize technical variation across species (**Fig. 1B; figs. S1** to **S3**). Following quality control (**Methods**), we obtained a total of 205,264 high-quality neuronal nuclei with a median detection depth of 2,038 genes per cell (2,191 in prairie vole, 2,389 in Japanese quail, 1,564 in green anole, and 1,751 in *Xenopus*). Complete analyses of sequencing depth and cell quality in each species can be found in **figs. S4** to **S8.** We also analyzed published hypothalamus single-cell RNA-seq datasets of zebrafish (*Danio rerio*) (*20*) representing teleosts.

**Fig. 1.**
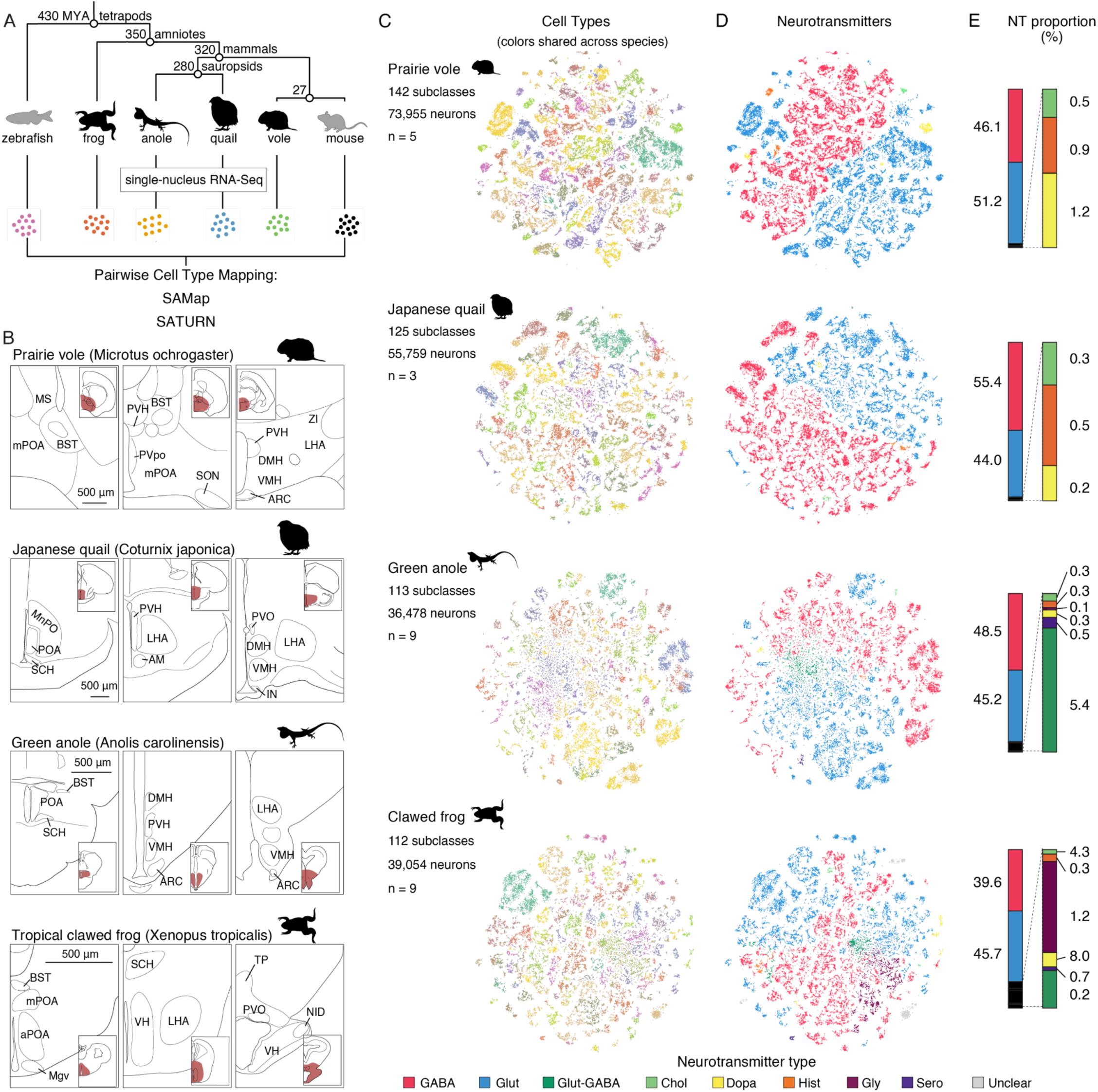
snRNA-seq of 205,264 neurons across the hypothalamus of prairie voles, Japanese quails, green anoles, and tropical clawed frogs. **(A)** Phylogenetic relationships of species analyzed. Data from zebrafish and mouse (gray) were obtained from prior studies (*20*, *21*). Numbers at branches indicate divergence times (millions of years ago). **(B)** Representative anatomical annotation of the hypothalamus in prairie vole (*120*, *121*), Japanese quail, green anole (*122*), tropical clawed frog using three coronal sections from anterior (left) to posterior (right) of one hemisphere. We used both the published reference and our own reference for anole. See detail and anatomical abbreviations in **figs. S1** to **S3**. Insets are zoom-out corresponding brain sections, with hypothalamic regions highlighted in red. **(C)** t-SNE (t-distributed Stochastic Neighbor Embedding) plots of neuronal transcriptomes across the four sequenced species. Cells are colored by subclass; cells sharing a color across species belong to homologous subclasses (**Fig. 2**). The numbers of sequenced neurons that pass quality control, subclasses, and animals used are listed to the left of each tSNE plot. **(D)** tSNE plots colored by neurotransmitter usage. Thresholds for neurotransmitter type classification are in Methods. **(E)** Proportions of all neurotransmitter types (NT, left column) and proportions among non-glutamatergic/non-GABAergic neurons (expanded in the right column). Neurons with unclear neurotransmitter expression were excluded from this analysis.

For cross-species comparisons, we leveraged a mouse (*Mus musculus*) whole brain single-cell atlas as a reference map (*21*), which hierarchically classified cell types at four levels of increasing resolution: classes— separating brain regions and neurotransmitter types; subclasses—largely corresponding to neural populations with shared spatial patterns that approximate nuclei in the hypothalamus; supertypes—which we found to be functionally distinct in some hypothalamic regions; and clusters—the finest transcriptomic grouping of cells. We performed cross-species comparisons at the subclass and supertype levels, as these are most relevant to the brain structures and functions that we analyzed.

Through cross-species mapping (**fig. S9**), we determined the clustering levels that corresponded to subclasses in the vole, quail, anole, *Xenopus* (**Fig. 1C**), and zebrafish (**fig. S8**). We classified the neurotransmitter identity of these subclasses using the expression of well-studied biosynthetic enzymes and neurotransmitter transporters (**Methods**). Glutamatergic and GABAergic neurons formed two major groups of hypothalamic neurons in similar proportions across all vertebrates (**Fig. 1, D** and **E**). However, whereas glycinergic neurons are largely restricted to the medulla and spinal cord in mammals (*22*, *23*), we found a substantial number of glycinergic neurons in the *Xenopus* hypothalamus (**Fig. 1, D** and **E**). Moreover, unlike mammals where serotonergic neurons exclusively reside in the raphe nuclei in the brainstem (*24*, *25*), they were present in the hypothalamus of *Xenopus* and anole and exhibited molecular signatures distinct from mammalian raphe serotonergic neurons (**Fig. 1E**). Using STARmap-based spatial transcriptomic analysis (*26*), we determined the spatial distribution of neurotransmitter types in *Xenopus* and identified a large cluster of glycinergic neurons in the anterolateral hypothalamus, and some serotonergic neurons in the posteromedial hypothalamus (**fig. S10**).

### Mapping of hypothalamic subclasses across vertebrates

We mapped cell types onto the mouse reference using two algorithms built on distinct principles, reasoning that agreement between them would provide a cross-check over method-specific limitations (*27*). SAMap, a graph-based method, aligns single-cell transcriptomes across species through iteratively refined gene homology relationships and cellular manifolds (*28*). SATURN is a deep learning method that embeds cells from different species into a joint manifold using a protein language model to represent their genes (*29*), providing an orthogonal basis for cell type comparisons.

We used several additional strategies to increase mapping confidence (**fig. S9**). First, we mapped our data to mouse data down-sampled to 250 cells per subclass with resampling, keeping only the most consistent labels across resampling runs (**Methods**). Second, because voles diverged from mice only ∼20 million years ago and should therefore share most neuronal subclasses, we mapped vole cells to the mouse and then performed triangular mappings between each focal species, mouse, and vole (using either SAMap or SATURN), keeping only labels supported by independent mappings to the same subclass in both mammals. Third, because our sequencing coverage was lower in the vole than in the mouse atlas, we additionally retained mouse-only mappings from a focal species if labeling was supported by both SAMap and SATURN.

SAMap and SATURN mappings were largely concordant: ∼70% of sequenced cells per species had the same assignment (the same subclass, or unmapped by both), while < 2% received contradictory subclass assignments, which we discarded. The remaining cells were labeled by only one method (predominantly SAMap, > 90% of these cells). Of these cells, SATURN had the same mapping to mouse in most (∼50%) cases, leaving us with 6, 6, 10, and 4 subclasses in the quail, anole, *Xenopus*, and zebrafish that were only mapped by SAMap. We retained these SAMap-only mappings because they (i) consistently mapped to both the mouse and the vole, (ii) expressed cell type-specific TFs and neurotransmitter-related genes consistent with their mouse counterparts (see below), and (iii) mapped to one another across species.

Using the whole mouse brain atlas (*21*) as reference, our strategy mapped 68 of the 73 mouse neuronal subclasses in the hypothalamus (and related extended amygdala) in the vole, and 46, 42, 45, and 11 of them in the quail, anole, *Xenopus*, and zebrafish, respectively (**Fig. 2, A** to **C**). These data reveal consistent cell type diversity across tetrapods and a marked divergence in zebrafish. Around 10% of neurons in the vole, quail, anole, *Xenopus*, and zebrafish mapped to mouse subclasses bordering the hypothalamus (**fig. S11** to **S15**), likely because we dissected slightly beyond the hypothalamus (**Fig. 1B**) to avoid missing hypothalamic cell types. Another ∼5% of vole, *Xenopus*, and zebrafish neurons mapped to mouse subclasses far from the hypothalamus. The quail and anole fractions were higher (∼10%), with roughly half of those cells mapping to the optic tectum, which lies far from the hypothalamus in mammals yet closely adjacent to it in birds and reptiles (*30*) (**fig. S1** and **S2**). Although such non-hypothalamic mappings might reflect genuine shifts in cell type location (*31*, *32*), confirming these will require whole-brain single-cell and spatial transcriptomic data from non-mouse species. Nevertheless, the fact that most cells were mapped within or near the hypothalamus demonstrates the specificity of our approach. We therefore focused our subsequent analyses on the hypothalamic subclasses.

**Fig. 2.**
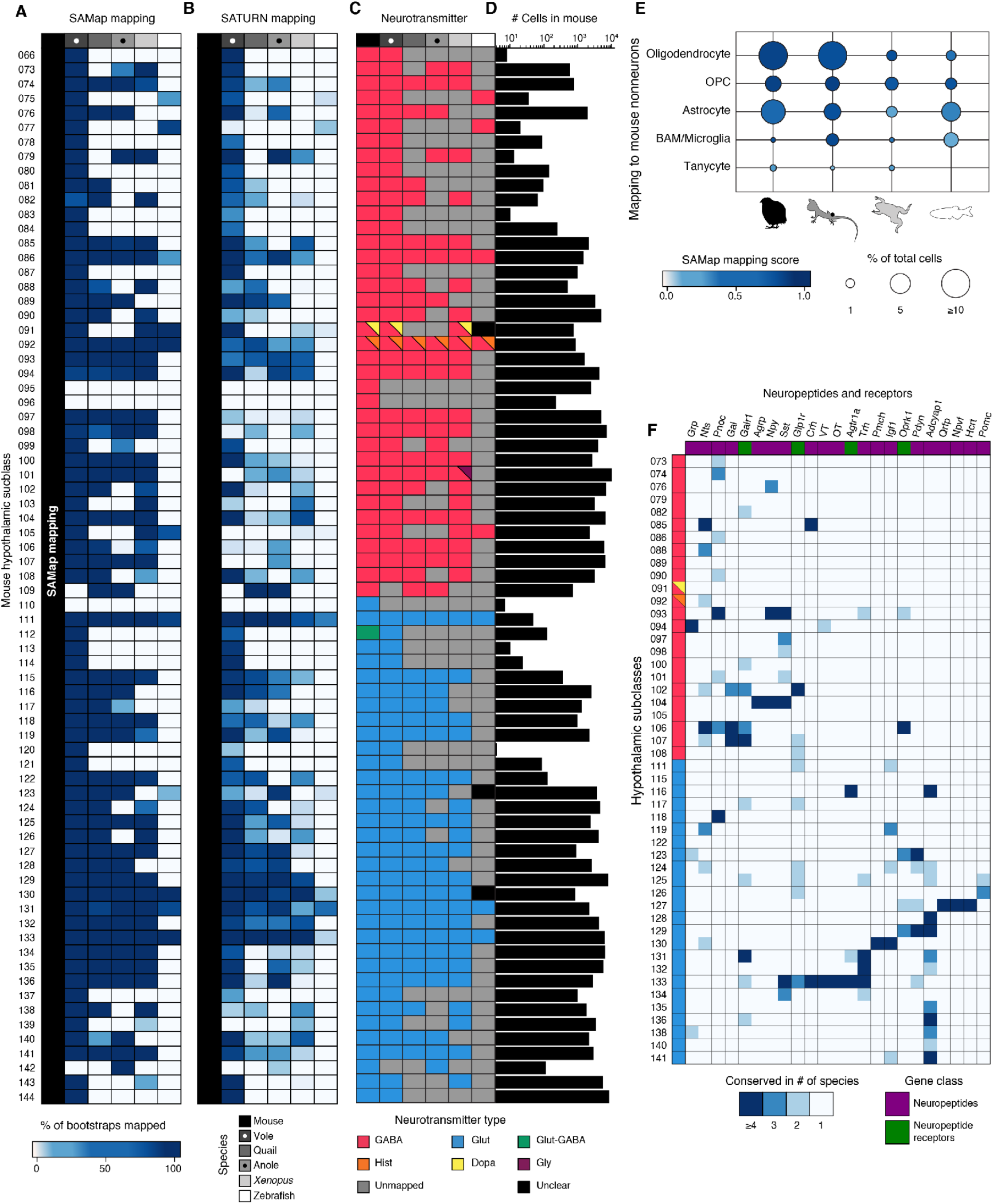
Mapping mouse hypothalamic subclasses across vertebrates. **(A** to **D)** Mapping of homologous cell types at subclass level. Each row represents one of the 73 hypothalamus subclasses in the mouse (*21*), with the subclass IDs on the left (full names provided in **table S1**). From left to right, heatmaps represent the fraction of resampling runs that mapped to the corresponding mouse subclass across 30 iterations of SAMap (A) or SATURN (B), mapped classes colored by their neurotransmitters (NT) type (C), and number of sequenced cells per subclass in the mouse hypothalamus (D). Gray in (C) represents mouse subclasses with no mapping in the corresponding species. **(E)** Dot plot comparing non-neuronal subclasses in the mouse with those in other species. OPC, oligodendrocyte precursor cell. Astrocytes and BAM/microglia each combined several subclasses (**fig. S16**) for simplicity. Vole data is not included, as non-neuronal cells were removed before sequencing (**Methods**). Color saturation indicates SAMap score and dot size indicates the percentage among all sequenced cells. **(F)** Conservation of enriched neuropeptides and four selected subclass-specific neuropeptide receptors. All genes are expressed in one subclass in at least four or more species except *Pomc*. Top, names of neuropeptides and receptors. Left, IDs for all hypothalamic subclasses mapped in four or more species. Color saturation shows the number of species where the gene is expressed in a mapped subclass. Expression patterns of all neuropeptides and receptors within each subclass across species are shown in **fig. S17** to **S28**.

In addition to neurons, we obtained a total of 33,588 high-quality non-neuronal nuclei from the quail, anole, and *Xenopus*, identified by the absence of canonical pan-neuronal markers. We mapped them, alongside non-neuronal cells in the zebrafish to the 23 non-neuronal subclasses in the mouse and found homology to 21 of the 23 in at least one species (**fig. S16**). These included 7 groups with homology across all four species— oligodendrocytes, oligodendrocyte precursor cells (OPCs), astrocytes, macrophages, arachnoid barrier cells, vascular leptomeningeal cells, and endothelial cells—the last 3 of which were rare vascular subclasses (∼0.1% of cells). Of the remaining 4 groups (**Fig. 2E**), oligodendrocytes and OPCs mapped with high scores in all species, but the proportion of oligodendrocytes was much higher in amniotes (reptiles, birds, and mammals) than in anamniotes (fish and amphibians), suggesting that amniotes have more myelinated axons in the hypothalamus. Consistent OPC abundance across species, despite large variation in oligodendrocyte numbers, supports the notion that OPCs have functions beyond producing oligodendrocytes (*33*). Both oligodendrocytes and OPCs had high mapping scores across species. By contrast, mammalian-typical astrocytes had lower mapping scores in *Xenopus* and zebrafish (**fig. S16**), suggesting further divergence of these subclasses from those in mammals. Macrophage-like cells could also be found in all species; however, microglia could not be resolved from border-associated macrophages (BAMs) in most species, as both are macrophages with overlapping transcriptomic signatures. Finally, tanycytes, specialized glial cells that line the hypothalamic ventricle and are responsible for sensing and responding to metabolic signals, were found in quail, anole, and *Xenopus*, but not in zebrafish (**Fig. 2E**).

### Conservation of neurotransmitter and neuropeptide expression

Small-molecule neurotransmitter usage was highly stable in mapped cell types across species (**Fig. 2C**). Exceptions included a GABAergic subclass (101) with a subset of cells also expressing glycinergic markers in *Xenopus*, and a mammalian-specific subclass (112) that was glutamate-GABA double positive in mouse, but only glutamatergic in the vole.

In addition to small-molecule transmitters, the hypothalamus expresses a diversity of neuropeptides with distinct physiological functions (*34*). We analyzed the conservation of neuropeptide expression along with their cognate receptors (*35*) (**fig. S17** to **S28**). We found 18 neuropeptides consistently enriched in at least one mapped subclass by four or more species (**Fig. 2F**). For example, arginine vasopressin (also called vasotocin, *VT*), oxytocin (*OT*), corticotropin-releasing hormone (*Crh*), and thyrotropin-releasing hormone (*Trh*) expression was highly conserved in subclass 133 across species, which in the mouse resides in the PVH. With the exception of *Trh* (*36*), these peptides and their homologs play highly conserved roles in regulating water balance, maternal and social behavior, stress, and metabolism from mammals to teleosts (*37*–*54*). Additionally, *Agrp*—encoding agouti-related peptide that acts as an appetite-stimulating neuropeptide in all tetropods (*55*, *56*)—shows conserved expression in subclass 104. *Pomc—*encoding pro-opiomelanocortin, a neuropeptide best known for inhibiting appetite and signaling stress (*55*, *57*, *58*)—was conserved in three species in subclass 126. Both *Agrp* and *Pomc* neurons in the mouse reside in the arcuate nucleus. *Gal—*encoding the neuropeptide galanin implicated across taxa in parental behavior (*59*–*61*) and feeding (*62*–*65*), as well as its receptor, were conserved in subclass 106 and 107, which in the mouse reside respectively in the medial preoptic area (MPOA) and dorsomedial hypothalamus (DMH). *Hcrt*—encoding hypocretin (orexin) and promoting wakefulness and feeding across vertebrates (*66*–*69*)—was conserved in subclass 127, which in the mouse resides in the DMH and lateral hypothalamus (LH). *Grp* (gastrin-releasing peptide)—regulating circadian rhythms *(70)*—was conserved across amniotes in subclass 94, which in the mouse resides in the suprachiasmatic nucleus (SCN). By contrast, two other neuropeptides that regulate circadian rhythms—vasoactive intestinal polypeptide (*Vip)* and prokineticin 2 (*Prok2*) (*71*, *72*)—were only enriched in subclass 94 in mammals.

Overall, neuropeptide receptors were more broadly expressed across the hypothalamus than neuropeptides (**Fig. 2F; fig. S17** to **S28**). One of the notable exceptions was *Agtr1,* encoding angiotensin II receptor type 1 involved in regulation of water balance (*73*–*77*). *Agtr1* was enriched in a few subclasses, including 116 in the mouse lamina terminalis and 133 in the mouse PVH.

### Evolution of gene regulatory network

TFs are key regulators of cell type identity (*78*, *79*). Indeed, each conserved subclass contained a distinct set of enriched TFs, and these combinations were largely conserved across species (**Fig. 3A**; **fig. S29** to **S34**). Nearly half of these conserved TFs were homeodomain TFs (**Fig. 3B**), which delineate neuronal cell types in *C. elegans* (*80*). However, relative to its family size, homeodomain TFs were not particularly conserved compared to other TF families.

**Fig. 3.**
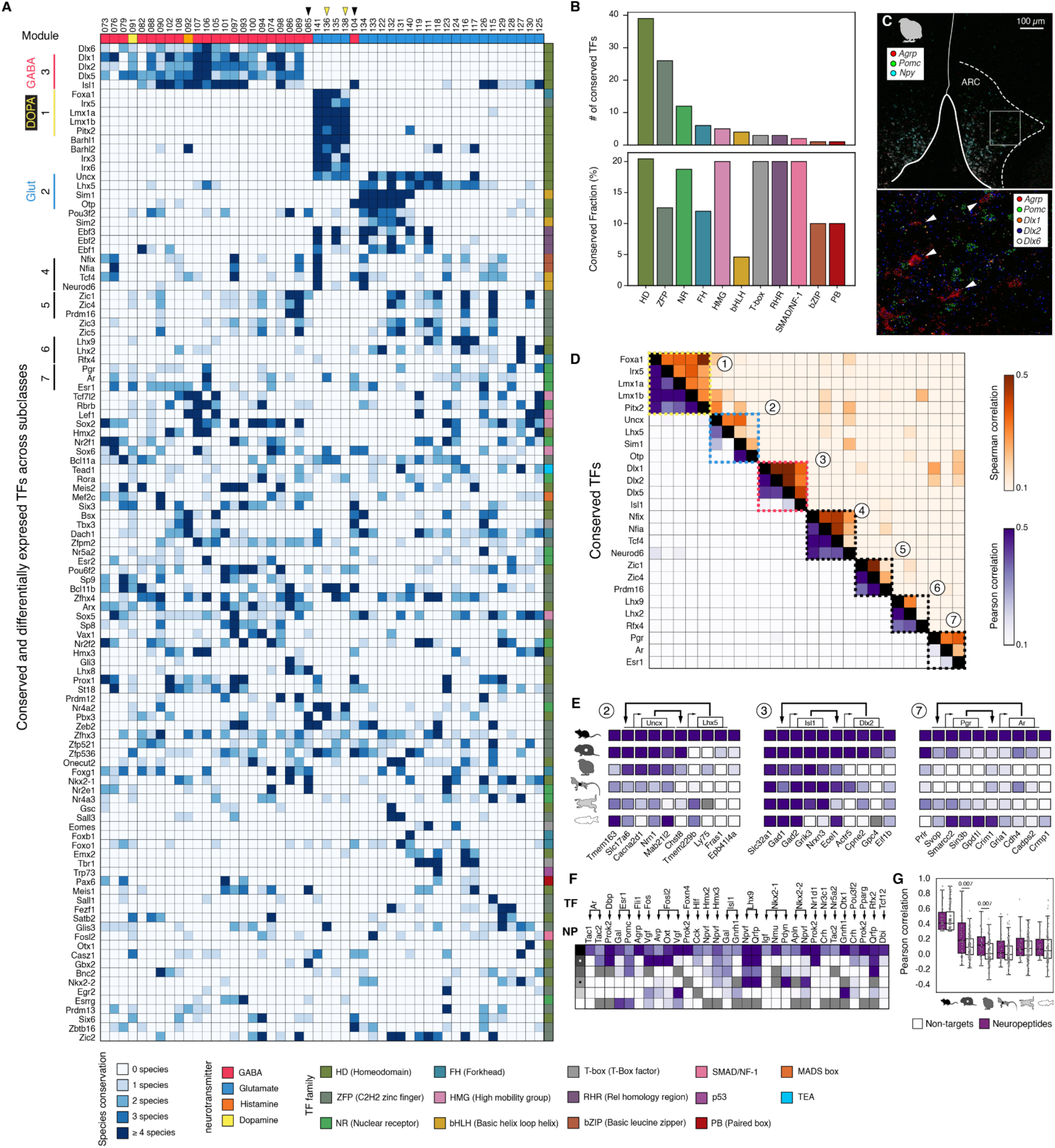
Evolution of transcription factor expression in hypothalamic subclasses across vertebrates. **(A)** Conservation of differentially expressed transcription factors across species. Top row, IDs for the mouse hypothalamic subclasses found in at least four species. Second row, neurotransmitter type of the corresponding hypothalamic subclass. Inverted black triangles indicate inhibitory neuron subclasses where Dlx are not conserved. Inverted yellow triangles indicate excitatory neuron subclasses that contain sub-populations of dopaminergic neurons. The heatmap shows the TFs shared by at least four species in any subclass except Pgr. Color indicates the number of species in which the TF is enriched in that subclass. Right column, TF family as defined by the JASPAR database (**Methods**; color index below). Abbreviation of TF classes, a full list of genes and subclasses, and the expression of each TF within each subclass can be found in **fig. S2G** to **S34**. **(B)** Top, total number of TFs from major TF families that had conserved expression in at least one cell type in at least four species. Bottom, percentage of the TFs in the family with conserved expression. **(C)** STARmap images showing *Dlx* expression in the quail arcuate nucleus. Arrows point at *Agrp*-expressing cells corresponding to subclass 104. Scale bar, 100 µm. **(D)** The next-to-worst correlation within conserved TF modules. We used the next-to-worst rather than the worst value to account for variation in gene annotation. Bottom triangle: Pearson correlation between genes; top triangle: Spearman correlation between genes. **(E)** Cross-species correlation for three conserved co-regulatory TF pairs and downstream genes (boxes are gray where the gene has no ortholog in the corresponding species), identified using SCENIC+ (**Methods)**. Plotted genes are the ten most correlated non-TF genes in the mouse that are regulated by at least one TF in the module above an area-under-the-curve (AUC) threshold according to SCENIC+ (unordered). Labels (2, 3, 7) correspond to module numbers in (D). See additional examples in **fig. S35**. **(F)** Correlations between TFs and downstream neuropeptides (NP) across species. Regulatory relationships between TFs and neuropeptides were identified using SCENIC+. **(G)** Comparison of the Pearson correlation between each TF and target neuropeptides identified in (F) (purple) and between each TF and 5 non-targets within the same distribution (white). Statistical analysis was performed using a Mann-Whitney U test. P-values < 0.05 are labeled.

Co-expression of TFs can reveal gene regulatory logic, so we next asked whether such relationships are conserved. Defining a TF pair as co-expressed when its Pearson correlation across hypothalamic subclasses exceeded a modest threshold (e.g., 0.3), we identified 43,625 pairs co-expressed in at least one species. However, only 267 pairs (∼0.6%) were conserved across ≥ 5 species, no more than expected by chance (346 pairs, 95% confidence interval: 245 – 456; see **Methods** for estimation accounting for dropout). The low fraction of conserved co-expressed TFs held true across correlation thresholds and metrics (**table S2**). Thus, even where cell type identity was conserved, TF–TF co-expression patterns were evolutionarily labile.

We next grouped co-expressed TFs into modules*—*sets of TFs that exhibit mutual correlation across individual cells and likely act together*—*and found several conserved modules that tracked small-molecule neurotransmitter expression. The *Foxa-Lmx1* module, associated with dopaminergic fate (*81*) (module 1, **Fig. 3 A** and **D**), was found in several glutamatergic subclasses, including those (136 and 138) containing dopaminergic neurons. The *Uncx*-*Sim1-Otp* module (module 2) was expressed in subsets of glutamatergic neurons. The *Dlx* module was expressed by most GABAergic neurons (module 3), consistent with the observations that inhibitory neuronal development requires *Dlx* TFs from fish to mammals (*82*, *83*). One notable exception was subclass 104, which contains the feeding-promoting *Agrp* neurons in the arcuate nucleus. STARmap confirmed that *Agrp* and *Dlx* were expressed in largely non-overlapping cells in the quail arcuate nucleus (**Fig. 3C**). These findings suggest that tetrapod *Agrp* neurons have a developmental and evolutionary history distinct from other GABAergic neurons. Indeed, a substantial fraction of mouse *Agrp* neurons originate from the same lineage that produces glutamatergic *Pomc* neurons, which signal satiety and thus have opposing function of *Agrp* neurons (*84*). Additional modules included neural differentiation factors including *Neurod6* (module 4), known to act together in cortical development (*85*–*87*), expressed in subclass 141 (located in mouse supramammillary nucleus); the *Prdm-Zic* module (module 5) in subclasses 89 (broadly distributed in mouse hypothalamus) and 115 (concentrated in mouse septum); the *Lhx-Rfx* module in subclass 127 of the mouse DMH (module 6); and the nuclear hormone receptor Pgr-Ar-Esr1 module in subclass 106 in inhibitory neurons of the mouse preoptic area involved in social behavior (*88*–*91*) (module 7).

It has been assumed that conserved TF modules provide evolutionary stability for their downstream effectors, with some TFs also regulating each other, forming feedback loops that promote evolutionary stability (*79*, *92*). To test these assumptions, we leveraged SCENIC+ (*93*) and a publicly available ATAC-seq dataset from the mouse brain (*94*) to reconstruct the mouse hypothalamic gene regulatory network, integrating both expression and *cis*-regulatory information. Among the 142 co-regulatory TF pairs identified by SCENIC+, 16 comprised conserved co-expressed TFs (**Fig. 3E; fig. S35**). The percentage of coregulatory TFs that were conserved (∼10%) was significantly higher than the percentage of coexpressed TFs (∼0.6%). However, the downstream effectors of these co-regulatory TFs underwent drastic turnover: their mouse targets often lost expression correlation even in the closely related vole. Although a few specific TF–effector relationships persisted across vertebrates—such as genes involved in GABA transmission with the *Dlx* module and *Slc17a6* (encoding vesicular glutamate transporter 2) with the *Uncx* module—these were exceptions rather than the rule (**Fig. 3E, fig. S35**).

Given the central role of neuropeptides in hypothalamic function and our finding that their expression is conserved in many subclasses (**Fig. 2F**), we next asked whether regulatory relationships between TFs and neuropeptides are preferentially conserved. For each TF predicted to regulate at least one neuropeptide in the mouse, we identified non-target genes whose correlation with that TF most closely matched its TF–neuropeptide correlation in the mouse, then compared cross-species conservation of the TF–neuropeptide pairs against these matched non-target pairs. We found that TF–neuropeptide relationships were not preferentially conserved in the anole, *Xenopus*, and zebrafish (**Fig. 3, F** and **G; fig S35**), suggesting that the TF–neuropeptide relationship also erodes over longer evolutionary distances.

### Conserved gene expression, cell type composition, and function in the PVH across vertebrates

Among hypothalamic neurons, 6 subclasses were conserved across all vertebrate species we analyzed (**Fig. 2, A** and **B)**, most of which have well-characterized functions in the mouse (**Table 1; fig. S36**). To examine how deeply hypothalamic cell types are conserved at the level of functionally distinct supertypes (a finer transcriptomic classification than the subclasses thus far examined), we focused on subclass 133, both because its functions are well described in the mouse and because it had the highest average mapping score across both SAMap and SATURN (**Fig. 2, A** and **B**). Subclass 133 contains neurons expressing neuropeptides VT, OT, Crh, Trh, and Sst that have distinct functions as discussed earlier; these neuropeptides also serve as markers for the supertypes in the mouse.

**Table 1.** Conserved subclasses across vertebrates and their potential functions.

| Subclass | NT<sup>1</sup> | Location<sup>2</sup> | Representative marker genes<sup>3,4</sup> | Potential functions<sup>5</sup> |
| --- | --- | --- | --- | --- |
| 086 | GABA | MPOA | *Lhx8*<sup>3</sup>, *NmU*<sup>4</sup> | Regulate anxiety-like behavior |
| 092 | GABA, His | TMN | *Hdc* | Promote wakefulness; suppress food intake |
| 111 | Glut | septum | *Tbr1*<sup>3</sup>, *Sln* | Promote social memory |
| 130 | Glut | LHA | *Otx13*<sup>3</sup>, *Pmch*<sup>4</sup> | Regulation of feeding, mood, sleep-wake cycle and energy balance |
| 131 | Glut | LHA | *Trh*<sup>4</sup> | Closely related; 133 constitutes most PVH cells; 131 is lateral to PVH. Regulate social and parental behavior, water balance, stress response, metabolism, and growth. |
| 133 | Glut | PVH, SON | *VT*<sup>4</sup>, *OT*<sup>4</sup>, *Crh*<sup>4</sup>, *Trh*<sup>4</sup>, *Sst*<sup>4</sup> |  |

<sup>1</sup> NT, neurotransmitter. His, histamine; Glut, glutamate; GABA, gamma-aminobutyric acid.
<sup>2</sup> Listed are primary location(s). MPOA, medial preoptic area; TMN, tuberomammillary nucleus; AHN, anterior hypothalamic nucleus; ZI, zona incerta; ARC, arcuate nucleus; DMH, dorsomedial hypothalamic nucleus; VMH, ventromedial hypothalamic nucleus; LH, lateral hypothalamic area; PVH, paraventricular hypothalamic nucleus; SON, supraoptic nucleus.
<sup>3</sup> Marker genes that encode transcription factors.
<sup>4</sup> Marker genes that encode neuropeptides. See table S3 for additional marker genes.
<sup>5</sup> A partial list based mostly on studies in the mouse. See table S3 for references.

We annotated the Allen Institute supertypes within subclass 133 based on neuropeptide and TF expression and mapped them across species using SAMap (**Fig. 4, A** and **B**). Most mapped supertypes expressed the same neuropeptide (**Fig. 4B**), although their abundance sometimes varied across species (pie charts in **Fig. 4, C** to **E**). For example, mice had an expanded *Crh*^+^ population, *Xenopus* had an expanded *Nkx2-2*^+^ population and a reduced *OT*^+^ population, and quail had a larger proportion of *VT*^+^ cells but few cells expressing *Crh* (**fig. S17** to **S22**).

**Fig. 4.**
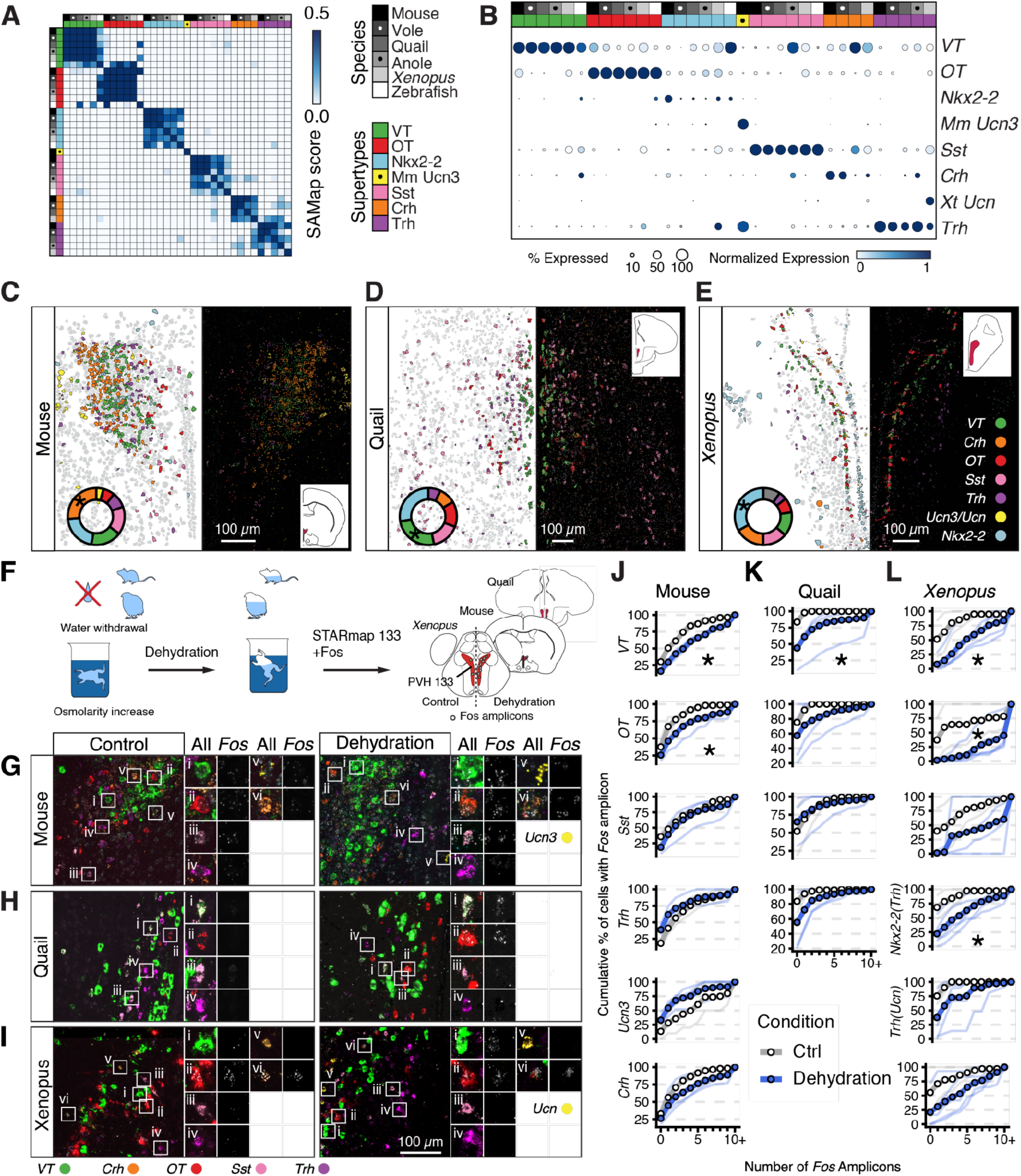
Conservation of PVH supertypes across species. **(A)** Heatmap showing the SAMap score among PVH supertypes (blue color index). Indices for species and supertypes are on the right and shared with (B). **(B)** Expression profiles of marker genes in the PVH across species. Mouse Mm_Ucn3 supertype is unique to mouse. Note that expression of some neuropeptides is not exclusive to a single supertype. **(C** to **E)** Left panels, representative STARmap images of PVH from mouse (C), Japanese quail (D), and *Xenopus* (E), showing amplicons from selected marker genes. Right panels, mirror of the left panels (across the midline) summarizing the supertypes identified in STARmap. Color codes are the same as (A), except that purple indicates *Trh*+ *Nkx2-*2 supertype, and yellow indicates *Ucn*+ *Trh* supertype in the *Xenopus*. Pie charts represent the proportion of cells in each supertype in the snRNA-seq data; asterisks indicate the expanded population. The grey color in the *Xenopus* pie charts indicates unmapped cells. Highlighted regions in the insets indicate where the images were taken. **(F)** Schematic illustration of the dehydration experiment. The last panel also indicates the proposed new location of the frog PVH based on the results in (E). Schematics of coronal brain sections are not to scale. **(G** to **I)** Example STARmap images showing *Fos* and supertype marker gene expression patterns in PVH sections of control and dehydrated mice (G), quail (H), and *Xenopus* (I). Dots represent amplicons. Insets, representative cells expressing supertype markers and *Fos*. **(J** to **L)** Quantification of *Fos* expression in (G to I) with cumulative plots. In the *Xenopus*, cells expressing *Ucn* belong to *Trh* supertype. Cells expressing *Trh* alone are a subset of *Nkx2-2* supertype. Thin lines represent individual sections (1 or 2 sections per animal). Thick lines with circles represent averages (*n* = 2 animals each for controls in the mouse, quail, and *Xenopus*; n = 3, 3, 4 animals for dehydration in the mouse, quail, and Xenopus, respectively). Asterisks denote statistically significant differences (*p < 0.05*) between the dehydration and control conditions, as determined by a generalized linear mixed-effects model (GLMM). A right shift of the cumulative plot means increased Fos signals.

We leveraged the expression of neuropeptides *VT*, *OT*, *Crh*, *Trh*, and *Sst* alongside conserved TFs to capture the spatial distribution of these supertypes in our representative bird and amphibian species using STARmap (**Fig. 4C** to **E**). Not only were these neuropeptides spatially clustered in quail and *Xenopus*, but also many conserved TFs including *Sim1*, *Otp*, and *Ebf3* were co-expressed in these cells (**fig. S37**). These data provide compelling evidence that the supertypes we identified in quail and *Xenopus* are homologous to those in the PVH of the mouse. This finding also prompted a revision of hypothalamic anatomy for *Xenopus*, delineating a clear location for the PVH (**fig. S10)**.

We next tested whether homologous cell types also share physiological responses. *VT*^+^ and *OT^+^* PVH neurons are known to be activated by dehydration in the mouse (*95*). We tested whether dehydration activates homologous supertypes in *Xenopus* and quail using the expression of immediate early genes (IEGs) *Fos* and *Egr1* to identify activated neurons (**Fig. 4, F** to **I**; **fig. S37**). We found that in both *Xenopus* and quail, dehydration activated IEG expression in the presumptive PVH neurons, preferentially in *VT* and *OT* supertypes, similar to that in mice (**Fig. 4, G** to **L; fig. S37**). These observations suggest that this subclass not only shares molecular identity but also functional properties even at finer resolutions.

### Diversification of cell types in the vertebrate hypothalamus

Having established conservation of cell types across the vertebrate PVH at both the transcriptomic and functional levels, we next asked where diversification has occurred by examining clade-specific subclasses. Thirteen mouse subclasses were not detected outside of the mammals (**Fig. 2, A** and **B**), and another three were divergent between mouse and vole. However, 10 of these 16 subclasses were exceedingly rare (< 0.1% of neurons in the mouse hypothalamus, **Fig. 2D**) and may be undetected in non-mammalian species due to lower sequencing coverage. The six remaining subclasses (**Fig. 5, A** to **C**; top rows) spanned the anterior-posterior axis of the hypothalamus (**Fig. 5D**). Most were spatially intermingled with other subclasses that mapped beyond mammals. An exception was subclass 144, which accounts for most excitatory neurons in the mammillary nuclei, a hypothalamic structure that has expanded in mammals and is involved in spatial memory (*19*, *96*).

**Fig. 5.**
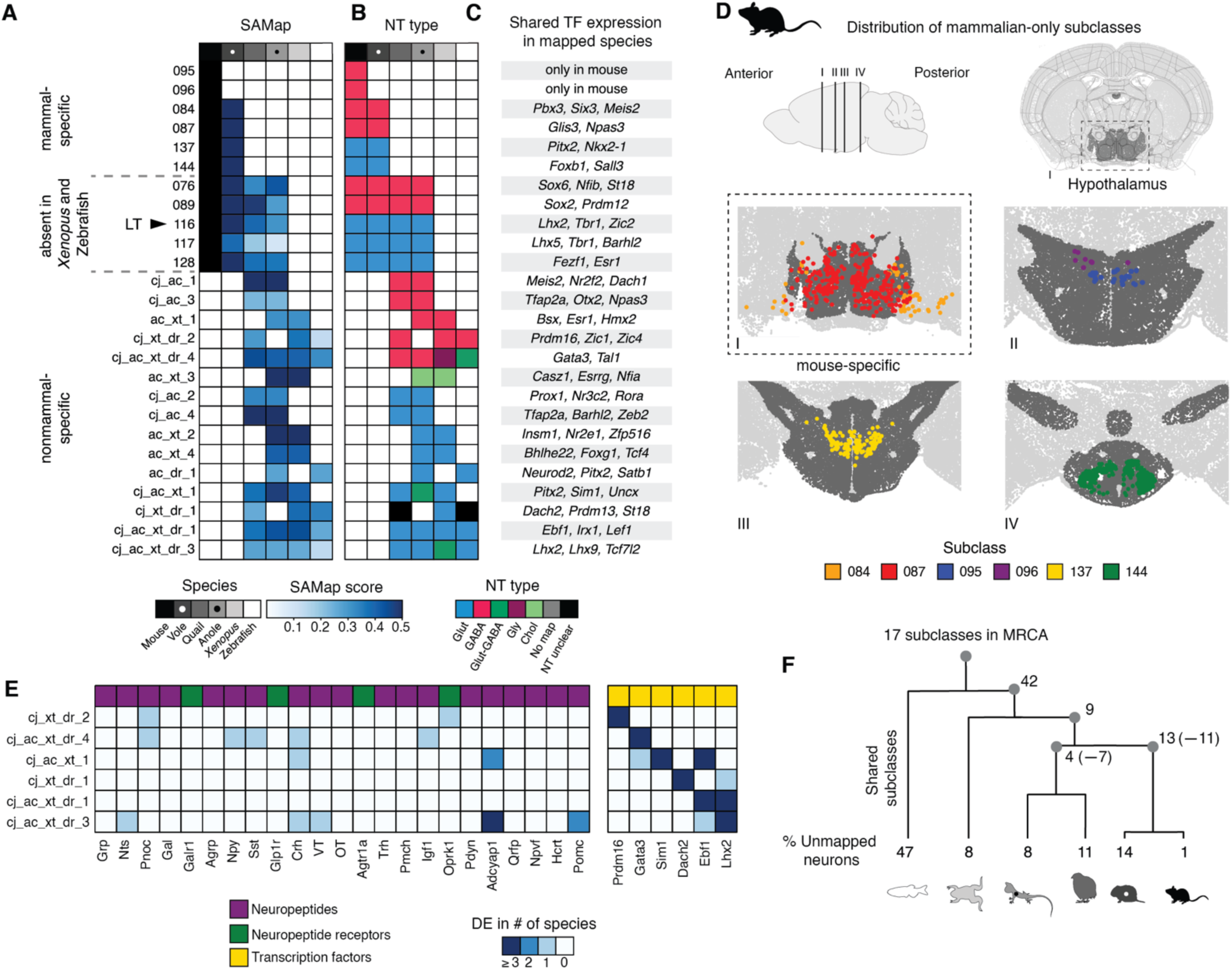
Analysis of clade-specific cell types. **(A)** Mapping of clade-specific subclasses. For subclasses with mouse mappings, boxes are colored by their SAMap score to the mouse subclass; for non-mammalian subclasses, boxes are colored by the average SAMap score across mappings to all other species in which the subclass is found. **(B)** Neurotransmitter types of the subclasses in (A). **(C)** List of differentially expressed transcription factors (TFs) conserved in each plotted subclass across all mapped species. **(D)** Localization of mammal-specific subclasses based on Allen Institute whole-mouse-brain MERFISH data (*21*). **(E)** Conservation of enriched neuropeptides (from **Fig. 2F**) and transcription factors across species. Top row, category of gene. **(F)** Summary of clade-specific subclasses superimposed on the evolutionary distance tree. Numbers at the gray nodes indicate the subclasses that are added or subtracted (–) to the most recent common ancestor (MRCA) of each clade (**Methods**). Numbers at the bottom indicate the percentage of unmapped neurons in each species.

Through pairwise mappings of cells that did not map to any mouse cell types (including non-hypothalamic types) between non-mammalian species (**Methods**), we identified 15 non-mammalian subclasses that were conserved between at least two species, which we named accordingly; for example, *cj_ac_1* denotes a subclass shared only between quail and anole (**Fig. 5, A** to **C**, bottom rows). Similar to the subclasses shared with mammals, the neurotransmitter types were largely conserved with a few exceptions (e.g., glycinergic markers found in *Xenopus cj_ac_xt_dr_4*).

To further characterize these non-mammalian subclasses, we examined neuropeptide expression. Surprisingly, most neuropeptides were not conserved across species within these non-mammalian subclasses (**Fig. 5E**, left): only *Pomc* and *Adcyap1* were partially conserved in specific subclasses. This contrasts both with mammalian mapped subclasses, many (42 of the 48 mammalian subclasses found in at least 4 species) of which had at least one conserved neuropeptide (**Fig. 2F**), and with the TFs of the same subclasses, which remained highly conserved (**Fig. 5A**; **Fig. 5E**, right). This high level of divergence extended to other effectors as well (**fig. S38**). One interpretation is that these non-mammalian subclasses tolerate greater turnover in their effector genes than highly conserved subclasses such as the PVH; further studies are needed to determine whether this corresponds to greater functional flexibility.

Each species in our dataset also contained species-specific, unmapped cell clusters (**Fig. 5F**). Notably, zebrafish had more than 4 times as many species-specific neurons as any tetrapod, suggesting that much of the hypothalamic cell type repertoire in the teleost lineage has diverged transcriptomically beyond our ability to map. Identifying the functions of these clade-specific cell types would be a valuable next step toward understanding how these cell types contribute to diversification of physiology and behaviors among vertebrates.

### Cell type diversification in the lamina terminalis across terrestrial and aquatic vertebrates

To gain deeper insights into the diversification of cell types, we next examined subclass 116, which appeared to be amniote-specific (**Fig. 5A**, middle). In the mouse, subclass 116 encompasses all glutamatergic neurons of the lamina terminalis (LT), which comprises the subfornical organ (SFO), organ of vascular lamina terminalis (OVLT), and the median preoptic nucleus (MnPO). The SFO and OVLT in mammals have fenestrated capillaries allowing local neurons to directly access the bloodstream and sense osmolarity and hormones that signal dehydration (*6*). Interestingly, subclass 116 was only found in terrestrial amniotes we sampled, but not in *Xenopus* or zebrafish. Given LT’s function as an osmometer was proposed to first appear in tetrapods (*97*, *98*) (**Fig. 6A**), but *Xenopus* is an amphibian that returned to being primarily aquatic (*99*), we asked whether the presence of subclass 116 coincides with terrestriality. Indeed, analyzing the snRNA-seq data of the hypothalamus from a terrestrial frog, the mimic poison frog (*Ranitomeya imitator*) (**Methods**), using the same mapping approaches (**fig. S9**), revealed that subclass 116 was present in this frog (**Fig. 6B**). These data suggest that subclass 116 was present in the last common ancestor of amniotes and amphibians but was subsequently lost (or substantially diverged beyond our ability to map) in *Xenopus*.

**Fig. 6.**
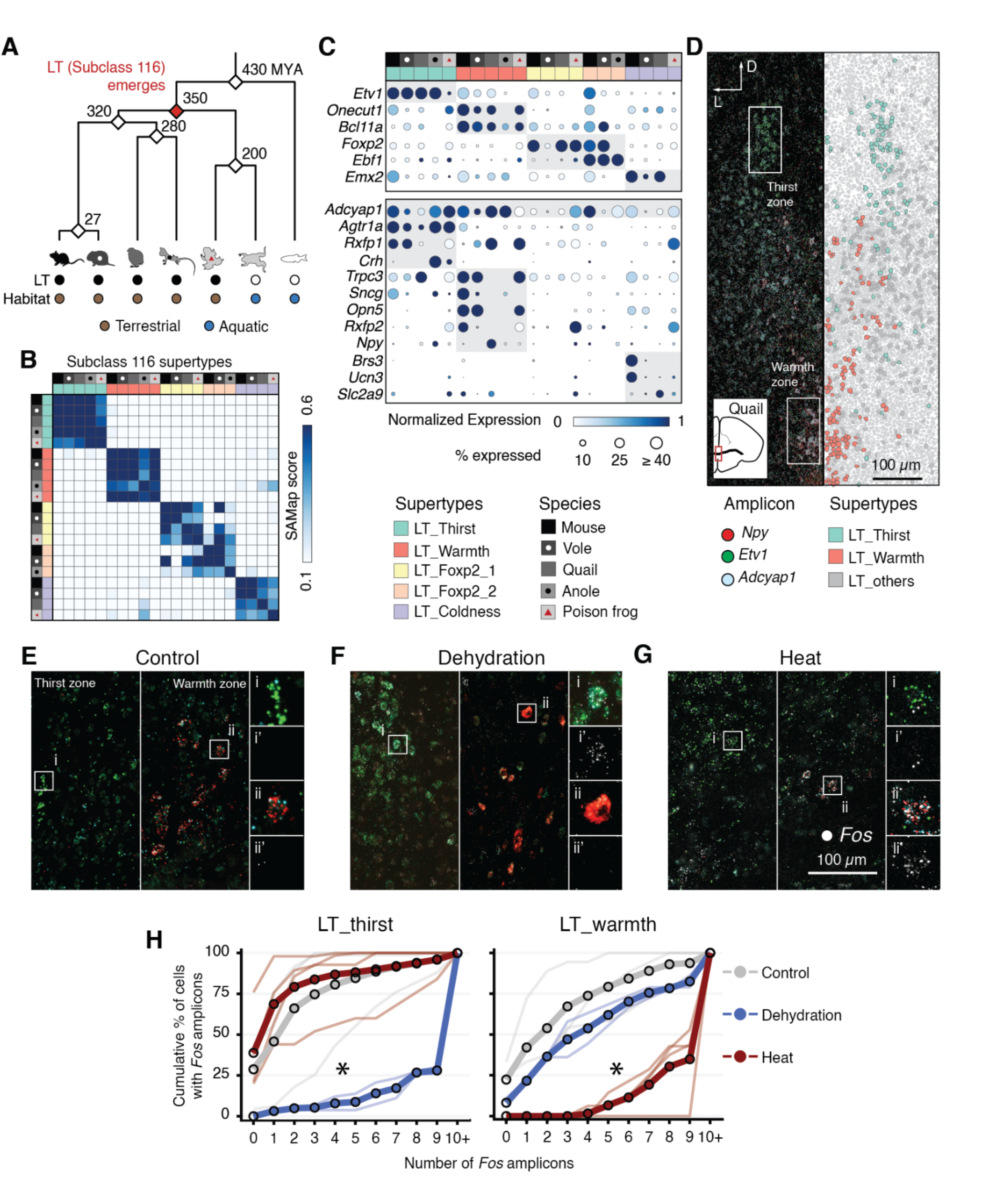
Analysis of lamina terminalis (LT) cell types. **(A)** Schematic of the proposed evolutionary emergence of the LT. **(B)** Mapping of supertypes within subclass 116. The color scale represents the SAMap score. The first row and column indicate the species. The second row and column indicate the supertypes. **(C)** Dot plots showing the expression of marker genes: TFs (upper panel) and other markers (lower panel). Color code is the same as in (B). **(D)** Representative STARmap image of the LT in the quail. The left panel shows amplicons from marker genes *Npy* (red), *Etv1* (green), *Adcyap1* (cyan); the right panel is a mirrored (across the midline) annotation of the left panel for thirst and warmth supertypes. The inset on the left bottom shows the outline of the section from which the image was taken. Rectangles indicate the approximate LT regions where the left and middle panels in (E to G) are from. **(E** to **G)** Representative STARmap images of LT with marker genes *Npy* (red) and *Etv1* (green), *Adcyap1* (cyan) and *Fos* (white) from quails under the control (E), dehydration (F), and heat (G) stimulation conditions. The left panels are from the thirst zone; the middle panels are from the warmth zone; the right panels show high magnification of the boxed regions, with the lower panel of each pair showing only *Fos* amplicons. Note that variations in the images are due to variations in section planes. **(H)** Quantification of *Fos* expression in (E to G) with cumulative plots. Thin lines represent individual sections (1 or 2 per animal). Thick lines with circles represent averages (*n* = 3, 2, 3 animals for control, dehydration, and heat conditions, respectively). Asterisks denote statistically significant differences (*p* < 0.05) between dehydration and control (left), or heat and control (right), as determined by a generalized linear mixed-effects model (GLMM). A right shift of the cumulative plot means increased Fos signals.

In the mouse, subclass 116 is marked by the expression of *Adcyap1* (encoding pituitary adenylate cyclase-activating polypeptide, PACAP) and contains five transcriptomically distinct supertypes (**Fig. 6, B** and **C**). These include *Agtr1a^+^* SFO, MnPO, and OVLT neurons that are activated by dehydration (*6*, *100*, *101*) (thirst neurons hereafter); gamma synuclein–encoding *Sncg^+^* (*102*) and UV opsin–encoding *Opn5^+^* (*103*) neurons in the MnPO that are activated by increases in environmental temperature (warmth neurons hereafter); *Ucn3* (encoding neuropeptide urocortin-3)*-* and *Brs3* (encoding Bombesin-like neuropeptide receptor)*-*expressing MnPO neurons, which are involved in infanticide (*104*) and respond to decreases in environmental temperature (*105*) (coldness neurons hereafter); and two additional supertypes expressing the transcription factor *Foxp2*. We used these labels to annotate supertypes across amniotes within subclass 116 and examined their evolutionary relationships using SAMap (**Fig. 6B**).

Supertypes that include thirst, warmth, and coldness neurons were present in every species that had subclass 116 except for a lack of *Brs3^+^*coldness neurons in the anole. The mapped supertypes expressed conserved TFs across species (**Fig. 6C**, upper panel), including *Etv1* in the thirst neurons and *Bcl11a* in the warmth neurons. However, neuropeptides, their receptors, and other effectors were highly divergent. Most mouse markers, such as *Sncg*, *Rxfp2* (encoding a neuropeptide receptor), *Ucn3*, and *Brs3* were no longer markers in other species (**Fig. 6C**, bottom panel). *Trpc3* (encoding a diacylglycerol-activated Trp channel) and *Opn5* were still markers in the vole supertypes that mapped to mouse warmth neurons, were lost in the homologous supertypes in quail and anole, yet were retained in the poison frog. *Rxfp1* (encoding a receptor for the neuropeptide relaxin) was a marker in thirst neurons in mice but was expressed in the quail and poison frog supertypes that mapped to mouse warmth neurons. Conversely, markers for quail supertypes mapped to the mouse warmth and coldness neurons—*Npy* (encoding neuropeptide Y) and *Slc2a9* (encoding a uric acid transporter), respectively—were unique to the quail. Likewise, *Crh,* a marker for the anole and poison frog supertypes mapped to the mouse thirst neurons was absent in other species (**Fig. 6C** bottom panel). STARmap experiments showed clustered but segregated expression of several markers near the LT of the quail (**Fig. 6D**), validating supertype composition and their LT identity for quail subclass 116.

Given the conservation of transcriptomically defined supertypes within subclass 116 and the striking divergence in effector gene expression, we asked whether homologous supertypes still perform similar physiological functions such as sensing dehydration and heat. We subjected quail to dehydration or heat and measured *Fos* and *Egr1* expression with STARmap. We found that dehydration and heat activated the quail supertypes that were transcriptomically homologous to the mouse thirst and warmth supertypes, respectively (**Fig. 6, E** to **H**; **fig. S39**). Dehydration-activated supertype in the quail expressed *Agtr1* as in the mouse (**Fig. 6F**), suggesting that birds and mammals may sense dehydration through shared angiotensin-related mechanisms (*106*). Despite divergence in marker gene expression in warmth neurons, our data suggest that mammals and birds nevertheless use homologous supertypes to sense heat, an essential component of homeostatic regulation of body temperature.

## Discussion

We generated hypothalamic snRNA-seq atlases across four vertebrate classes and spatial transcriptomes for birds and amphibians, establishing resources for tracing the evolution of hypothalamic cell types that regulate essential bodily functions and innate behavior. Our comparative transcriptomic analyses reveal that hypothalamic neuronal and glial cell types are diverse in all vertebrate lineages: glutamatergic and GABAergic neuronal types are the most prevalent, in roughly equal proportion, across vertebrates; some cell types are conserved across all vertebrates whereas others are clade-specific; and many TFs and neuropeptides have conserved expression patterns in homologous cell types.

Among those cell types conserved across vertebrates, most carry essential, well-characterized physiological functions (**Table 1**)—PVH neurons that regulate water balance, DMH neurons that sense coldness, and tuberomammillary nucleus histaminergic neurons that promote wakefulness. We propose that these represent ‘keystone’ cell types: evolutionarily ancient populations whose functions are so fundamental that they have been maintained across all vertebrate lineages, and around which more specialized, clade-specific types may have been elaborated.

Another central implication of our findings is that the transcriptomic identity and function of a cell type can be conserved even when the marker genes traditionally used to define functional subtypes are not. This was supported by our analysis of regulatory architecture, in which TF codes were stable but the regulatory linkages between TFs and their downstream targets were labile. This pattern is reminiscent of the extensive evolutionary turnover of *cis*-regulatory elements documented for developmental enhancers (*107*, *108*) and extends this principle to the specification of neuronal cell types (*109*). However, we also observed neurotransmitter stability across vertebrate evolution, supporting the idea that neurotransmitter identity is specified by master regulators that control suites of genes for entire neurotransmitter systems (*110*–*112*). Such partitioned stability and flexibility allow evolution to alter how a cell type executes its function without dismantling the TF code and its primary communication channel that specifies the cell type itself.

At the functional level, the deep conservation of neural responses to dehydration in PVH is remarkable given the diverse ways animals meet the challenge of water homeostasis. The same neurons and endocrine signals (i.e., VT) respond to dehydration, yet the peripheral structures involved differ, from the kidneys in mammals to salt glands in birds and reptiles, and the skin in amphibians (*7*).

Alongside deep conservation, each clade we examined also elaborated hypothalamic cell types of its own. We focused on innovations associated with LT subclass 116 (**Fig. 6**), which we hypothesize accompanied the osmotic and thermal challenges of life on land. Our finding that subclass 116 is present in a terrestrial but not an aquatic amphibian species supported this hypothesis. Yet such elaboration was not unique: while many mouse cell types were shared across vertebrates, each non-mammalian lineage carried many of its own, most strikingly zebrafish, which had roughly four times as many unmapped neurons as any tetrapod (**Fig. 5F**). Although our use of the mouse as the mapping reference might inflate this asymmetry, the cell types shared among non-mammalian species and the scale of the teleost expansion together indicate that hypothalamic repertoires have been elaborated repeatedly and independently across the vertebrate tree.

Together, our data suggest several non-exclusive models for the evolution of the neural architecture that supports physiological functions critical for survival and fitness. In the first, a conserved core set of TFs is paired with flexible effectors, allowing functional diversification to meet diverse physiological needs while preserving cell type identity. In the second, multiple regulatory paths converge on similar neuropeptide expression and similar neuronal responses to environmental cues. This would suggest stabilizing selection on gene expression (e.g., of neuropeptides) despite regulatory drift. Integrating these models points to a pattern of mosaic evolution, in which hypothalamic functions evolve at different rates according to the physiological demands of clade-specific specializations. Such a pattern would accommodate both the keystone functions of highly conserved cell types, such as the PVH water-balance neurons, and newer lineage-specific functions such as endothermy in birds and mammals.

## Materials and Methods summary

### Animals

Experiments used adult males of six species: mice (*Mus musculus*, CD1-IGS), prairie voles (*Microtus ochrogaster*), Japanese quail (*Coturnix japonica*), green anoles (*Anolis carolinensis*), tropical clawed frogs (*Xenopus tropicalis*), and mimic poison frogs (*Ranitomeya imitator*). Animals were obtained from commercial suppliers or established breeding colonies and maintained under species-appropriate housing, lighting, diet, and temperature conditions. All procedures were approved by the Institutional Animal Care and Use Committees of Stanford University, Cornell University, and Rhodes College.

### Anatomical references

Whole-brain anatomical references for quail and *Xenopus* were generated by iDISCO-style clearing (*113*), propidium iodide labeling, and light-sheet imaging; anole references were vibratome-sectioned, DAPI-stained, and confocally imaged.

### Single-nucleus RNA sequencing

Hypothalamic regions were microdissected from fresh-frozen brains, and nuclei were isolated, NeuN/DNA-stained, flow-sorted, and used to construct 10x Genomics 3′ v3.1 libraries; distantly related species were multiplexed within libraries to reduce doublets. Libraries were sequenced on a NovaSeq X, aligned with STAR (*114*) to species’ genomes (including a purpose-built *R. imitator* assembly), and processed with standard quality filtering, doublet removal (Scrublet (*115*)), ambient-RNA correction (SoupX), demultiplexing, normalization, dimensionality reduction (SAM (*116*)), batch correction (Harmony (*117*)), and Leiden clustering.

### Cross-species cell-type integration

Subclasses were aligned across species using two complementary approaches—SAMap (*28*) and SATURN (*29*)—each anchored to mouse and prairie vole across 30 resampled runs. Labels were merged into consensus assignments by cross-species and cross-method agreement, low-confidence and discordant cells were excluded, and residual unlabeled populations were clustered *de novo*. Subclasses were assigned to their most recent common ancestor to reconstruct evolutionary gains and losses.

### Cell type, neurotransmitter type, and molecular annotation

Neuronal and non-neuronal identities, neurotransmitter/neuromodulator classes, and supertypes were annotated from curated marker gene panels, with ambiguous cases resolved manually. Differentially expressed genes were called uniformly across species. Neuropeptides and their receptors were drawn from a published catalog, and transcription factors were defined from GO annotations with one-to-one orthologs assigned by EggNOG (*118*, *119*); TF co-regulatory modules and TF–neuropeptide regulatory links were identified in mouse using SCENIC+ (*93*) on our RNAseq data supplemented with published mouse single-cell ATACseq data (*94*).

### Physiological manipulations

To probe conserved hypothalamic responses, subsets of animals underwent water restriction (mice, quail), graded osmotic challenge (*Xenopus*), or acute heat exposure (quail) prior to tissue collection.

### STARmap

Selected hypothalamic sections were profiled by STARmap (*26*); amplicons were imaged over multiple sequencing rounds, decoded, and segmented to assign transcripts to cells for spatial cell-type mapping. Detailed descriptions of all experimental protocols and analyses are provided in the **Supplementary Materials**.

## Acknowledgements

We are grateful to members of the Luo, Wang, O’Connell, and Quake laboratories for providing feedback and comments throughout the project. We thank Karl Deisseroth and Stanford University Cell Sciences Imaging Facility (RRID:SCR_017787) for providing the access to microscopes, and Mark L. Andermann and Alexander L. Starr for critical reading of the manuscript.

## Author contributions

Y.W., P.K., B.C.G., S.R.Q., L.A.O., B.W., and L.L. designed the study; Y.W. collected the single-nucleus RNA-seq data with the help of A.H., B.C.G., J. Lee, and R.C.G.; P.K. and Y.W. generated code and analyzed the data with the help of G.Y., C.C., T.H.S. and J.H.S.; Y.R. performed SATURN mapping under the supervision of J. Leskovec; J.N., L.L.S., D.K., A.G.O. and L.A.O. provided materials; Y.W., B.C.G., and H.M.T. performed STARmap; Y.W. and B.C.G. constructed serial brain sections and annotations with the help of J. H. S.; D.W.K. and H.Z. shared mouse whole-brain transcriptome data prior to publication; Y.W., P.K., B.C.G., L.A.O., B.W., and L.L. wrote the paper with feedback from all authors. B.W. and L.L. supervised the study with the help of S.R.Q. and L.A.O.

## Data and materials availability

Single-nucleus RNA-seq data and the *R. imitator* draft genome will be deposited in GEO and Zenodo (DOI: 10.5281/zenodo.21312279), respectively. Custom analysis code is available at https://github.com/Bo-Wang-Lab/Vertebrate_hypothalamus/. All other data are available in the main paper or supplementary materials. All materials are available through requests to the corresponding authors.

## Supplementary Materials

### Materials and Methods

#### Animals

All animal procedures were approved by the Institutional Animal Care and Use Committee at Stanford University, Cornell University, and Rhodes College.

Adult male CD1-IGS mice (Charles River Laboratory, Hollister, CA, USA) are housed in a laboratory colony (12L12D cycle, lights on at 07:00, ∼21°C) within a ventilated rodent housing rack system (Innovive, San Diego, CA, USA) to house disposable IVC cages (37.3 cm × 23.4 cm × 14.0 cm) bedded with Alpha-Dri (Shepherd Specialty, MA, USA) supplemented with Enviro-Dri as enrichment. Rodents were provided chow (Teklad Global 18% Protein Rodent Diet, Inotiv, Lafayette, IN, USA) and chlorinated water (1.0–3.0 ppm, < 80 ppb trihalomethanes, Aquavive M-WB-300C, Innovive, San Diego, CA, USA) available *ad libitum*, except when placed on water restriction. Experiments were done during the light phase.

Outbred, sexually mature male prairie voles were obtained from the Ophir lab breeding colony at Cornell University, Ithaca, NY, from pairs that were offspring of wild-caught animals in Champaign County, IL. Animals were weaned and housed with littermates on postnatal day (PND) 21, and then housed with same-sex littermates after PND42–45. All animals received rodent chow (Laboratory Rodent Diet 5001, LabDiet, St. Louis, MO, USA) and water *ad libitum*, and were maintained under standard laboratory conditions (14L10D cycle, lights on at 08:00, 20 ± 2°C) in transparent polycarbonate cages (46.5×25×15.5 cm) lined with Sani-chip bedding and provided nesting material. All males used in this study were unrelated and sexually mature, and approximately 60 days old.

Adult male quails (*Coturnix japonica*) were purchased from Stellar Gamebirds, Poultry, and Waterfowl LLC (Ruskin, FL, USA), and AA Egg Labs (Los Angeles, CA, USA) and housed in a laboratory colony (12L12D cycle, lights on at 07:00, ∼21°C) within cages (61.60×33×31.5 cm) bedded with polypads (Shepherd Specialty, Scituate) and straw. We provided dust baths (Sani-Chip, Inotiv) in corrugated cardboard boxes (15.24×22.86×11.43 cm) as enrichment. Quail received chow (Purina Game Bird Maintenance, Arden Hills, MN, USA) and supplemented with grass seeds, oil seeds, and whole grain, along with premium millet spray (Sleek & Sassy, Monroe, OR, USA) and purified water (1.0–3.0 ppm, <80 ppb trihalomethanes, Aquavive M-WB-300C, Innovive, San Diego, CA, USA) *ad libitum*.

Adult male green anoles (*Anolis carolinensis*) were obtained from Carolina Biological Supply and housed singly within terraria (30.5 cm H × 26 cm W × 51 cm L) covered with a wire-mesh lid. Room lighting (14L10D) was supplemented with full-spectrum lighting and heat was supplemented with a 60-W incandescent light bulb secured above one end of the terrarium. Lizards were fed crickets thrice weekly and had access to standing water, as well as being misted daily. Lizards were allowed 19 to 21 days of acclimation to lab conditions before the commencement of experimental procedures.

Wild-type, sexually mature male tropical clawed frogs (*Xenopus tropicalis, Xtr.Ivory-Coast^NXR^*) were purchased from The National *Xenopus* Resource at Marine Biological Laboratory (Falmouth, MA, USA). *X. tropicalis* males (n = 3) were housed together in one 2.5-gallon tank and fed Nasco Frog Brittle (Fort Atkinson, WI, USA) thrice weekly. *Xenopus* were housed in DI water treated with reverse osmosis conditioner (RO Rx, Josh’s Frogs, Owosso, MI, USA), which was refreshed (50% water replacement) every 3 days and maintained at 25°C.

Sexually mature male poison frogs (*Ranitomeya imitator*) were captive bred or purchased from Ruffing’s Ranitomeya (Tiffon, Ohio, USA). Frogs were group housed in terraria (90 × 45 × 45 cm) bedded with a sphagnum moss substrate, driftwood, live Pothos plants, horizontally mounted film canisters as egg deposition sites, and film canisters filled with water (treated with reverse osmosis R/O Rx, Josh’s Frogs, Owosso, MI) for tadpole deposition. Terraria were automatically misted ten times daily for 20 seconds each, and frogs were fed live *Drosophila melanogaster* flies dusted with a vitamin powder and springtails three times per week. The observation housing was set on a 12:12 light cycle from 6:00 to 18:00. The average temperature and humidity of the observation was recorded for each day of observation, usually around 23.5°C and 95% humidity within the tank.

#### Tissue processing for anatomical references

##### Xenopus tropicalis and Coturnix japonica

To generate the quail and frog brain references, we followed the whole-brain clearing method described previously (*30*). Briefly, whole brains were dissected out and postfixed overnight in 4% PFA at 4°C. Unless otherwise noted, all labeling and clearing steps were performed on a rocker at room temperature for 1 hour. Brains were washed 3× in 1× PBS and once in B1n (1:1000 Triton X-100, 2% wt/vol glycine, 1:10,000 10N NaOH, 0.02% sodium azide), then dehydrated stepwise into 100% methanol diluted in B1n (20, 40, 60, 80%), followed by two additional washes in 100% methanol. Brains were incubated overnight in 2:1 dichloromethane (DCM):methanol, then washed twice in 100% DCM and three times in 100% methanol the following day. Brains were bleached for 4 hours in a 5:1 methanol:30% hydrogen peroxide mix, then rehydrated into B1n (80, 60, 40, 20% methanol) and permeabilized in PTxwH (1× PBS, 1:1000 Triton X-100, 1:2000 Tween-20, 2 µg/µL heparin, 0.02% sodium azide) containing 0.3 M glycine and 5% DMSO. Samples were washed 3× in PTxwH and incubated with propidium iodide (Biolegend, Cat #421301, San Diego, CA, USA) for 7 days while rocking at 37°C, then washed 4× in PTxwH at 37°C. Samples were then dehydrated stepwise in methanol (using water in place of B1n; 20, 40, 60, and 80%), followed by three additional washes in 100% methanol. Brains were placed in a 2:1 DCM:methanol mixture overnight at RT, washed twice the following day with 100% DCM, and cleared in ethyl cinnamate (ECI), first for 4 hours in a fresh tube and then stored in a second fresh tube of ECI.

Brains were imaged on an UltraMicroscope Blaze light-sheet microscope (Miltenyi Biotec, Bergisch, Germany) using a 1× objective, N.A. (numeric aperture) 0.1 at 1× optical zoom. Light sheet was generated using objectives with maximal N.A. (∼0.159) and a 560-nm laser. Images were acquired with 60-ms exposure and 5-µm z-step. Brain length (anterior to posterior) was ∼13,000 µm for the quail and ∼6,000 µm for the frog.

##### Anolis carolinensis

Whole brains were dissected out and fixed in 4% PFA in 1× PBS overnight at 4°C, embedded in 4% low-melting-point agarose, and serially sectioned into 200 µm sections with a vibratome (VT1000s, Leica Biosystems, Nussloch, Germany). Sections were stained with DAPI and imaged on a confocal microscope (Zeiss LSM 900) with a 10× lens (EC Plan-Neofluar 10x/0.30 M27).

#### Tissue processing for single-nucleus RNA-sequencing (snRNAseq)

##### Microtus ochrogaster

Voles were anesthetized following the recommended ethical and regulatory guidelines by an overdose of isoflurane gas and euthanized by rapid decapitation. The brains were immediately extracted, frozen on powdered dry ice, embedded in OCT (Tissue-Tek, Sakura Finetek USA, Torrance, CA, USA) in paraffin embedding molds before being stored at –80°C until processing.

##### Coturnix japonica

Quails were anesthetized using carbon dioxide and rapidly decapitated. Whole brains were dissected out within ∼2 minutes and embedded in OCT in a paraffin embedding mold (Peel-A-Way Embedding molds, Ted Pella, inc. Cat# 27110, Redding, CA, USA) in liquid nitrogen vapor, after which they were stored in –80°C until processing.

##### Anolis carolinensis

Anoles were euthanized via rapid decapitation, and brains were immediately dissected out. The olfactory bulbs and spinal cord remnants were removed, and the remaining tissues were rapidly frozen using liquid nitrogen, stored at –80°C until further processing.

##### Xenopus tropicalis and Ranitomeya imitator

Frogs were anesthetized with 20% benzocaine and rapidly decapitated. Brains were rapidly dissected out and embedded in OCT in an aluminum cryomold floating on liquid nitrogen, after which they were stored in –80°C until processing.

#### Single nuclear preparation, purification, and snRNAseq library construction

The full stepwise protocol and details of all reagents are available on Protocols.io (*120*): Briefly, we sectioned the OCT blocks into serial 200 µm sections with a cryostat (Ephredia CryoStar NX50, Kalamazoo, MI, USA) between –14 and –18°C. We placed a glass slide on a cold block cooled by dry ice, transferred the sections onto the cooled slide, dissected the regions corresponding to hypothalamus (**Figure S1**–**S3**) under a dissection microscope, and collected the dissected regions into 1 mL of nuclear isolation mix (NIM3, composed of 9.8 mM Tris pH 8.0, 245 mM sucrose, 49 mM KCl, 4.9 mM MgCl_2_, 0.98 mM DTT, 1× protease inhibitor, 0.2 unit/µL RNasin plus RNAse inhibitor, 0.1% BSA). We homogenized the tissue with a Dounce homogenizer using the loose pestle for 10 times, followed by the tight pestle for 5 times (Fisher 06-434). We collected the homogenized tissue by centrifuge at 400*g* relative centrifugal force (RCF), 4°C for 5 minutes. We treated the homogenized tissue by NIM3 with 1% Triton X-100 for 1 minute on ice to lyse the cell and permeabilize the nuclei. To separate large tissue blocks from the nuclei, we filtered the lysate with a 70-µm filter, followed by a 40-µm filter. To separate small debris from the lysate, we centrifuged the lysate over 1 mL of a cushion solution made of NIM3 with 1% BSA at 400*g*, 4°C, for 5 minutes. We then suspended the pellet with 1 mL of nuclei freezing medium (NIM2 with 10% DMSO and 0.2 unit/uL RNAse inhibitor) and stored the nuclei as aliquots at –80°C until use.

We thawed the 1-mL nuclei aliquot on ice, added 1 mL of nuclei staining buffer (1×PBS, 1% BSA, 0.2 unit/µL RNasin plus RNAse inhibitor), centrifuged at 400*g*, 4°C for 5 minutes. We stained the pellet with 200 µL staining buffer with 1 µL NeuN antibody conjugated with Alexa fluor 488 (MilliporeSigma, Cat# MAB377X) at 4°C for 20 minutes, added 800 µL nuclei staining buffer, pelleted at 400*g*, 4°C for 5 minutes, resuspended in 500 µL nuclei staining buffer with SiR-DNA (Cytoskeleton Cat #CY-SC007, Denver, CO, USA) or Draq5 (Biolegend Cat #424101, San Diego, CA, USA), filtered with a 40-µm filter. We sorted nuclei staining positive (NeuN positive) for vole, and all single nuclei for quail, anole, and *Xenopus* with a Sony SH800 cytometer (Sony, Minato, Tokyo, Japan) into RT mix (24,000 nuclei with 70-µm nozzle, 18,000 nuclei with 100-µm nozzle) and proceeded directly with droplet generation. We did not detect NeuN staining for Japanese quail, anole, and *Xenopus*, likely due to variation in epitopes. Nonetheless, we proceeded with the same staining regardless to maintain consistency in sample preparation to reduce technical variations.

We used 10X 3’ assay v3.1 to construct the snRNAseq library according to the manufacturer’s instructions. To increase nucleus loading while keeping doublet fraction low, we mixed 2 distant species in some libraries (prairie vole with green anole, Japanese quail with poison frog *Ranitomeya imitator*).

#### snRNAseq library sequencing, sequence alignment, and data processing

Libraries were sequenced on a NovaSeq X (28/90/10/10 bp for read 1, read 2, index 1, and index 2) aiming at ≥ 50,000 reads per nucleus. Raw snRNAseq reads were aligned to the genomes listed in **Table S4** using STAR (v2.7.10b or v2.7.11a) (*111*). Low-quality cells were removed by applying minimum thresholds on the number of genes and UMIs detected per cell (**Table S5**). Genes detected in fewer than 3 cells were removed, and doublets, if present, were removed using Scrublet (v0.2.3) with default parameters and automatically set doublet thresholds (*112*). Ambient RNA levels exceeded 10% in mixed libraries from the vole and anole, and were corrected using SoupX (v1.6.2, thresholds listed in **Table S5**) (*121*). Mixed libraries were separated by thresholding the total UMI counts from the other species (**Figure S4–S6, Table S5**).

The *R. imitator* frog data were separated from the quail by aligning to a draft genome (DOI:10.5281/zenodo.21312279; **Table S4**) using kallisto (v0.48.0) and bustools (v0.41.0), as the current assembly (GCA_032444005.1) used for other analyses was not available at the time. To construct the draft genome, brain, lung, eye, testes, tongue, leg muscle, ventral skin, dorsal skin, intestine, liver, and heart tissues from an adult male *R. imitator* were provided (‘tarapoto’ color morph) to Dovetail Genomics for genome sequencing (PacBio HiFi + Omni-C), assembly, and annotation. GCA_032444005.1 was assembled through the Vertebrate Genome Project (*122*) using the same sequencing data. Additional details will be described in an upcoming manuscript (Wu, Kalakuntla, Goolsby et al., in preparation).

The filtered UMI counts were log-normalized, scaled, and dimensionality-reduced in SAM (v1.0.1) (*113*) using default parameters. Batch correction was performed using Harmony (v0.0.9) (*114*) by library, and cells were clustered using Leiden clustering at a resolution of 1.0. Neural clusters with low UMI counts (less than half of the cluster mean and out of the main distribution) were removed. Additionally, clusters with markers dominated by housekeeping/mitochondrial genes and within the bottom 15 percentile of mean UMI counts were removed. These steps led to the removal of 9,322 vole cells (11% of all cells), 1,683 quail cells (2%), 161 anole cells (0.3%), 40 *Xenopus* cells (0.01%), and 5,188 zebrafish cells (8%). The same data was normalized with scTransform and clustered in Seurat using Louvain clustering. Additional clusters without detectable marker genes were also removed. This led to the removal of an additional 2,198 quail cells (3%), and 2,892 anole cells (6%).

#### Annotation corrections for the *X. tropicalis* genome

The *Xenopus* genome (UCB_Xtro_10.0.112.gtf) had two annotation errors, which we manually corrected. ENSXETG00000005500 (incorrectly annotated as *slc17a6*) was removed. ENSXETG00000004362 (incorrectly annotated as *pomc*) was renamed as ‘*pomc*.1_*Gfra*’ because this gene is a homolog of *Gfra*. The correct genes are ENSXETG00000008393 (*slc17a6*) and ENSXETG00000017532 (*pomc*).

#### Mapping subclasses across species using SAMap

To run SAMap (*123*), we performed pairwise tblastx between all species pairs using the longest isoforms of each gene (gene annotations in **Table S4**), other than in the anole where a random isoform was used. Gene pairs with an E-value <10^-6^ were used in SAMap. The processed snRNAseq data from each species were mapped against the Allen Institute whole brain scRNAseq dataset (hereafter the mouse dataset), including subclasses outside the hypothalamus and non-neuronal subclasses. Mouse data were resampled at 250 cells per subclass (roughly matching the number of cells sequenced in other species) 30 times without replacement. Subclasses with fewer than 250 cells were used in full.

To establish an independent validation reference, mouse and vole data were first mapped using SAMap (v0.2.3). The vole data were deliberately over-clustered to capture potential rare populations. To select the optimal clustering resolution, Leiden clustering was performed over a range of resolutions from 5 to 80 in steps of 5. At each resolution, every vole cluster was assigned a mouse subclass label based on the highest alignment score (a measure of transcriptomic similarity), provided that the alignment score exceeded 0.2 and the cluster contained more than 25 cells. The optimal resolution was determined as the level at which the number of uniquely transferred mouse subclass labels plateaued (resolution = 60). A subclass label was retained only if it was assigned in at least 75% of resampling runs. Clusters sharing the same mouse subclass label were then merged into a single cluster.

Similarly, quail, anole, *Xenopus*, and zebrafish data were mapped to the mouse (with resampling) and to the vole. This dual-reference approach was adopted because requiring a cell type to map consistently to homologous types in both mouse and vole—forming a "triangle" of mutually agreed cross-species mappings— imposes a more stringent, higher-confidence label transfer criterion. As with the prior vole mappings, the data were initially over-clustered to capture rare hypothalamic subclasses and to maximize the number of transferred labels. During this round of mapping, we used only vole labels that were consistent in >90% of resampling runs when mapped to mouse (instead of 75%) ensuring that only the highest-confidence vole labels were propagated. The species-to-vole label was then compared with the species-to-mouse label. If the labels agreed, the label was retained; otherwise, the cluster was grouped into the category of “Unlabeled”. Unlike the mouse-to-vole mapping, no alignment score threshold was applied here, as we reasoned that agreement between the species-to-vole and species-to-mouse mappings was itself sufficient.

#### Mapping subclasses across species using SATURN

As an alternative algorithm to SAMap for cross-species comparisons, SATURN (*29*) was used to project cells from different species into a shared embedding. Briefly, SATURN first identifies genes with similar properties across species, inferred from protein language model (ESM2-15b (*124*)) embeddings, and projects them into a shared feature space (‘macrogenes’). Using these macrogene features, SATURN learns a joint cellular embedding by pretraining an autoencoder with zero-inflated negative binomial loss to reconstruct gene expression, then fine-tuning the encoder with weakly supervised metric learning: predicted cell type clusters from each species are used to pull cells with similar embeddings from different species closer together while pushing apart cells of different types within the same species. SATURN was run on each species’ dataset, paired with mouse or vole data, using 8,000 highly variable genes, 3,000 macrogenes, 200 pretraining epochs, and 50 metric learning epochs. Each pairing was run 30 times, each using either a different mouse subset (as in the SAMap mapping above) or the whole vole data and a different SATURN training seed. To transfer labels from mouse or vole, a 1-nearest-neighbor classifier (*125*) was fit on the 256-dimensional SATURN embeddings and applied it to each non-mouse/non-vole species, independently for all 30 seeds and species pairs. The triangular mapping strategy was applied to SATURN-based label transfer as well. The subclass label from species-to-mouse mapping and species-to-vole mapping within each cluster was compared and the label was kept only if the two labels were concordant and if the cluster contained more than 25 cells. This process was repeated across 30 mappings.

#### Generating consensus subclass labels

A cluster was assigned a consensus label if it received a label from either SAMap or SATURN alone, or from both if their labels agreed. When SAMap and SATURN assigned different labels (2.7% of vole cells, 1.4% of quail, 2.2% of anole, and 0.3% of *Xenopus*), the cluster received a combined name (e.g. *Mm*_077_106) and was excluded from downstream analyses.

Labels received from the species-to-mouse mappings alone were also retained if SAMap and SATURN labels agreed. For each labeled subclass, mapping confidence was quantified as the fraction of resampling runs in which the SAMap or SATURN mouse-label matched the consensus label (**Fig. 2A**).

A small number of unlabeled cells were manually supplemented to subclasses 104 and 126 in anole (87 and 41 cells, respectively) and to subclass 104 in *Xenopus* (109 cells) based on differential expression of specific neuropeptides and TFs: cells added to subclass 104 expressed *Agrp*, *Isl1*, and *Otp* and were GABAergic and cells added to subclass 126 were glutamatergic and expressed *Pomc* and *Isl1*. The manually added cells comprised ∼50% of labeled cells in the anole subclass 104, and ∼20% of *Xenopus* subclass 104. These cells, however, might be convergent in their expression of the marker genes. As such, a note has been appended in the data (‘manually labeled’) to distinguish manually labeled neurons from algorithmically labeled neurons within the same subclass.

#### Mapping non-mammalian subclasses

For each species, Leiden clustering was performed across a range of resolutions, and the adjusted Rand score (scikit-learn v1.0.2) between the resulting clusters and the assigned subclass labels was computed on the labeled cells. The resolution that maximized the adjusted Rand score while minimizing the number of clusters was selected and then applied to re-cluster the unlabeled cells within each species; the selected resolutions were 0.4 for vole, 0.8 for quail, 0.6 for anole, 1.0 for *Xenopus*, and 0.8 for zebrafish, respectively. The unlabeled clusters were treated as candidate subclasses without a mouse homolog.

These unlabeled subclasses were then mapped pairwise between non-mammalian species using SAMap. Two subclasses were considered mapped if they were reciprocal best matches amongst unlabeled clusters and had an alignment score larger than 0.2. Unlabeled clusters from the same species with overlapping mapping relationships to subclasses in other species were merged. We then manually examined the molecular signatures of each candidate subclass, retaining only labels that shared at least 2 differentially expressed TFs across all mapped organisms.

#### Neuronal and non-neuronal labels

Non-neuronal cells were separated from neurons in quail, anole, *Xenopus*, and zebrafish, using gene markers and thresholds listed in **Table S6**, clustered at a Leiden resolution of 3.0, mapped to mouse non-neurons (downsampled to 2,000 cells per subclass), and annotated with the mouse label having the highest alignment score, provided the alignment score to the mouse non-neuronal subclass exceeded 0.2 and the cluster contained >25 cells. Non-neuronal clusters with the same label were merged and the non-neurons that did not pass these thresholds were grouped into the category of “Unlabeled”. Mouse subclass labels were combined in two cases: astrocytes (316, 317, 318, 319, 320) and BAMs/microglia (334, 335), yielding populations labeled 339-astrocyte-like, and 340 macrophages). These two subclass numbers are not used in the mouse brain atlas. This was done because multiple transcriptomically similar mouse subclasses had similar alignment scores (difference in SAMap score between best mapping and second best mapping < 0.2 and ratio < 2) to a single cluster in 4 and 2 species respectively.

Although non-neurons were depleted by FACS prior to sequencing in the vole, a small number remained, likely because FACS separation was not complete. These non-neuronal cells were identified using marker genes in **Table S6**. Because the non-neuronal fraction was very low (<2% of sequenced cells), it could not be analyzed reliably and was not used in downstream analysis.

#### Assigning subclasses to most recent common ancestor

The number of subclasses gained over the vertebrate phylogeny in **Fig. 5F** was determined by assessing how recently each subclass appeared. For example, of the 17 subclasses in the MRCA of vertebrates, 6 were found in all sequenced species, 6 (*cj_xt_dr_2*, *cj_ac_xt_dr_4*, *ac_dr_1*, *cj_xt_dr_1*, *cj_ac_xt_dr_1*, and *cj_ac_xt_dr_3*) were traceable back to zebrafish but were absent in mice, and 5 (075, 077, 091, 105, and 123) were found in both zebrafish and mammals but appeared lost in at least one other species. Similarly, we determined that 42 subclasses were likely present in the MRCA of tetrapods, and that 13 were gained in the MRCA of amniotes.

To count the subclasses lost over the vertebrate phylogeny, we identified subclasses present in at least two species of an outgroup (e.g., anamniotes) but absent from the corresponding in-group (e.g., amniotes). For example: 7 subclasses (074, 077, 091, 103, 105, 139, 143) were found in *Xenopus* or zebrafish and mouse but not in quail or anole.

#### Determining relative location of neuronal subclasses to hypothalamus from MERFISH data

A subclass was determined to be within, near the boundary of, or far away from the hypothalamus by looking at the location of the majority of cells within the subclass using the Allen Brain Atlas MERFISH-C57BL6J-638850 common coordinate framework (CCF). Classes 11–16 of the whole-mouse-brain atlas (*21*) were defined as hypothalamic; classes 08, 09, 10, 18, 22, 23, and 26 as proximal to the hypothalamus; and classes 01–07, 17, 19–21,24–25, and 27–29 as distal to the hypothalamus. Mouse MERFISH data were registered to the Allen Brain Atlas MERFISH-C57BL6J-638850 common coordinate framework (CCF) throughout the paper (*21*, *126*).

#### Neurotransmitter type analysis

Neurotransmitter and neuromodulator types were annotated using a modified version of a previously described marker gene list (8) (**Table S7**). Because expression was evaluated at the subclass level, small within-subclass populations of a different neurotransmitter type were sometimes obscured by averaging: for example, mouse subclass 106 contains dopaminergic neurons (887 of 6,150 cells) but is labeled GABAergic in **Fig. 2**, reflecting the majority assignment. The anole 092 (histaminergic) subclass also expressed *Tph1*, *Slc17a6*, *Ddc*, and *Slc18a2* but *Tph1* and *Slc17a6* expression arose from a single supertype within the subclass that lacked the TFs otherwise characteristic of 092 across species, so the subclass was labeled GABA-Hist. Vole subclass 123 contained only a small fraction of cells expressing GABAergic markers, so they were not labeled GABAergic. *Xenopus* subclass 101 and cj_ac_xt_dr_4 were labeled glycinergic based on expression of the glycine transporter GlyT2; whether they transmit glycine also requires further experimental validation. For the ease of visualization, in **Fig 1E**, non-GABAergic and non-glutamergic neurons with different secondary neurotransmitters were combined (e.g. Gly-GABA and Gly-Glut were both labeled as Gly).

#### Differential expression (DE) analysis

We used “differentially expressed” and “enriched” interchangeably for genes highly expressed in specific cell types compared to others in the paper. They were quantitatively identified as follows. Prior to testing for enrichment, genes were ranked within each subclass using the marker-gene-ratio function in SAM, a metric quantifying the expression of a gene within a cell group relative to its expression across all remaining cells, and the top 7,000 genes per subclass were retained. A gene was labeled as enriched in a given subclass if, relative to at least one of the subclass’s 10 closest hypothalamic neighbors in PC space (by Euclidean distance), it showed either (i) >2-fold enrichment or (ii) a large difference in detection rate between foreground and background cells (q.diff.th > 0.8) and detection in at least 1% of cells within the subclass (8), together with a Bonferroni-corrected Wilcoxon rank-sum p-value < 0.05 when compared to all other hypothalamic cells. Subclasses that mapped outside the hypothalamus were excluded from background for DE analysis, from gene expression normalization (**Figs. S17** to **S34**), and from all correlation calculations.

#### Ortholog assignment

Genes across all organisms were assigned to orthogroups using eggNOG-mapper (v2.1.12) (*115*, *116*) with default parameters at the vertebrate taxonomic level. Where multiple paralogs in a species belong to the same orthogroup, a single representative gene was chosen for that species based on existing annotations (**Table S8**). If no annotation existed, we chose the gene with the highest Pearson correlation to the mouse ortholog across subclass pseudobulk means, restricted to genes detected in more than 0.01% of cells. The correlation-based selection was used to account for paralog substitutions (*28*) and primarily affected ortholog assignment in zebrafish, owing to the whole genome duplication. The expression threshold was set to prevent genes found in a small number of cells from skewing the Pearson correlation. Two TFs not captured by this strategy [*Lmx1a* (zebrafish) and *Zic4* (anole)] were added manually based on cross-species BLAST; in both the percent identity with the mouse ortholog was greater than 85% to their respective mouse ortholog. Additionally, *Pomc* was manually added with putative orthologs across all 6 species.

#### Neuropeptide analysis

The list of neuropeptides and neuropeptide receptors was obtained from a published catalog of candidate mouse and human neuropeptides and receptors (*127*), accessed via <u>neuropeptides.nl</u> (last accessed 20 October 2025). We manually added the following neuropeptides that were not in the database: *Prok1*, *Prok2*, *Npvf*, *Tac2, Kiss2*, *Insl3*, *Igf3*, *Gnas*, *Enho*, *Cgrp*, *Dbi*, and *Pcsk1n*. Receptors were obtained from IPUHAR/BPS GtoPdb database (v2026.2) (*128*) alongside additional manually identified receptors that were not in the database. A full list of neuropeptides and receptors is provided in **Table S9.** Neuropeptides with orthologs in at least 4 species and differentially expressed in any subclass in at least 4 species were plotted in **Fig. 2F**. *Pomc* was included despite the fact that it was only conserved in 3 species due to its relevance in the feeding behavior. Normalized expression of all neuropeptides/receptors found in at least 4 species were plotted in **Figs. S17** to **S29**.

#### Transcription factor analysis

The MGI DNA-binding TF list (GO:0003700, accessed 3 April 2025) was used to identify TFs. The list was manually curated to remove transcriptional coregulators (e.g., *Ski*, *Skor1*) and to add known TFs that were missing. The curated list is provided in **Table S10**. In **Fig. 3A, B, D,** and **E**, TFs with assigned orthologs and expression across all 6 species were used, yielding a set of 454 TFs. All TFs that are differentially expressed in at least 4 species in any subclass were plotted in **Fig. 3A** alongside *Pgr*, which was included to complete module 7. In **Fig. 3F** and **G**, TFs with orthologs across 4 of the 6 species were used. TF family annotations were obtained from JASPAR (v11) (*129*); TFs absent from JASPAR were annotated using GeneCards (*130*).

TF modules were defined as cliques of a graph, groups of nodes in which every node is connected to every other node in the group, where nodes are individual TFs and edges connected TF pairs with a Pearson or Spearman correlation above 0.3 in at least 5 of the 6 species (*131*). Two modules were manually combined (*Uncx-Lhx5/Sim1-Otp* and *Pgr-Ar/Pgr-Esr1*) based on cross-member correlations that exceeded 0.3, even though not all members had pair-wise correlations above 0.3. This yielded 323 unique modules consisting of 226 unique genes. A manually curated selection of the modules is shown in **Fig. 3C** and the complete set is provided in **Table S11**.

Conserved coregulatory modules were identified by testing whether pairs of TFs within these modules showed reciprocal regulation, using SCENIC+ (v1.0a1) (*132*). Mouse hypothalamic snATACseq and scRNAseq datasets were obtained from previously published atlases (*21*, *133*); to balance the two modalities, the mouse data was subset to region_of_interest_acronym "HY", then downsampled to 800 cells per subclass. Cell-by-gene and cell-by-peak matrices were used as SCENIC+ input, with an FDR of 0.001 for motif clustering, a log_2_ fold-change threshold of 1.0, an AUC threshold of 0.005, and an adjusted p-value threshold of 0.05 for motif enrichment. Regulatory links between TFs and downstream genes were inferred using SCENIC+’s GRNBoost2 implementation with a rho threshold of 0.05, retaining only TFs with at least 10 target genes. Gene annotations were taken from Ensembl (nov2020 archive, *M. musculus*); all other parameters were left at default.

Downstream target genes were selected in two steps. First, non-TF genes predicted by SCENIC+ to be positively regulated by at least one member of the TF coregulatory module were identified, and only those with an average Pearson correlation (calculated on subclass pseudobulk means) above 0.3 with the two module members were retained. Of this set, the 10 genes with the highest maximum Pearson correlation to either member of the coregulatory module in the mouse were selected. Only genes with assigned orthologs in all 6 species were used. For each species, the maximum Pearson correlation between these 10 genes and either module member was calculated and plotted, rather than the average, to illustrate the maximum degree of expression conservation between the TF modules and their downstream genes across species.

TF–neuropeptide pairs were defined as those with a Pearson correlation above 0.3 in mouse for which SCENIC+ additionally supported positive regulation of the neuropeptide by the TF. As a non-target comparison set, for each TF we identified non-neuropeptide/non-TF genes in the mouse whose correlation with that TF was closest to the mouse TF–neuropeptide correlation, drawing five times as many non-target genes per TF as it had regulated neuropeptides, and restricting to genes with orthologs in at least 4 of the 6 species to match the criterion applied to neuropeptide-TF relationships. TF, neuropeptide, and non-target gene for each species are listed in **Table S8**.

#### Random sparsity matched genes for comparison with TF-TF co-expression

To prevent bias in co-expression analysis due to dropout, sparsity-matched control genes were generated by dividing the 454 one-to-one orthologous TFs into 10 decile bins by detection rate (the fraction of cells in a subclass expressing the gene averaged across the 6 species), assigning all non-TF genes to the same bins, and randomly drawing non-TF genes to match the per-bin TF distribution. Random selection was repeated 300 times to obtain a mean and 90% confidence interval of the number of conserved co-expressed genes.

#### Supertype labeling

Mouse PVH and LT supertypes were labeled using neuropeptide and TF expression together with previously described functions. Mouse PVH supertype 0588 contained both Avp+ and Oxt+ clusters and was split into two supertypes, Avp^+^ and Oxt^+^. Additionally, Mouse PVH supertype 0587 contained both Crh+ and Trh+ clusters and was split into two supertypes, Crh+ and Trh+.

Supertypes in the PVH and LT of the other species were labeled by first subsetting the subclass of interest within each species in which it was present, then mapping cells from that subclass using SAMap across all species pairs sharing the subclass. A range of Leiden clustering resolutions was tested for each subclass to maximize the number of mapped mouse supertypes. A resolution of 2.0 was selected for the PVH across all species, and resolutions of 1.0, 0.5, 1.0, and 0.25 were selected for the LT in the vole, quail, anole, and imitator frog, respectively. The resulting clusters were used to identify homologous supertypes, with each cluster labeled by the mouse supertype having the highest SAMap alignment score. Clusters sharing the same label were then grouped. Clusters with maximum alignment scores below 0.2 to the mouse supertypes were excluded from analysis.

#### Experimental conditions for dehydration

##### C. japonica

In accordance with prior literature (*134*), quails were water-restricted for 48 hrs before collection. No mortality or apparent signs of distress were observed.

##### M. musculus

In accordance with prior literature (*97*), mice were water-restricted for 48 hrs before collection. No mortality or apparent signs of distress were observed.

##### X. tropicalis

For osmolarity assays, animals were exposed to 200 mL water solutions with increasing osmolarity (235 mOsm [baseline plasma saline level], 310 mOsm, 400 mOSm, 600 mOSm [published tolerance extreme, (*135*)] in 32 oz plastic deli containers exposed to open air, with one frog receiving each treatment, starting from lowest salinity. If frogs move frequently or move to the surface of the water immediately (which *Xenopus* do in the wild when leaving high-salinity pools in Africa to migrate to less salty ponds), the frog was removed from the trial and returned to its home tank. Once we identified the highest salinity tolerable without evoking distress (moving every ≤ 10 seconds, or immediately attempting to escape to the surface), we recorded *Xenopus* in this saline solution for 1 hr, after which frog tissue was collected. The osmolarity saline solution evoking dehydration was found to be between 310 mOsm and 400 mOsm. 400mOsm was used for the STARmap experiment shown in **Fig. 6**.

#### Experimental conditions for heat treatment

##### C. japonica

Quails were removed from home cages and placed into a warm incubator (Scigen model 2000) at 37°C for 1 hr. This is lower than the lethal temperature for Japanese quail (40–45°C) (*136*).

#### STARmap

We performed STARmap as described previously (*137*) with recent improvements (*138*, *139*). Mice and quails were euthanized by CO_2_ followed by cervical dislocation. *Xenopus* were euthanized by benzocaine followed by cervical dislocation. After euthanasia, we quickly dissected the brain and embedded the brain in OCT, which was then snap frozen in liquid nitrogen vapor until the block was solid. The tissue block remained under –80℃ until use. We sectioned the tissue block into 10-µm sections with a cryostat (Leica CM1900 UV, Wetzlar, Germany) at –20℃ and mounted the sections onto a silanized glass slide. We dried the sections briefly at room temperature (RT), fixed the sections with 4% PFA for 10 minutes, washed the sections with 1× PBS 3 times briefly, permeabilized the sections with precooled methanol for 1 hour under –80℃.

We performed the following steps under room temperature (RT) between 20–25℃ unless specified. After permeabilization, we brought the slide to RT and rehydrated with PBSTR (0.1 u/µL SUPERasin, 1× PBS, 0.1% Tween-20) for 2 minutes. We hybridized slides in hybridization mix (0.01 µM each probe, 2× SSC, 10% deionized formamide, 1% Tween-20, 20mM Ribonucleoside Vanadyl Complex [RVC, Cat #S1402S New England Biolabs, Ipswich, MA, USA], 0.1 mg/mL salmon sperm DNA) at 40℃ overnight.

After hybridization, we incubated the sections with PBSTV (1× PBS with 0.1% Tween-20, 2 mM RVC) for 20 minutes twice, followed by 4× saline-sodium citrate (SSC) in PBSTR for 20 minutes at 37℃. Slides were then washed with PBST (1× PBS with 0.1% Tween-20) briefly, then incubated in a ligation mix (0.1u T4 ligase, 1× T4 ligase buffer, 0.2u SUPERasin, 0.1 mg/mL BSA) for 2 hours. We washed slides in PBSTR for 2 minutes twice, then incubated slides in RCA mix (1× Phi29-XT DNA polymerase [New England Biolabs, Cat #E1603S], 1× Phi29-XT buffer, 1mM each of dNTP, 0.08 mM 5-3-aminoallyl-dUTP, 0.4u SUPERasin, 0.1mg/mL BSA) for 4 hours at 42℃. Slides were then washed in PBST for 2 minutes twice, and incubated in 20 mM methacrylic acid N-hydroxysuccinimide ester (MilliporeSigma, CAT#730300) in 0.9% sodium bicarbonate overnight.

We then washed sections briefly in PBST, incubated the sections in monomer solution (4% acrylamide, 0.2% N,N’-methylenebiacrylamide—TEMED for short—in 2× SSC) for 30 minutes. We added 10 µL gel solution (monomer solution, 0.2% TEMED, 0.2% ammonium persulfate—APS for short—, and 0.25% 2,2′-Azobis [2-(2-imidazolin-2-yl)propane] dihydrochloride, also known as VA-044) to the center of each section, covered the sections with a coverslip coated with Gel Slick solution (Lonza, Basel, Switzerland) and allowed the gel to polymerize for 1 hour at 37℃. After the polyacrylamide gel formed, we immersed the slide with PBST in a jar, and removed the coverslip carefully with forceps. We then incubated the gel-sections with proteinase K digestion buffer (2× SSC, 1% SDS, and 1 mg/mL proteinase K) for 1 hour at 37℃, followed by a 5 minute wash with 1 mM 4-(2-Aminoethyl) benzenesulfonyl fluoride hydrochloride (AEBSF) to inhibit remaining proteinase K. We then washed the gel-sections with PBST briefly and stored the gel-sections in wash buffer (2× SSC, 10% deionized formamide) in 4℃ in a jar until sequencing.

Upon sequencing, we washed the gel-sections twice with stripping buffer (60% formamide, 0.1% Triton X-100) and then dephosphorylated with Antarctic phosphatase (0.2 mg/mL BSA, 1× reaction buffer, 0.1 unit/µL Antarctic phosphatase, Cat #M0289S, NEB) for 1 hour under 37℃. We then washed gel-sections with wash buffer in a jar for 5 minutes. For each round of sequencing, we incubated the gel-sections in ligation mix (4% PEG8000, 0.25µM of F1-488, F3-647, F4-750 probes, 0.05 µM of F2-546 probe, 1µM OR probe, 1× T4 DNA ligase buffer, 0.1 unit/µL T4 DNA ligase) for 1–3 hours, washed the gel-sections with wash buffer with DAPI (1:1000 from stock) for 10 minutes 3 times. We imaged the amplicons using a Nikon Eclipse spinning disk microscope with 40× oil lens (Plan Fluor 40×/1.30 oil, WD 0.24) or Zeiss LSM 980 Airyscan confocal microscope (Zeiss, Oberkochen, Germany) with a 40× glycerol lens (LD LCI Plan-Apochromat 40x/1.2 Imm Corr DIC M27). After each round of sequencing, we removed the fluorophores with stripping buffer for 10 minutes 3 times under 40℃ with rotation, then with wash buffer for 5 minutes, and stored the sections in the same buffer under 4℃ if not imaged immediately.

Oligonucleotides used in the STARmap libraries were listed in **Table S12**.

#### STARmap data processing

We analyzed the STARmap data using the maximal intensity projected image of the z stacks. We performed pixel classification using ilastik (*140*) to generate probability maps for amplicon (dot-like) signals and DAPI staining. Images from different sequencing rounds were aligned by identifying Speeded-Up Robust Features (SURF) in the DAPI channel and warping each round to match shared features. Amplicons were called using the “Find Maxima…” function in FIJI; this approach undercounts amplicons in crowded regions and for genes with very high expression. Cell boundaries were segmented with Baysor (*141*) using the parameters:

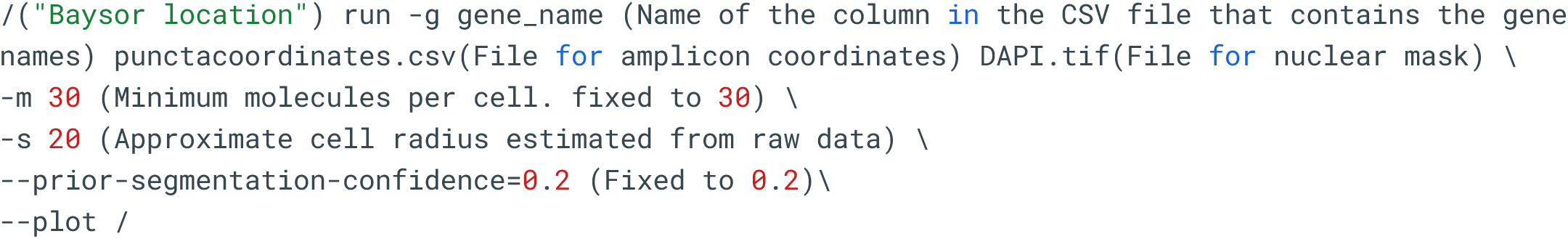

We used the DAPI channel as a nuclear mask for Baysor. The amplicons with the cell’s boundary were counted in R. A cell by gene count table was generated. Cells without any amplicon were discarded. Cells were hierarchically clustered by marker gene (such as *VT, OT, Crh, Sst, Trh,* and *Nkx2-2* for subclas 133) expression and manually annotated based on known marker gene patterns (e.g. *VT* supertype expressed *VT* at a much higher level than other supertypes within subclass 133) or the pattern seen in the snRNASeq data (e.g. *Ucn* supertype in *Xenopus* 133 expressed both *Trh* and *Ucn*). For LT STARmap analysis, the images were processed the same way described above and *Fos* amplicons were counted manually in FIJI.

---

**Fig. S1.**
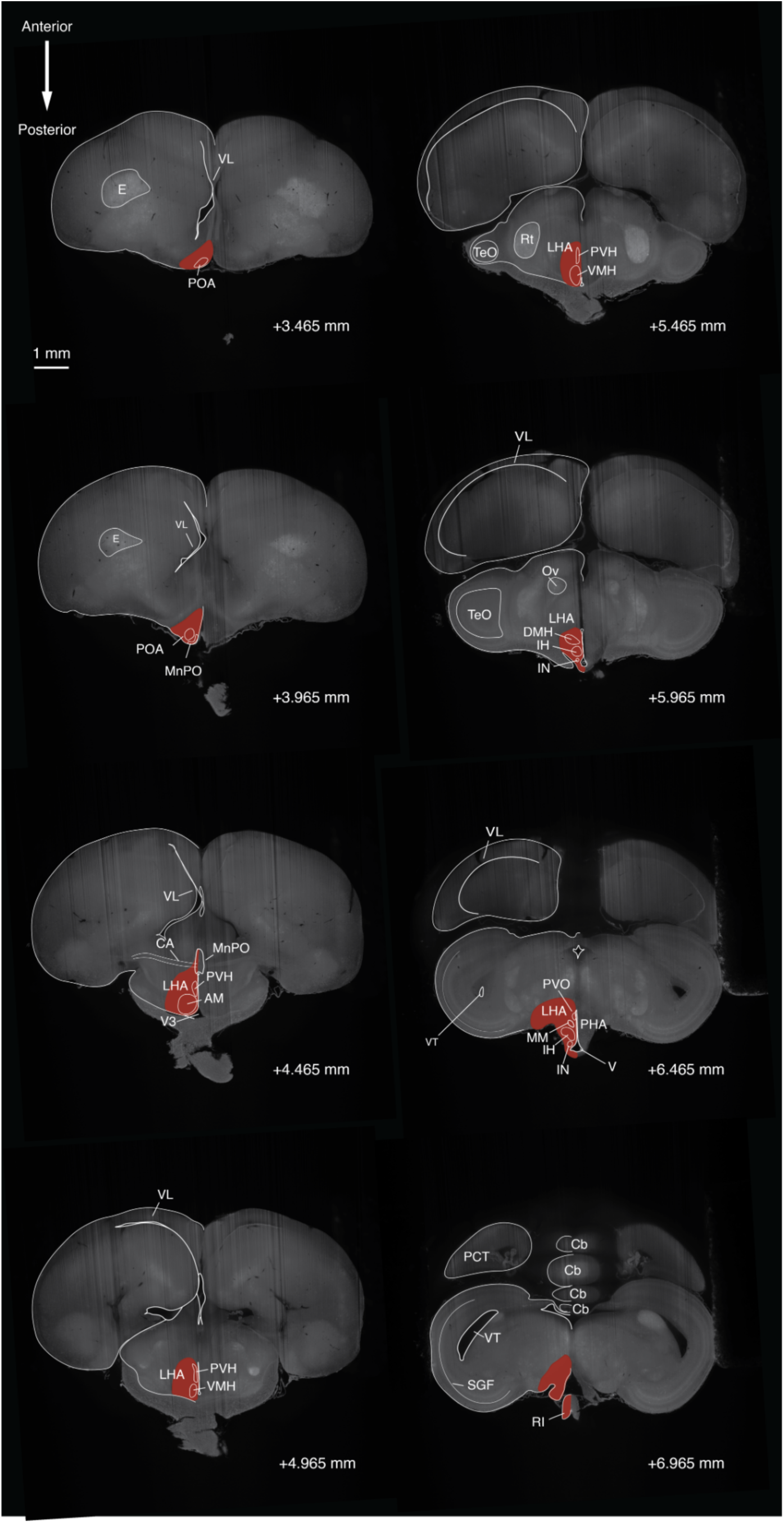
Anatomical reference for Japanese quail (*Coturnix japonica)* hypothalamus dissection. Serial coronal sections of *C. japonica* brain stained with propidium iodide that contains hypothalamus, visualized via light sheet microscopy. The + mm values at the bottom right represent the distance of the section from the anterior end of the telencephalon. Regions for cryodissection for snRNAseq are indicated in red (only one side is indicated, but we dissected both sides). Labels with white borders represent approximate locations of select landmark brain regions and major hypothalamic nuclei adapted from (*145*, *146*). **Abbreviations:** AM, anterior medialis nucleus; CA, anterior commissure; Cb, cerebellum; DMH, dorsomedial hypothalamus; E, ectostriatum; IH, inferior hypothalamus; IN, infundibular nucleus; LHA, lateral hypothalamus; MnPO, median preoptic nucleus; OV, nucleus ovoidalis; PCT, posterior/caudal end of telencephalon; PHA, posterior hypothalamic area; POA, preoptic area; PVH, periventricular nucleus of the hypothalamus; PVO, paraventricular organ; RI, infundibular recess; Rt, nucleus rotondus; SGF, stratum griseum et fibrosum superficiale; TeO, Optic Tectum; V3, third ventricle; VL, lateral ventricle; VMH, ventromedial hypothalamus; VT, mesencephalic-tectal ventricle.

**Fig. S2.**
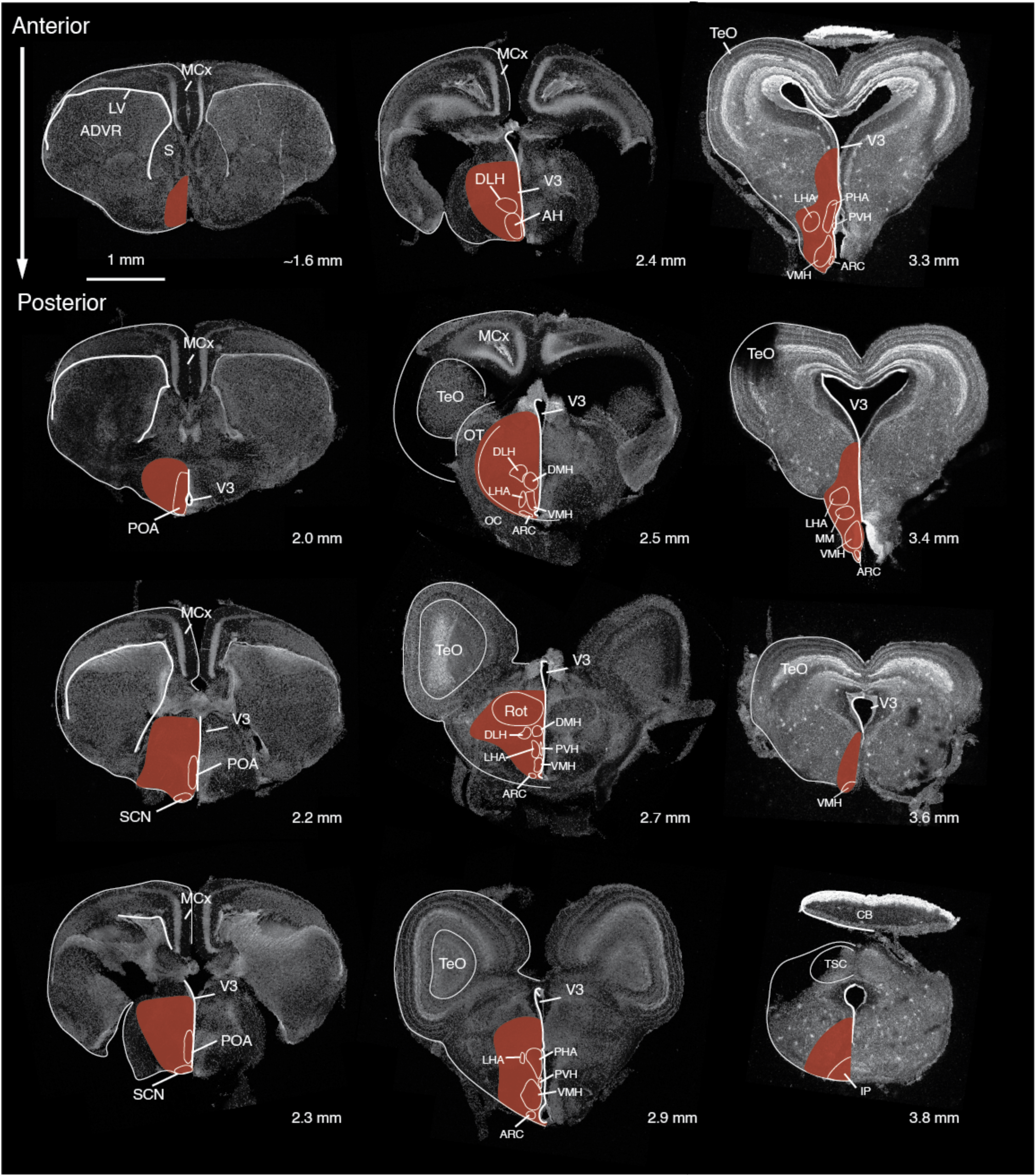
Anatomical reference for green anole (*Anolis carolinensis)* hypothalamus dissection. Serial coronal sections of *A. carolinensis* brain (DAPI stain), visualized via confocal microscopy. The + mm values at the bottom right represent the distance of the section from the anterior end of the telencephalon. Regions for cryodissection for snRNAseq are indicated in red (only one side is indicated, but we dissected both sides). Labels with white borders represent approximate locations of select landmark brain regions and major hypothalamic nuclei adapted from (*2*, *122*, *147*). **Abbreviations:** ADVR, anterior dorsal ventricular ridge; AH, anterior hypothalamus; ARC, arcuate nucleus; CB, cerebellum; DLH, dorsolateral hypothalamic nucleus; DMH, dorsomedial hypothalamic nucleus; IP, interpeduncular nucleus; LHA, lateral hypothalamus; LV, lateral ventricle; MCx, medial cortex; OC, optic chiasm; OT, optic tract; PHA, posterior hypothalamic area; POA, preoptic area; PVH, paraventricular hypothalamus; Rot, rotund nucleus; S, Septum; SCN, suprachiasmatic nucleus; TeO, optic tectum; TSC, torus semicularis, central; V3, Third Ventricle; VMH, ventromedial hypothalamus.

**Fig. S3.**
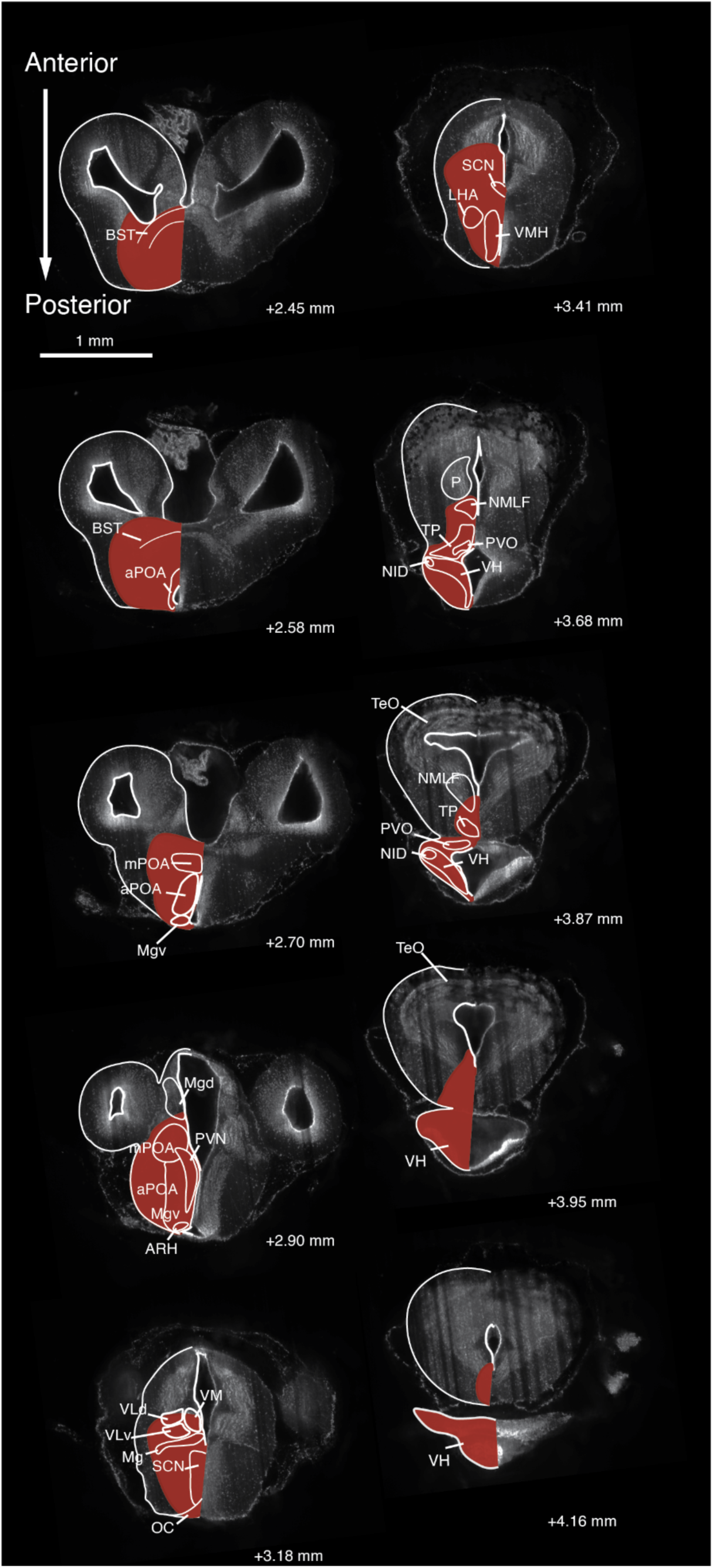
Anatomical reference for tropical clawed frog (*Xenopus tropicalis)* hypothalamus dissection. Serial coronal sections of *X. tropicalis* brain stained with propidium iodide, visualized via light sheet microscopy. The + mm values at the bottom right represent the distance of the section posterior to the anterior end of the olfactory bulb. Regions for cryodissection for snRNAseq are indicated in red (only one side is indicated, but we dissected both sides). Labels with white borders represent approximate locations of select landmark brain regions and major hypothalamic nuclei. Note that we revise some locations of hypothalamic nuclei in the amphibian reference atlas, which is a modest departure from the prior published atlas (*60*). The additional atlases were referenced: (*148*–*154*). Note that we specified a location for the paraventricular hypothalamus (PVH) from our study on subclass 133 (see **Fig. 4** for detail). We additionally identified proposed locations the nucleus infundibularis dorsalis (NID) and arcuate nucleus (ARH) from populations of galanin neurons and *POMC*/*Agrp* neurons, respectively. The paraventricular organ is specified based on expression of serotonin and location along the fourth ventricle, in agreement with where this nucleus was originally proposed to be (*149*, *152*). **Abbreviations:** aPOA, anterior preoptic area; ARH, arcuate nucleus; BST, bed nucleus stria terminalis; Dp, dorsal pallium; H, habenula; La, lateral thalamic nucleus; LHA, lateral hypothalamus; Lp, lateral pallium; Mg, magnocellular preoptic nucleus; Mgd, magnocellular preoptic nucleus, dorsal; Mgv, magnocellular preoptic nucleus, ventral; Mp, medial pallium; mPOA, medial preoptic area; NMLF, nucleus of the medial longitudinal fasciculus; OT, optic tectum; P, posterior thalamic nucleus; Pd, pallidum; PVH, paraventricular hypothalamus; PVO, paraventricular organ; SCN, suprachiasmatic nucleus; Tel, telencephalon; TP, posterior tuberculum; VLd, ventrolateral thalamic nucleus, dorsal; VLv, ventrolateral thalamic nucleus, ventral; VM, ventromedial thalamic nucleus; VH, ventral hypothalamic nucleus.

**Fig. S4.**
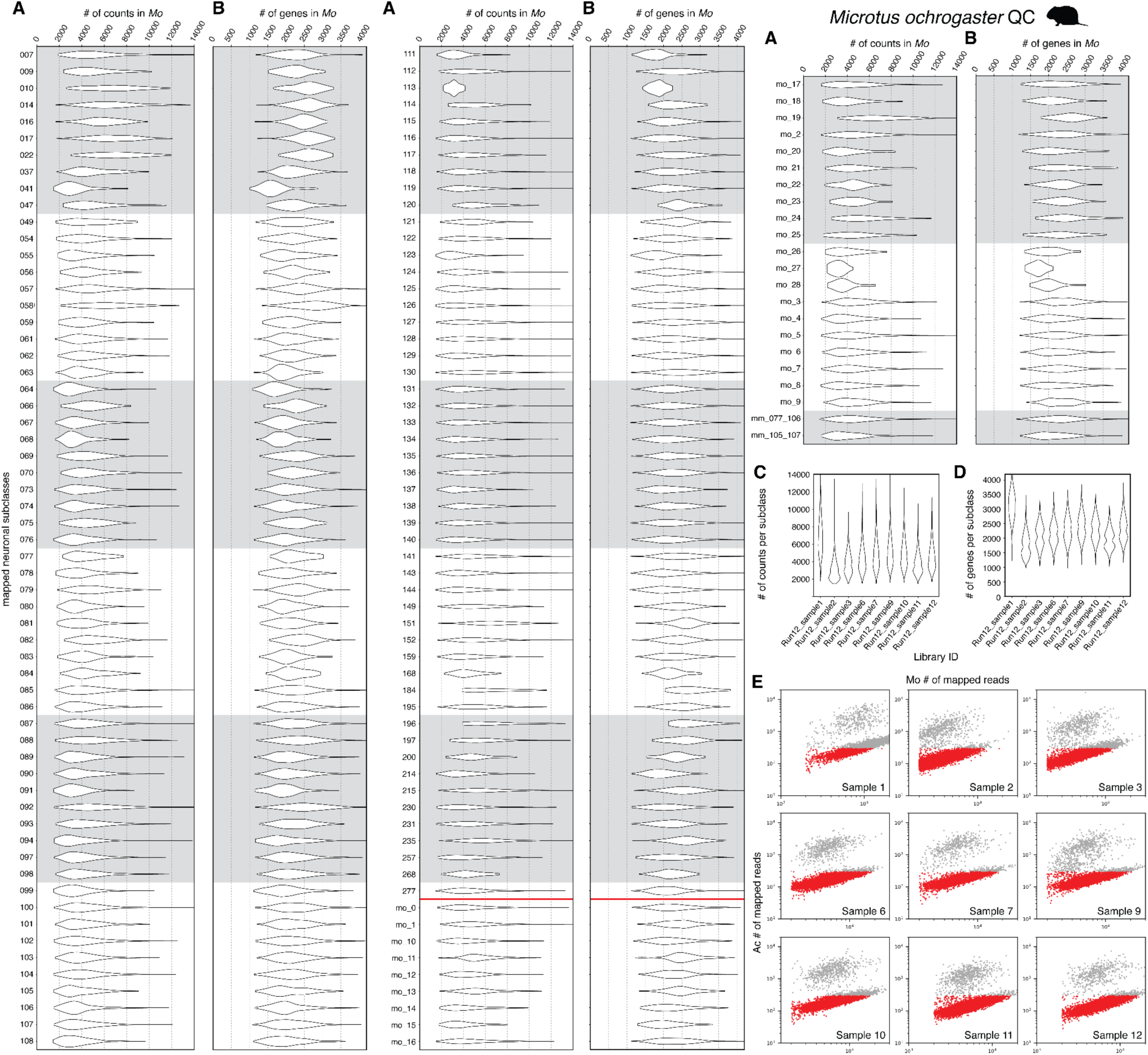
Quality metrics of snRNAseq data for prairie vole (*Microtus ochrogaster*). **(A)** Number of UMI counts per subclass (*21*). This list is larger than the set shown in **Fig. 2**, as it includes both the hypothalamic and non-hypothalamic neuronal subclasses. Non-neuronal subclasses were not included, owing to their removal for *M. ochrogaster* (**Methods**). **(B)** Number of detected genes per subclass. **(C)** Number of counts/subclass per *M. ochrogaster* library. **(D)** Number of genes/subclass per *M. ochrogaster* library. **(E)** Separation of cells in mixed-species libraries. Cells (highlighted in red) were selected for analysis based on the number of mapped reads to a particular genome (*M. ochrogaster* or *A. carolinensis*).

**Fig. S5.**
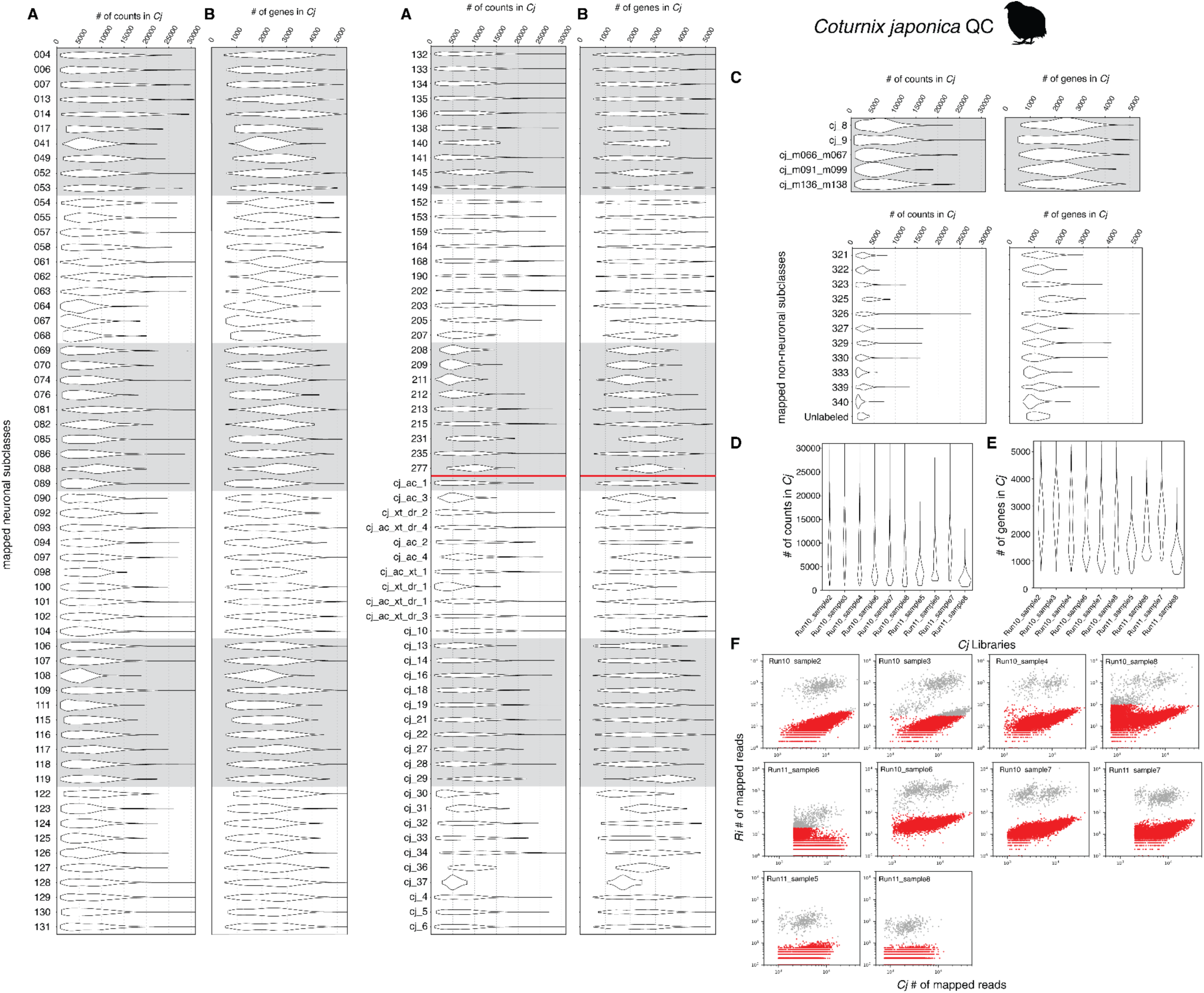
Quality metrics of snRNAseq data for Japanese quail (*Coturnix japonica*). **(A)** Number of counts per subclass(*21*). This list is larger than the set shown in **Fig. 2**, as it includes the mapped hypothalamus neuronal subclasses, non-hypothalamic neuronal subclasses, unmapped subclasses, and non-neuronal subclasses. All neuronal subclasses following the red horizontal line are not found in mice. **(B)** Number of genes per subclass. For (A) and (B), the top section of rows represents neuronal subclasses, and the bottom section represents non-neuronal subclasses. **(C)** Number of counts/subclass per *C. japonica* library sample. **(D)** Number of genes/subclass per *C. japonica* library sample. **(E)** Separation of cells in mixed-species libraries. Cells (highlighted in red) were selected for analysis based on the number of mapped reads to a particular genome (*C. japonica* or *R. imitator* [Wu, Kalakuntla, Goolsby, *et al.,* in preparation]).

**Fig. S6.**
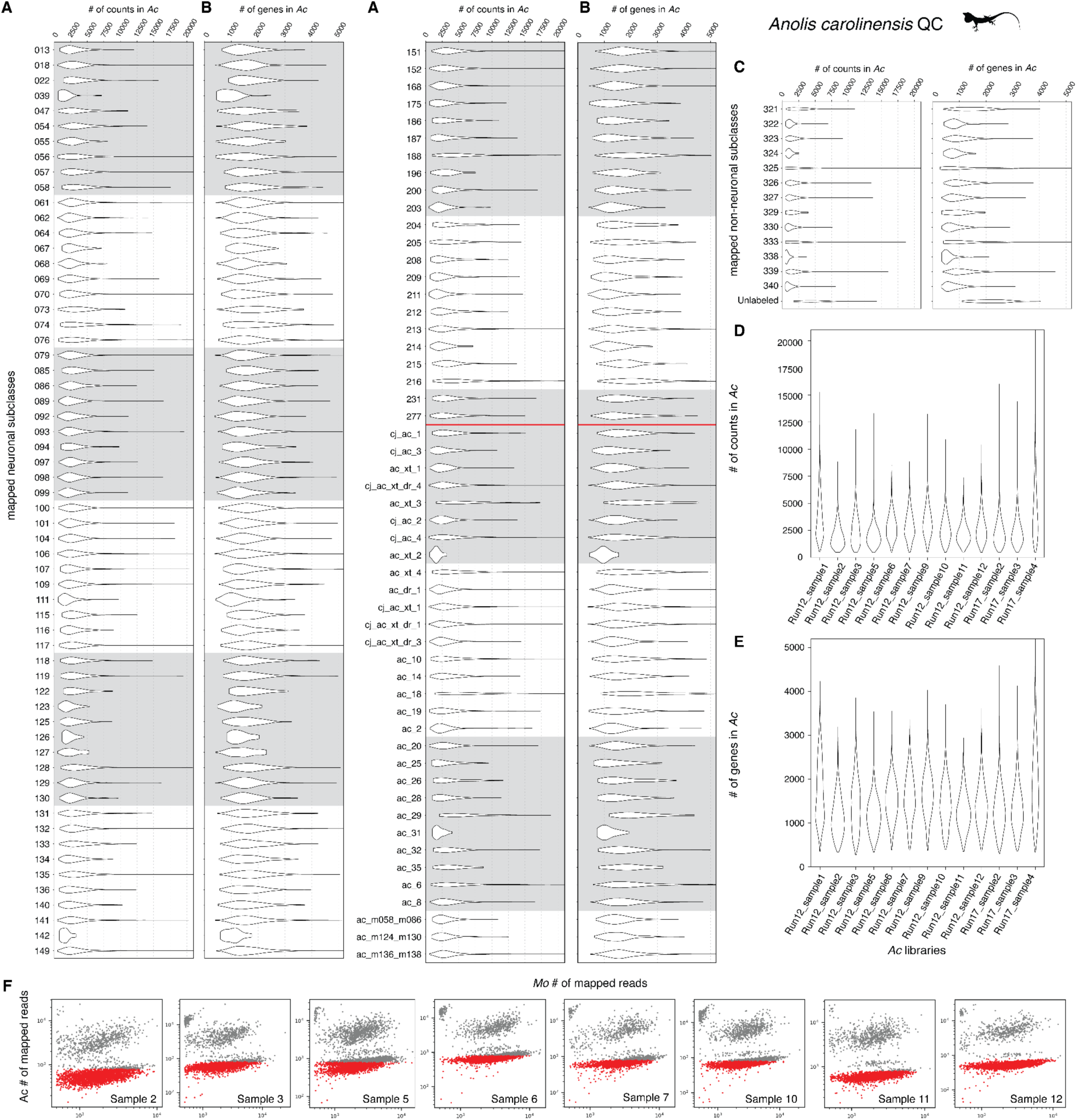
Quality metrics of snRNAseq data for green anole (*Anolis carolinensis*). **(A)** Number of counts per subclass(*21*). This list is larger than the set shown in **Fig. 2**, as it includes the hypothalamus neuronal subclasses, non-hypothalamic neuronal subclasses, and non-neuron subclasses. All neuronal subclasses following the red horizontal line are subclasses not found in mice. **(B)** Number of genes per subclass. For (A) and (B), the top section represents neuronal subclasses, and the bottom section represents non-neuronal subclasses. **(C)** Number of counts/subclass per *A. carolinensis* library sample. **(D)** Number of genes/subclass per *A. carolinensis* library sample. **(E)** Separation of cells in mixed-species libraries. Cells (highlighted in red) were selected for analysis based on the number of mapped reads to a particular genome (*M. ochrogaster* or *A. carolinensis*).

**Fig. S7.**
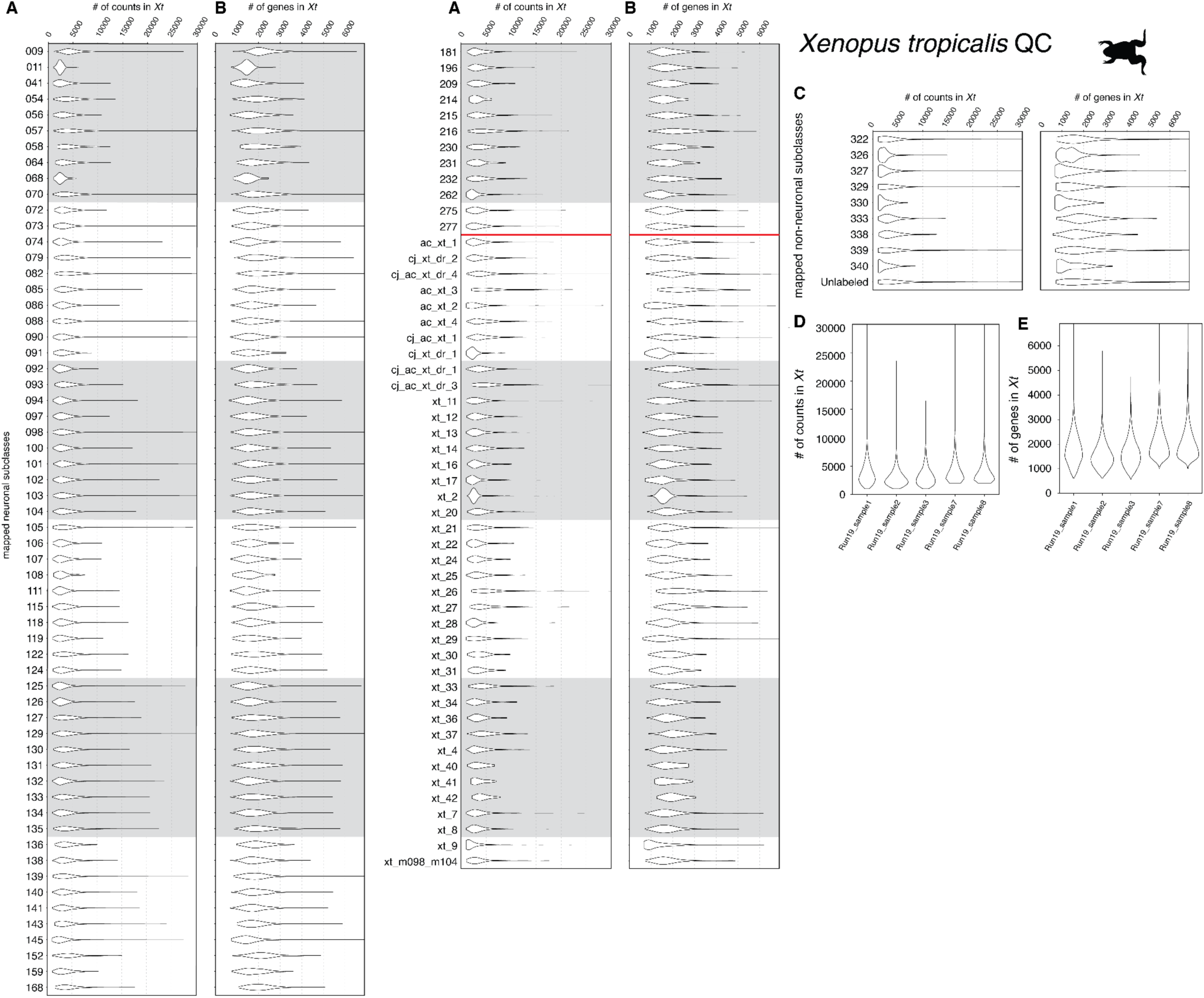
Quality metrics of snRNAseq data for tropical clawed frog (*Xenopus tropicalis*). **(A)** Number of counts per subclass(*21*). This list includes both the hypothalamus neuronal subclasses, non-hypothalamic neuronal subclasses, and non-neuronal subclasses. All neuronal subclasses following the red horizontal line are subclasses not found in mice. **(B)** Number of genes per subclass. For (A) and (B), the top section of rows represents neuronal subclasses, and the bottom section of rows represents non-neuronal subclasses. **(C)** Number of counts/subclass per *X. tropicalis* library sample. **(D)** Number of genes/subclass per *X. tropicalis* library sample.

**Fig. S8.**
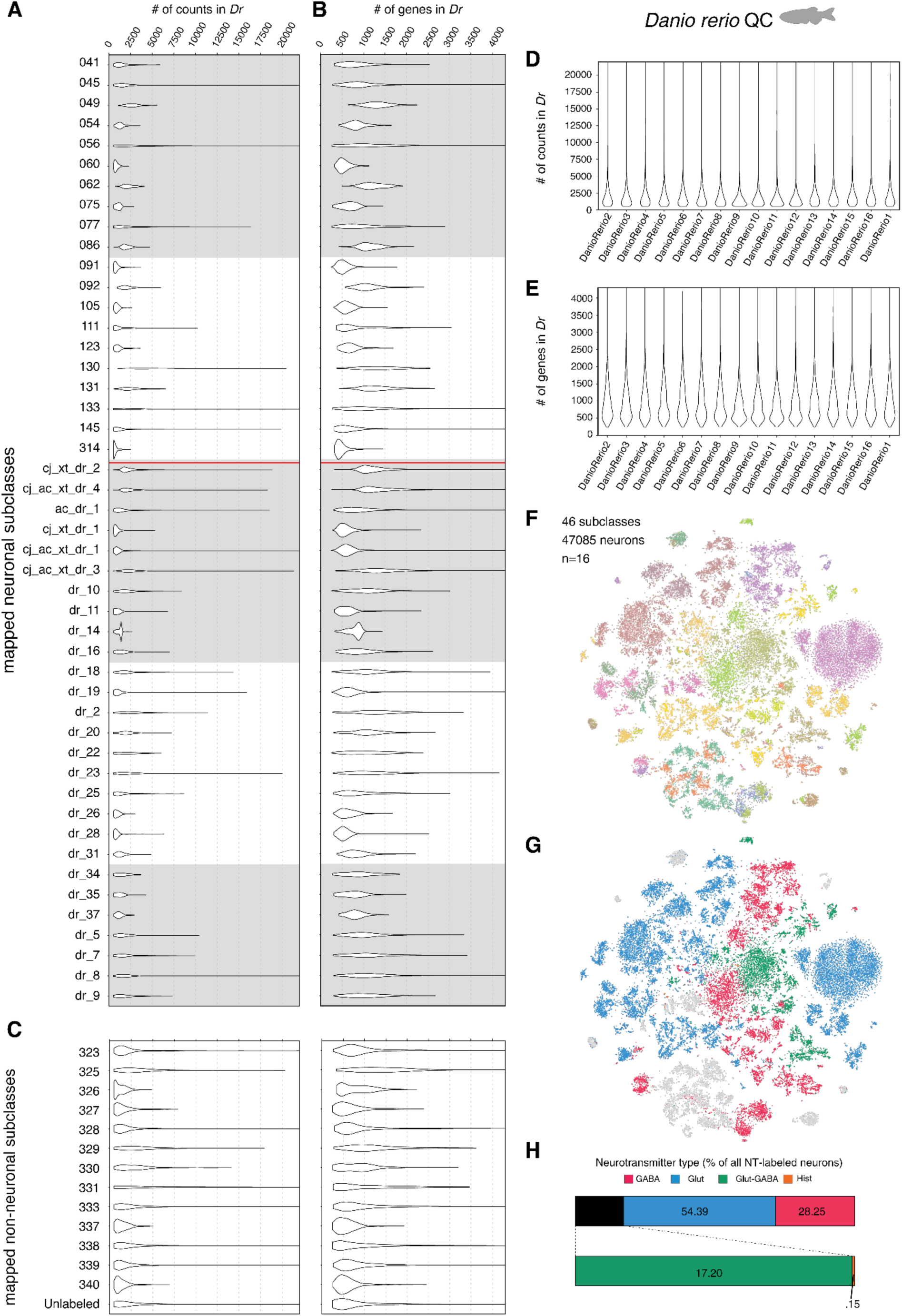
Quality metrics of snRNAseq data for zebrafish (*Danio rerio*). **(A)** Number of counts per subclass. **(B)** Number of genes per subclass. For (A) and (B), the top section represents neuronal subclasses, and the bottom section represents non-neuronal subclasses. All neuronal subclasses following the red horizontal line are subclasses not found in mice. **(C)** Number of counts/subclass per *D. rerio* library sample. **(D)** Number of genes/subclass per *D. rerio* library sample. **(E)** tSNE plot of neuronal transcriptome. Cells are color-coded based on subclasses. Cells sharing the same color across species are part of homologous subclasses (**fig. S9** and **Figs. 1, 2**). Number of sequenced neurons that pass quality control, subclasses, and number of animals used are listed to the left. **(F)** tSNE plot with colors indicating neurotransmitter usage. The thresholds for neurotransmitter classification can be found in **Methods.** Below, proportion of all neurotransmitters (top) and the proportion among non-glutamatergic/GABAergic neurotransmitters (bottom). Neurons with unclear neurotransmitter expression were excluded from this analysis.

**Fig. S9.**
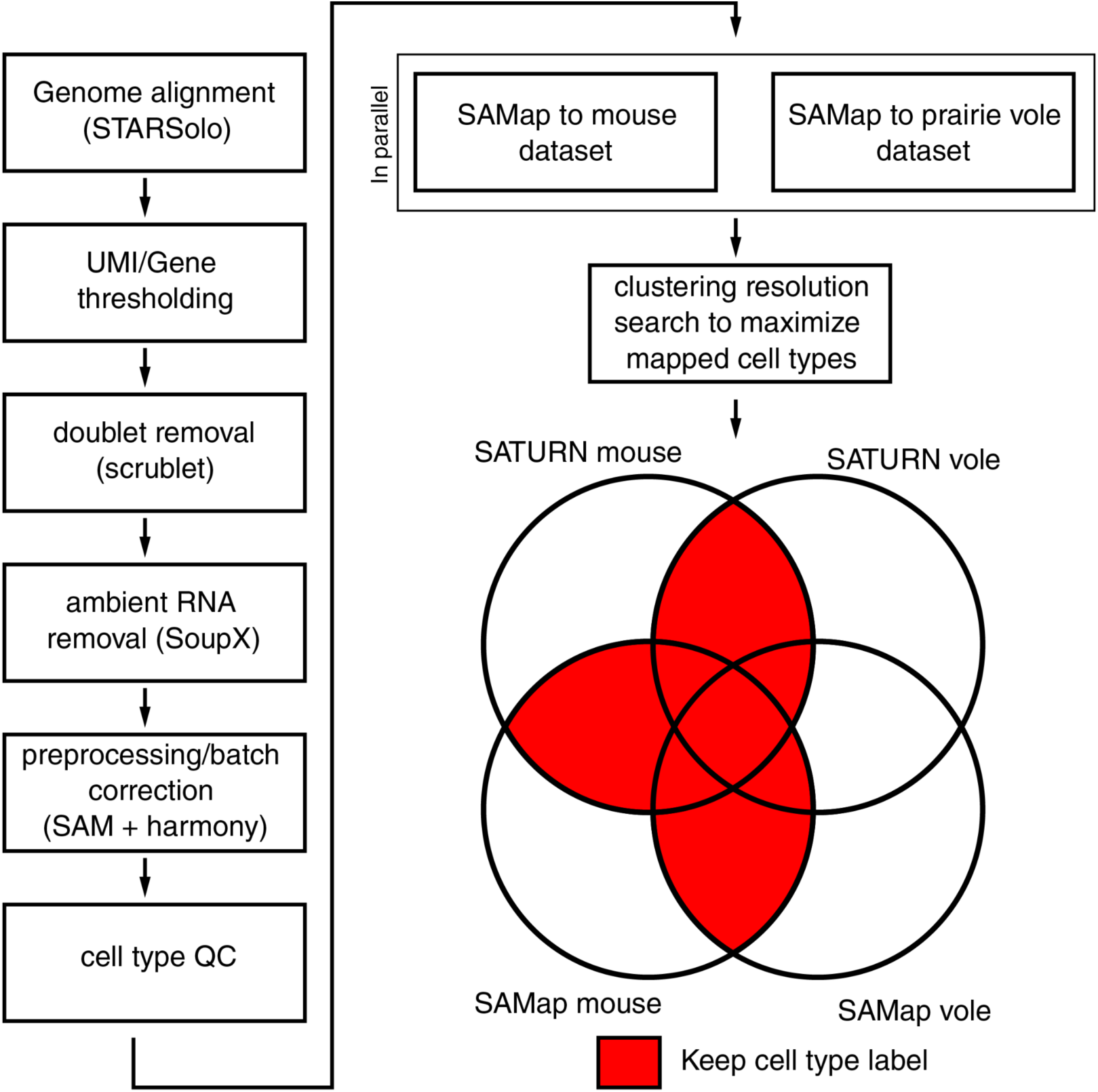
Data processing and mapping in quail, anole, *Xenopus*, and zebrafish. Raw snRNAseq data is aligned to the genome using STARsolo, a function within STAR (*155*) which is then passed through UMI/gene thresholding to remove debris and empty doublets. Then, we used Scrublet (*115*) to detect and remove doublets (co-encapsulated cells). Data were then passed through ambient RNA removal through SoupX (*124*) to remove cell-free RNA contamination. Cleaned data were then passed through SAM (*116*) and Harmony (*117*) for preprocessing and batch correction, after which the mapped subclasses were evaluated for quality. The remaining cells were then passed through SAMap (*28*) to map to the mouse and prairie vole datasets in parallel. In parallel, cells were mapped using SATURN (*29*). Datasets were compared between SAMapped mouse labels, SATURN-mapped mouse labels, SAMapped vole labels, and SATURN-mapped vole labels. If a cell type overlapped between one of the two datasets or more (the one exception being if a label only overlapped in SATURN-mapped vole and SAMapped vole, but no mouse datasets), the mapped cell type label was kept.

**Fig. S10.**
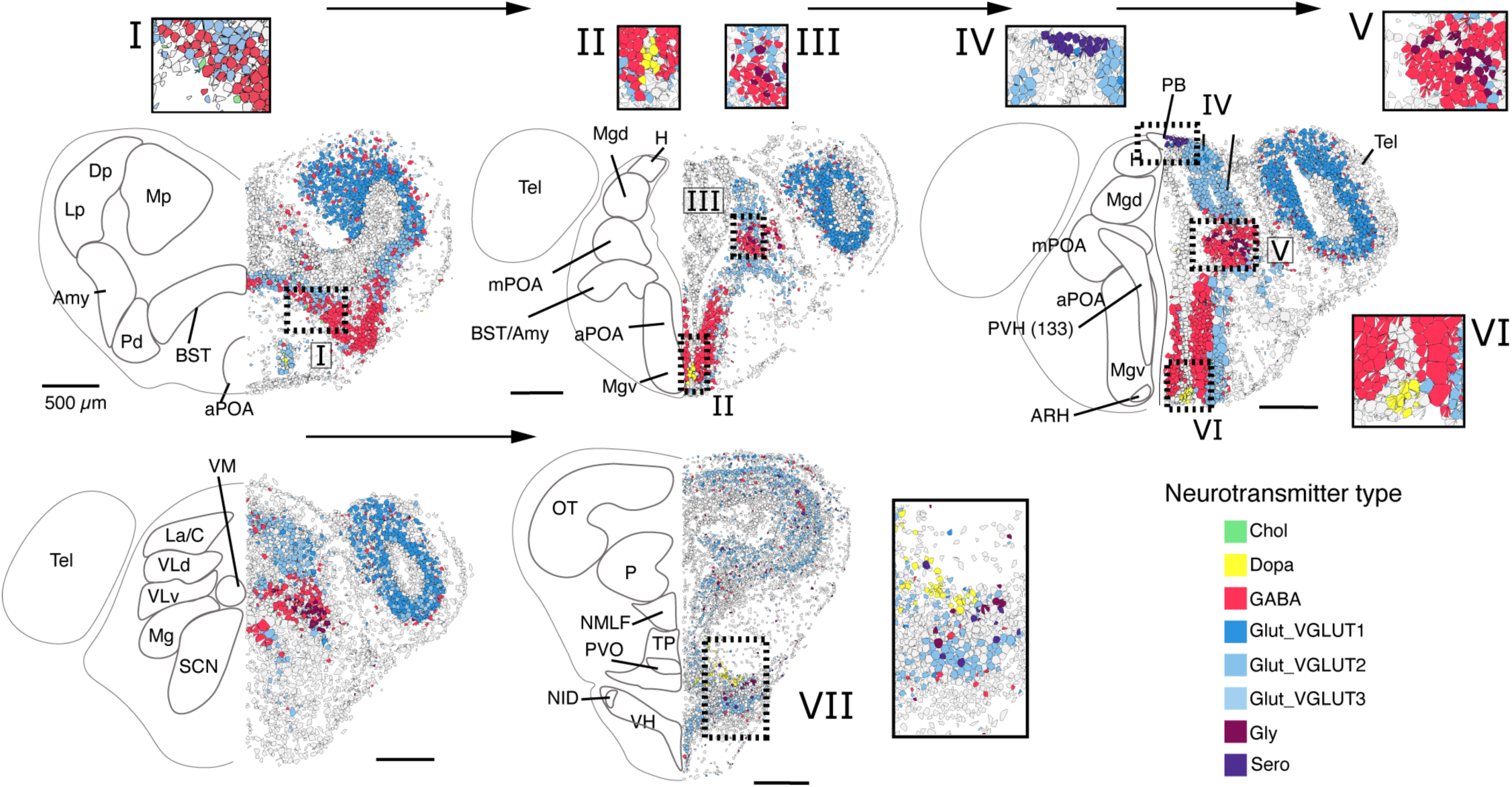
Spatial distribution of neurotransmitter types in the hypothalamus of *Xenopus tropicalis*. Serial coronal sections with markers for neurotransmitter types visualized through STARmap. Horizontal arrows point from the anterior-to-posterior direction. Insets are magnified from the dashed boxes. Note that we revise the locations of some hypothalamic nuclei in the amphibian reference atlas, which is a modest departure from the prior published atlas (*60*). The additional atlases were referenced: (*148*–*154*). Notably, we specify a location for the paraventricular hypothalamus (PVH) which was driven by studies on subclass 133 (see **Fig. 4** for detail). We additionally identified proposed locations the nucleus infundibularis dorsalis (NID) and arcuate nucleus (ARH), respectively. The paraventricular organ is specified based on expression of serotonin and location along the fourth ventricle, in agreement with where this nucleus was originally proposed to be (*149*, *152*). **Abbreviations:** Amy, amygdala; aPOA, anterior preoptic area; ARH, arcuate nucleus; BST, bed nucleus stria terminalis; C, central thalamic nucleus; Dp, dorsal pallium; H, habenula; La, lateral thalamic nucleus; Lp, lateral pallium; Mg, magnocellular preoptic nucleus; Mgd, magnocellular preoptic nucleus, dorsal; Mgv, magnocellular preoptic nucleus, ventral; Mp, medial pallium; mPOA, medial preoptic area; NMLF, nucleus of the medial longitudinal fasciculus; OT, optic tectum; P, posterior thalamic nucleus; PB, pineal body; Pd, pallidum; PVH, paraventricular hypothalamus – equivalent to subclass 133; PVO, paraventricular organ; SCN, suprachiasmatic nucleus; Tel, telencephalon; TP, posterior tuberculum; VLd, ventrolateral thalamic nucleus, dorsal; VLv, ventrolateral thalamic nucleus, ventral; VM, ventromedial thalamic nucleus; VH, ventral hypothalamic nucleus.

**Fig. S11.**
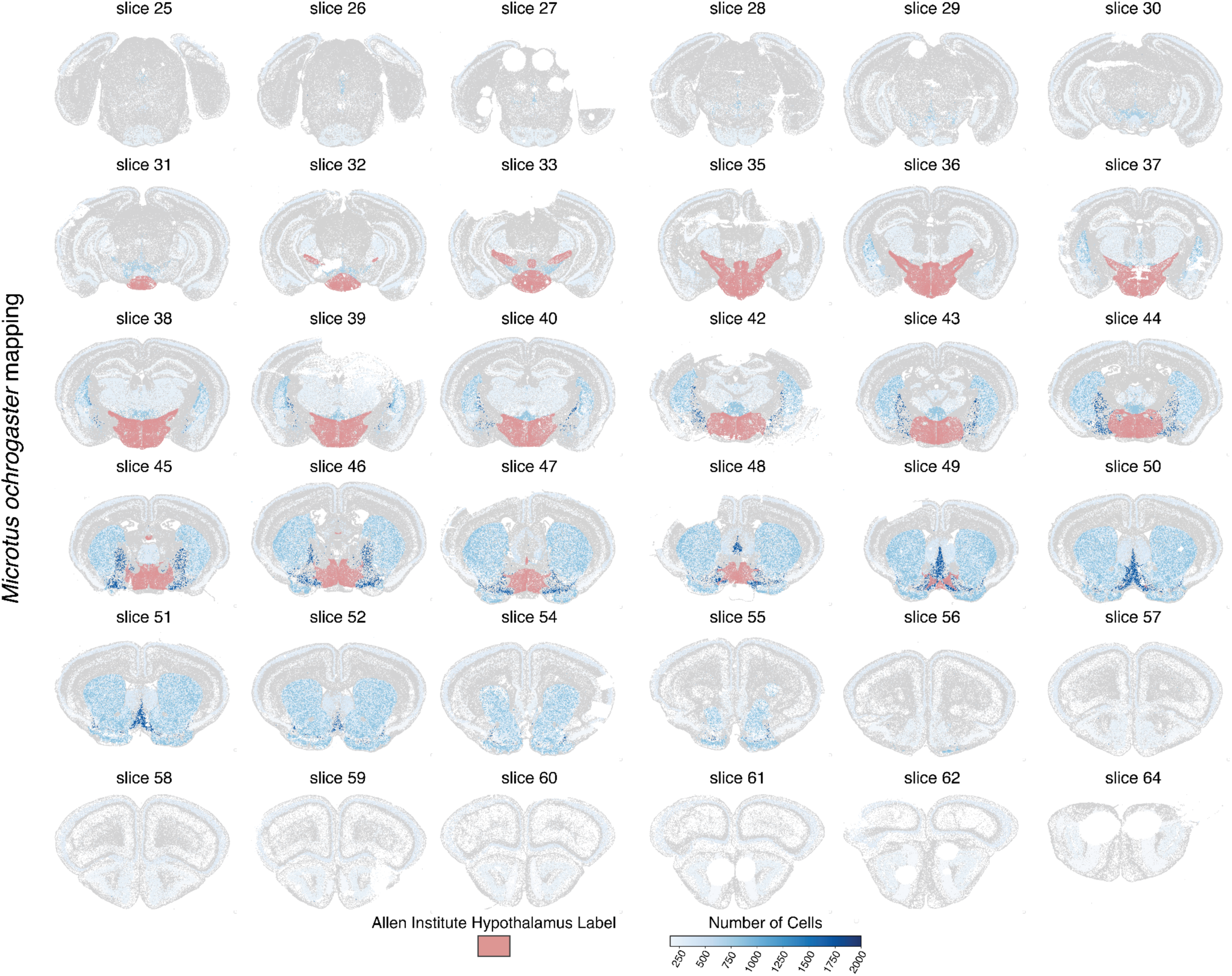
Mapping of cell types outside the mouse hypothalamus in *Microtus ochrogaster.* We mapped subclasses bordering the mouse hypothalamus that were present in our *M. ochrogaster* data. This was visualized onto MERFISH slice 25-64 from Allen Brain Atlas MERFISH-C57BL6J-638850 common coordinate framework (CCFv3) (*21*, *129*). Slices 34, 41, 53, and 63 are missing from the original dataset. Blue coloration scales with cell density. Red coloration is the tracing of the Allen Institute’s Hypothalamus label from CCFv3. Mappings of cells are relative to an entire brain region rather than just the representative slice shown. Anatomical regions outside of the hypothalamus are defined based on the Allen Institute and their subclasses. 86% of neurons in vole mapped to the mouse. Of those neurons, 83% mapped in the hypothalamus.

**Fig. S12.**
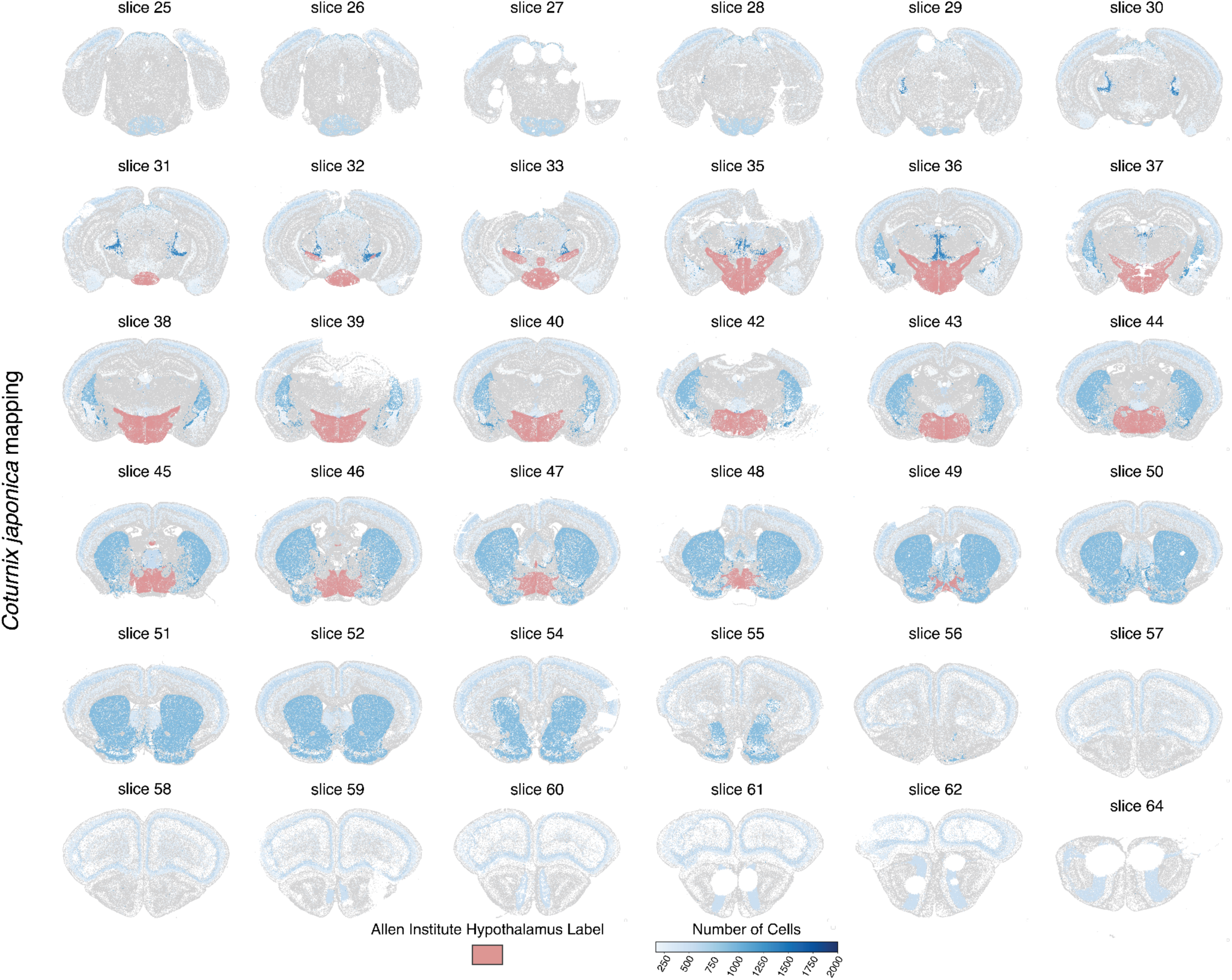
Mapping of cell types outside the mouse hypothalamus in *Coturnix japonica.* We mapped subclasses bordering the mouse hypothalamus that were present in the quail data. This was visualized onto MERFISH slice 25-64 from Allen Brain Atlas MERFISH-C57BL6J-638850 common coordinate framework (CCFv3) (*21*, *129*). Slices 34, 41, 53, and 63 are missing from the original dataset. Blue coloration scales with cell density. Red coloration is the tracing of the Allen Institute’s Hypothalamus label from CCFv3. Mappings of cells are relative to an entire brain region rather than just the representative slice shown. Anatomical regions outside of the hypothalamus are defined based on the Allen Institute and their subclasses. 80% of neurons in the quail mapped to the mouse. Of those neurons, 68% mapped to the hypothalamus.

**Fig. S13.**
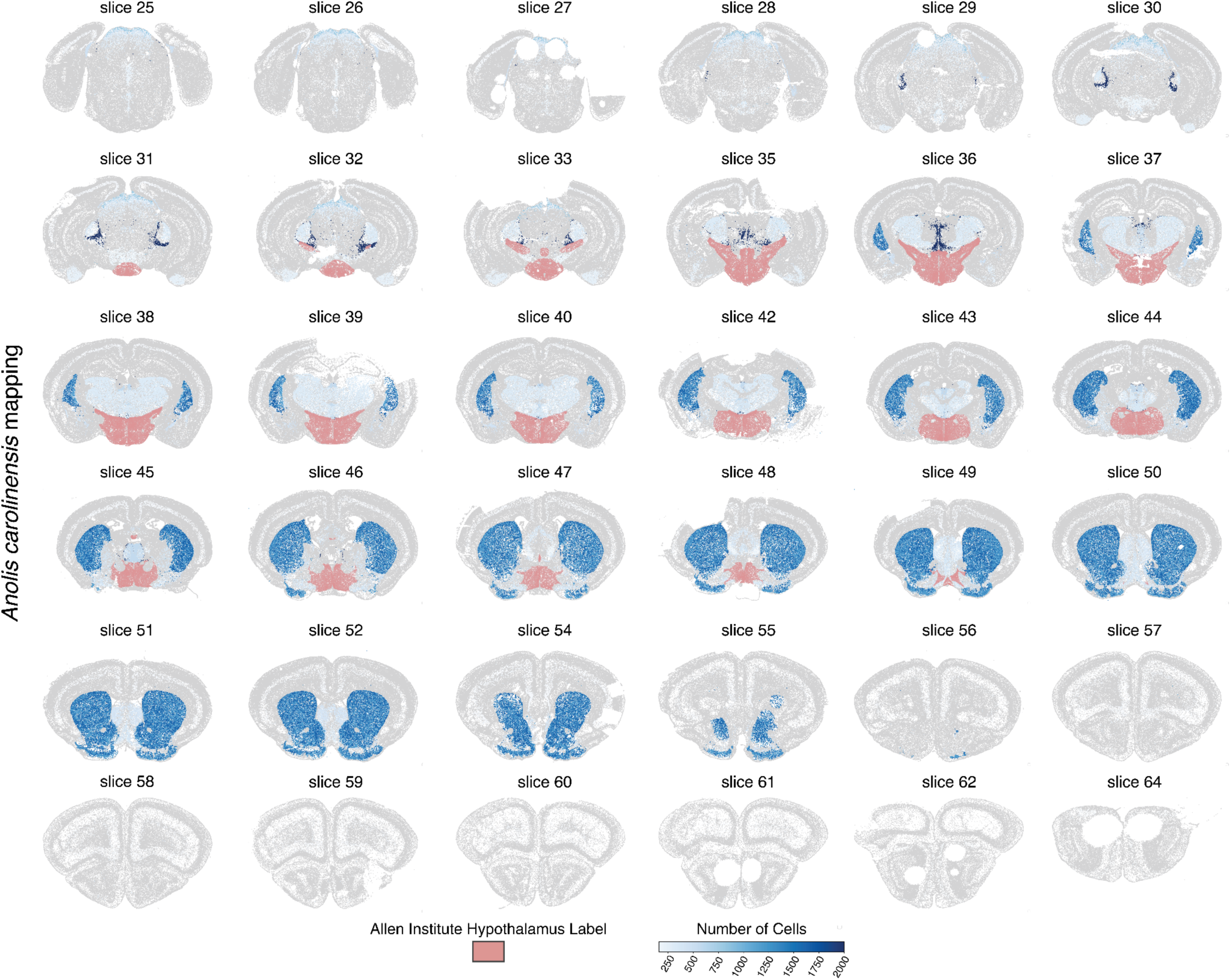
Mapping of cell types outside the mouse hypothalamus in *Anolis carolinensis.* We mapped subclasses bordering the mouse hypothalamus present in the anole data. This was visualized onto MERFISH slice 25-64 from Allen Brain Atlas MERFISH-C57BL6J-638850 common coordinate framework (CCFv3) (*21*, *129*). Slices 34, 41, 53, and 63 are missing from the original dataset. Blue coloration scales with cell density. Red coloration is the tracing of the Allen Institute’s Hypothalamus label from CCFv3. Mappings of cells are relative to an entire brain region rather than just the representative slice shown. Anatomical regions outside of the hypothalamus are defined based on the Allen Institute and their subclasses. 69% of neurons in the anole mapped to the mouse. Of those neurons, 49% mapped to the hypothalamus.

**Fig. S14.**
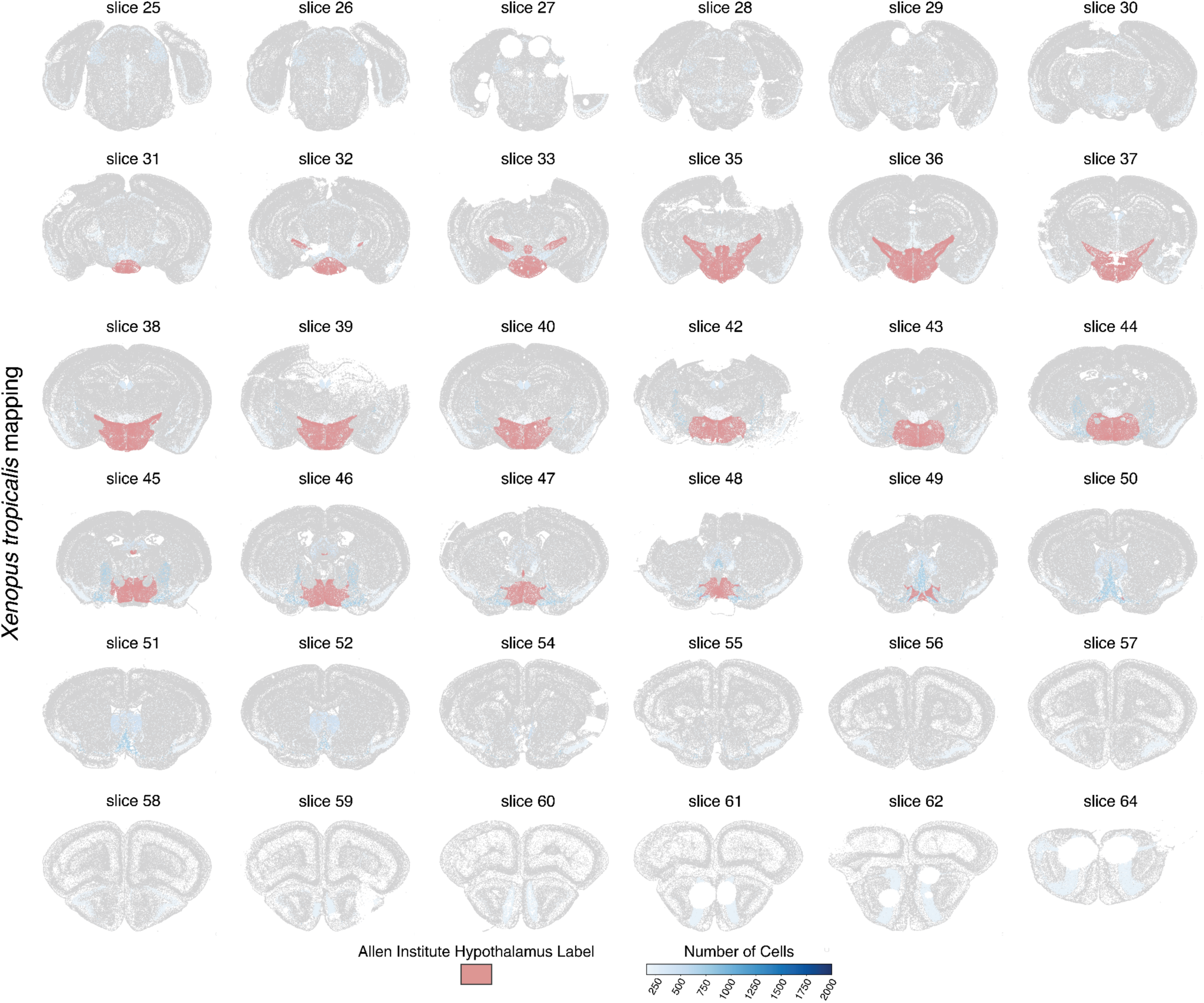
Mapping of cell types outside the mouse hypothalamus in *Xenopus tropicalis.* We mapped subclasses bordering the mouse hypothalamus present in the *Xenopus* data. This was visualized onto MERFISH slice 25-64 from Allen Brain Atlas MERFISH-C57BL6J-638850 common coordinate framework (CCFv3) (*21*, *129*). Slices 34, 41, 53, and 63 are missing from the original dataset. Blue coloration scales with cell density. Red coloration is the tracing of the Allen Institute’s Hypothalamus label from CCFv3. Mappings of cells are relative to an entire brain region rather than just the representative slice shown. Anatomical regions outside of the hypothalamus are defined based on the Allen Institute and their subclasses. 72% of neurons in the *Xenopus* mapped to the mouse. Of those neurons, 81% mapped to the hypothalamus.

**Fig. S15.**
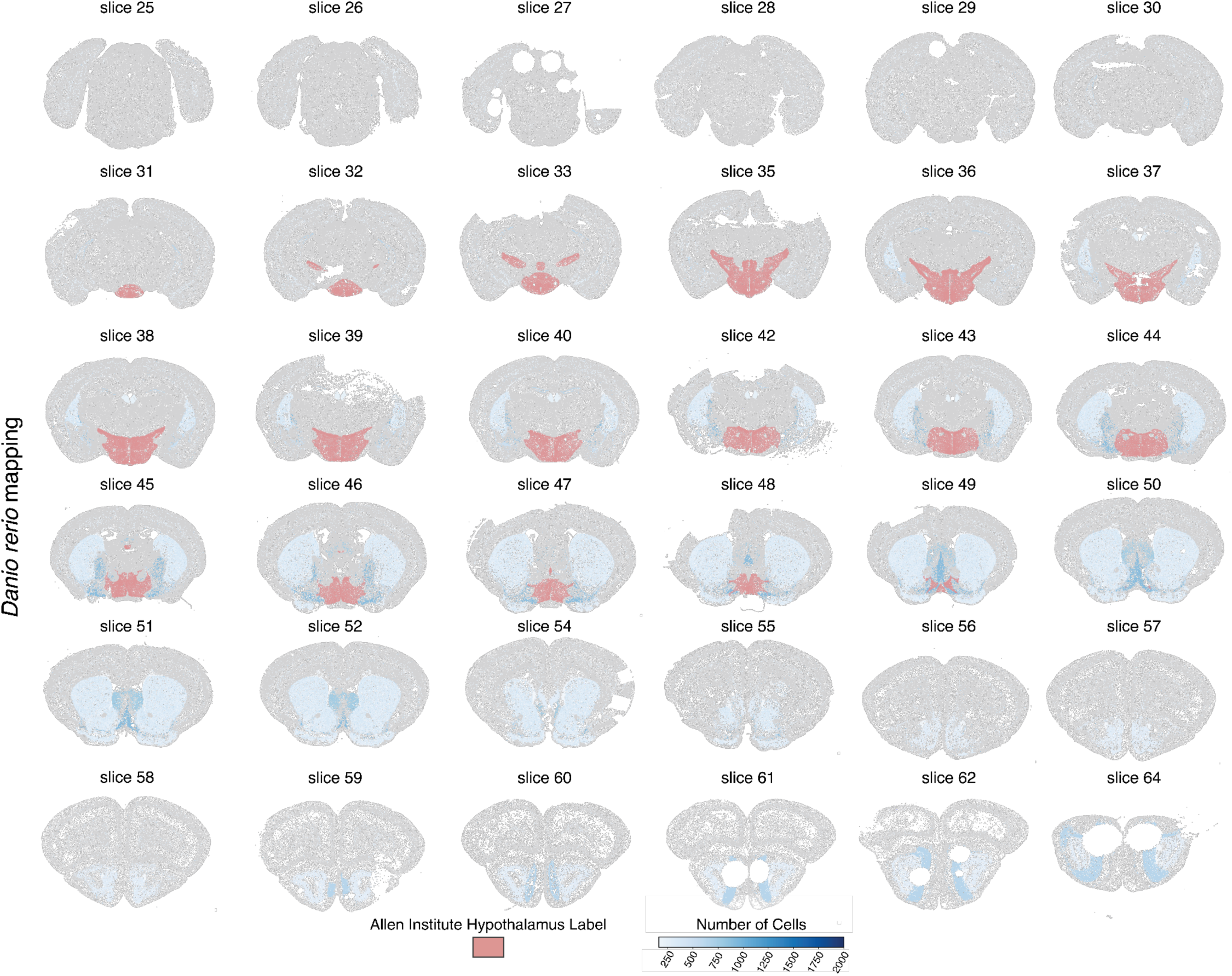
Mapping of cell types outside the mouse hypothalamus in *Danio rerio.* We mapped subclasses bordering the mouse hypothalamus present in the zebrafish data. This was visualized onto MERFISH slice 25-64 from Allen Brain Atlas MERFISH-C57BL6J-638850 common coordinate framework (CCFv3) (*21*, *129*). Slices 34, 41, 53, and 63 are missing from the original dataset. Blue coloration scales with cell density. Red coloration is the tracing of the Allen Institute’s Hypothalamus label from CCFv3. Mappings of cells are relative to an entire brain region rather than just the representative slice shown. Anatomical regions outside of the hypothalamus are defined based on the Allen Institute and their subclasses. 15% of neurons in the zebrafish mapped to the mouse. Of those neurons, 55% mapped to the hypothalamus.

**Fig. S16.**
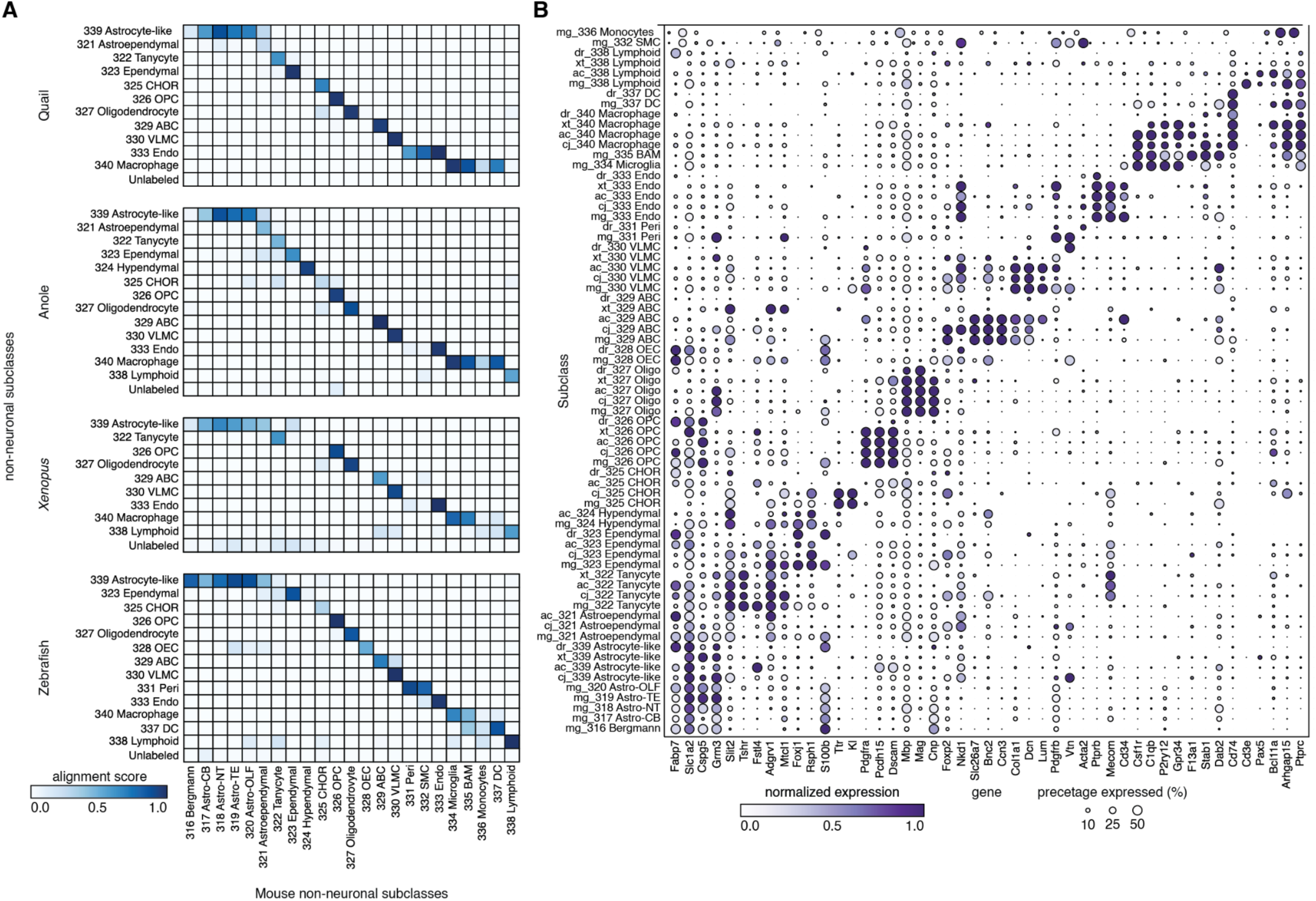
Identification of non-neuronal subclasses and their marker genes. **(A)** SAMap alignment scores between non-neuronal subclasses in mouse and non-neuronal subclasses in quail, anole, *Xenopus*, and zebrafish. 316, 317, 318, 319, and 320 in mouse were combined for astrocytes in **Fig. 2F**, 334 and 335 in mouse were combined for BAM/microglia in **Fig. 2F**. (**B**) Identification of species-specific non-neuronal subclasses and the average expression of their marker genes. For each gene, mean expression was computed within each subclass of a given species, and the resulting per-subclass values were min–max scaled by subtracting the minimum and dividing by the range (maximum minus minimum) across subclasses.

**Fig. S17.**
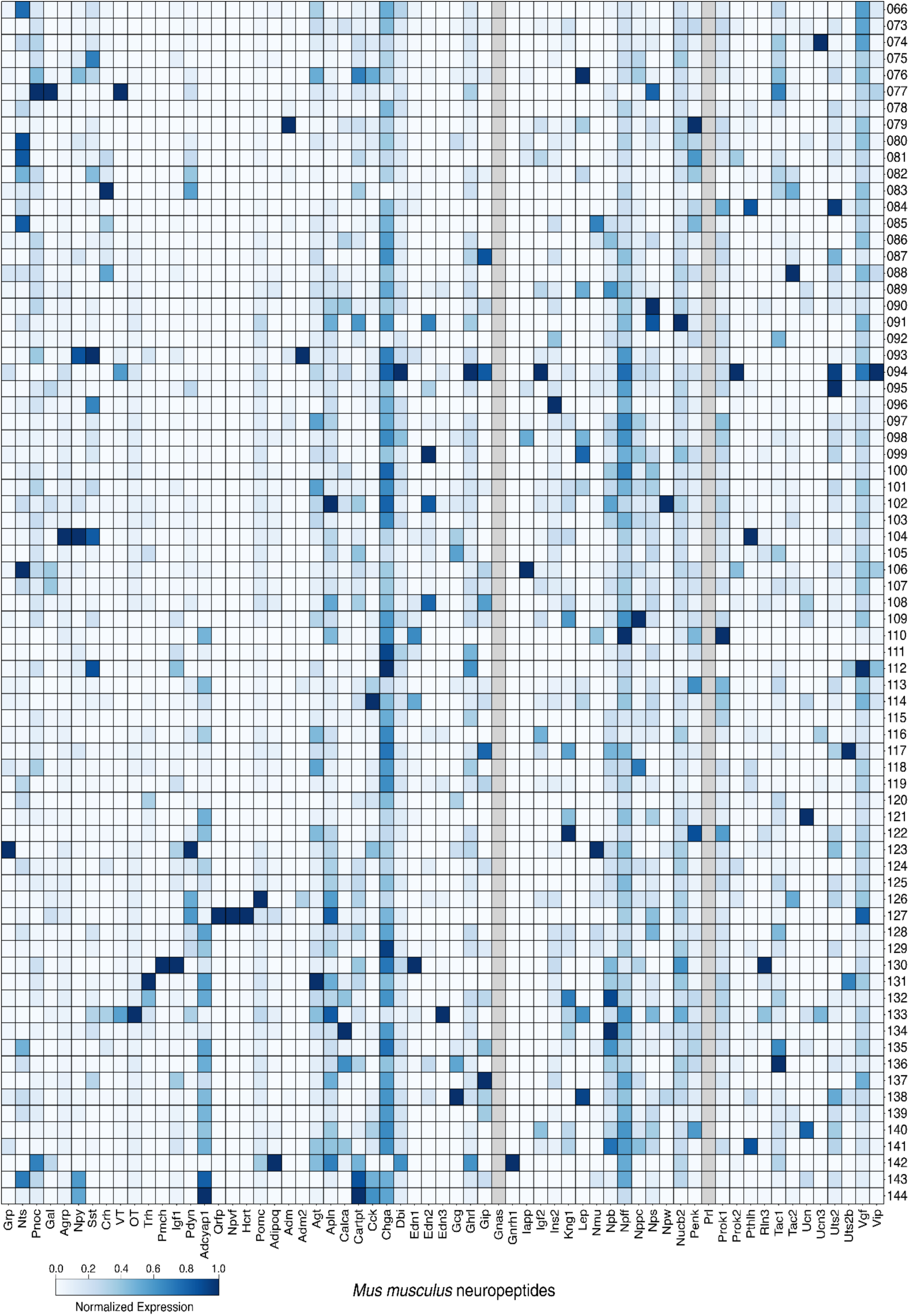
Neuropeptide expression in *Mus musculus* subclasses. Expression of neuropeptides with orthologs in 4 of 6 species in mapped subclasses. Bottom, names of selected neuropeptides. Right, IDs for all mapped hypothalamic subclasses. Color represents min-max normalized expression across hypothalamic subclasses in the corresponding subclass (scale at the bottom). Grey indicates that the gene is either not expressed, or no ortholog is found in the genome.

**Fig. S18.**
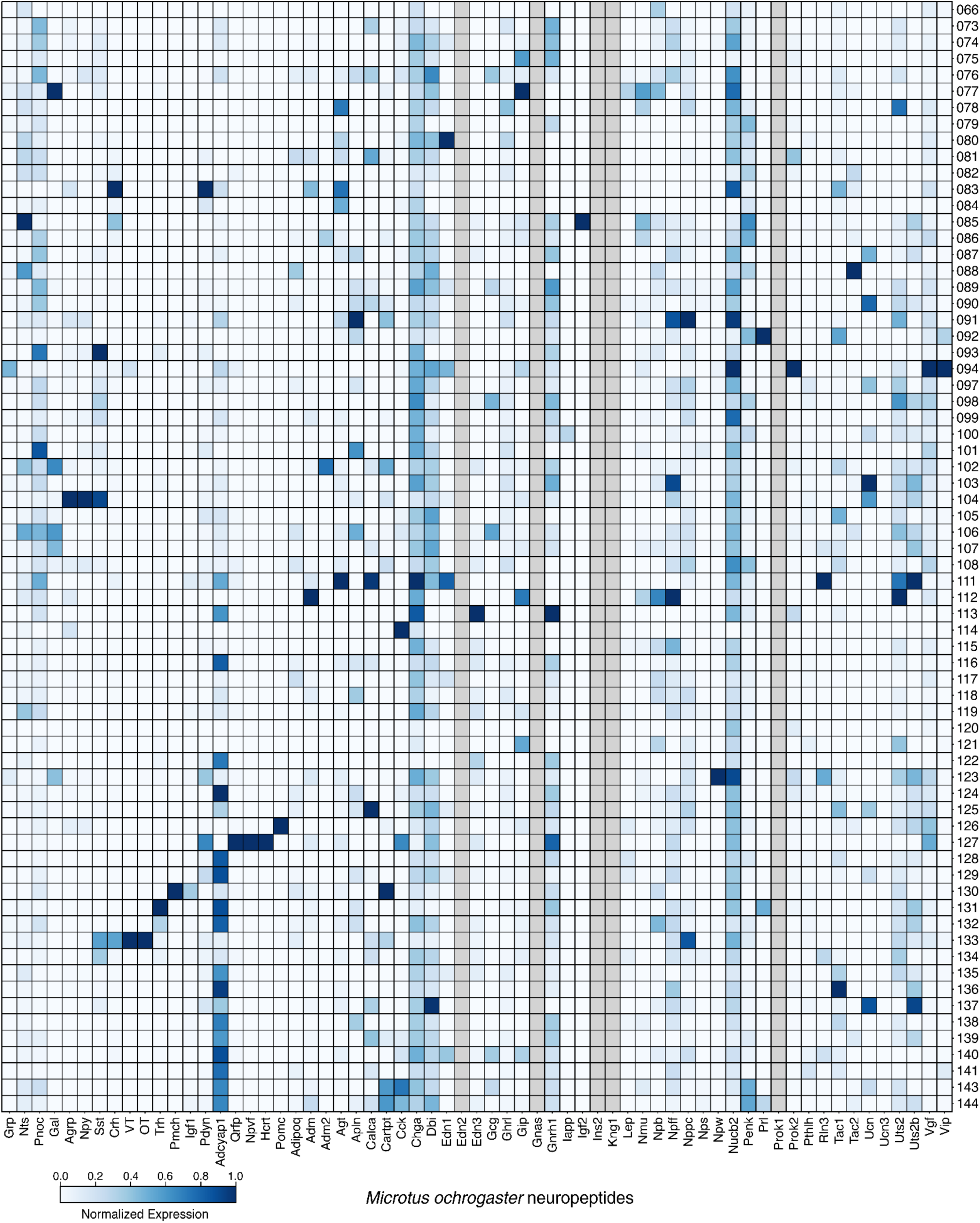
Neuropeptide expression in *Microtus ochrogaster* subclasses. Expression of neuropeptides with orthologs in 4 of 6 species in mapped subclasses. Bottom, names of selected neuropeptides. Right, IDs for all mapped hypothalamic subclasses. Color represents min-max normalized expression across hypothalamic subclasses in the corresponding subclass (scale at the bottom). Grey indicates that the gene is either not expressed, or no ortholog is found in the genome.

**Fig. S19.**
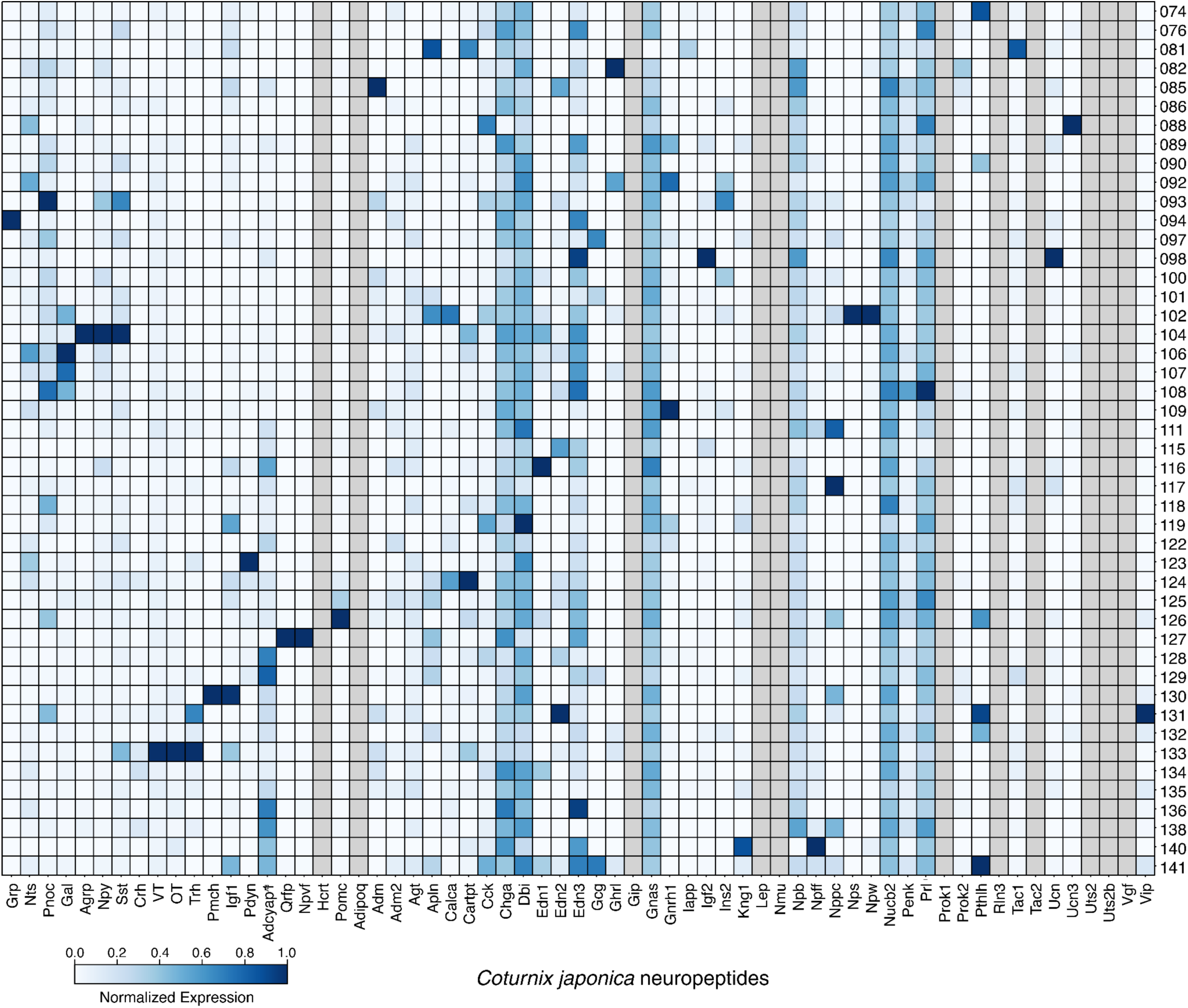
Neuropeptide expression in *Coturnix japonica* subclasses. Expression of neuropeptides with orthologs in 4 of 6 species in mapped subclasses. Bottom, names of selected neuropeptides. Right, IDs for all mapped hypothalamic subclasses. Color represents min-max normalized expression across hypothalamic subclasses in the corresponding subclass (scale at the left). Grey indicates that the gene is either not expressed, or no ortholog is found in the genome.

**Fig. S20.**
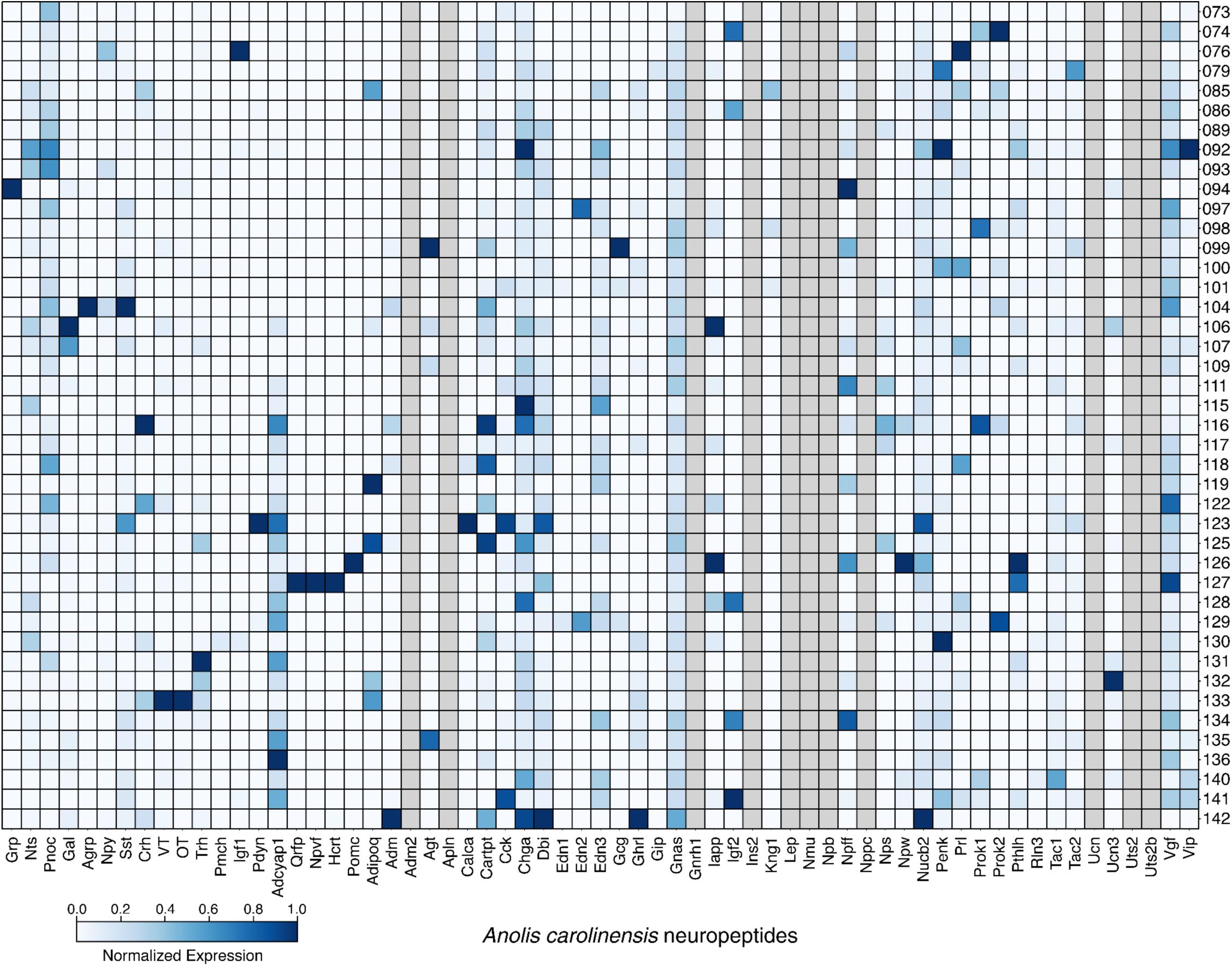
Neuropeptide expression in *Anolis carolinensis* subclasses. Expression of neuropeptides with orthologs in 4 of 6 species in mapped subclasses. Bottom, names of selected neuropeptides. Right, IDs for all mapped hypothalamic subclasses. Color represents min-max normalized expression across hypothalamic subclasses in the corresponding subclass (scale at the left). Grey indicates that the gene is either not expressed, or no ortholog is found in the genome.

**Fig. S21.**
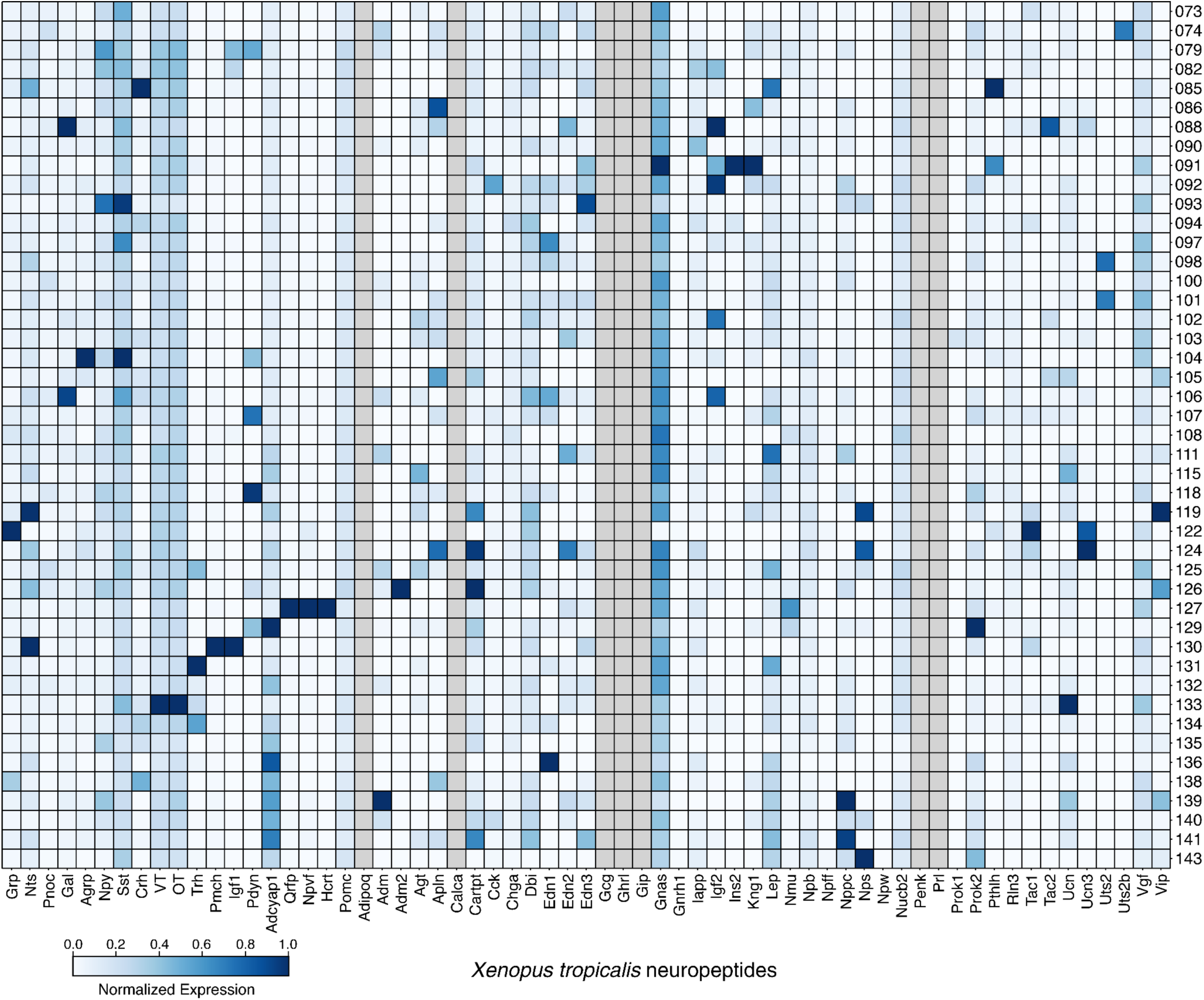
Neuropeptide expression in *Xenopus tropicalis* subclasses. Expression of neuropeptides with orthologs in 4 of 6 species in mapped subclasses. Bottom, names of selected neuropeptides. Right, IDs for all mapped hypothalamic subclasses. Color represents min-max normalized expression across hypothalamic subclasses in the corresponding subclass (scale at the left). Grey indicates that the gene is either not expressed, or no ortholog is found in the genome.

**Fig. S22.**
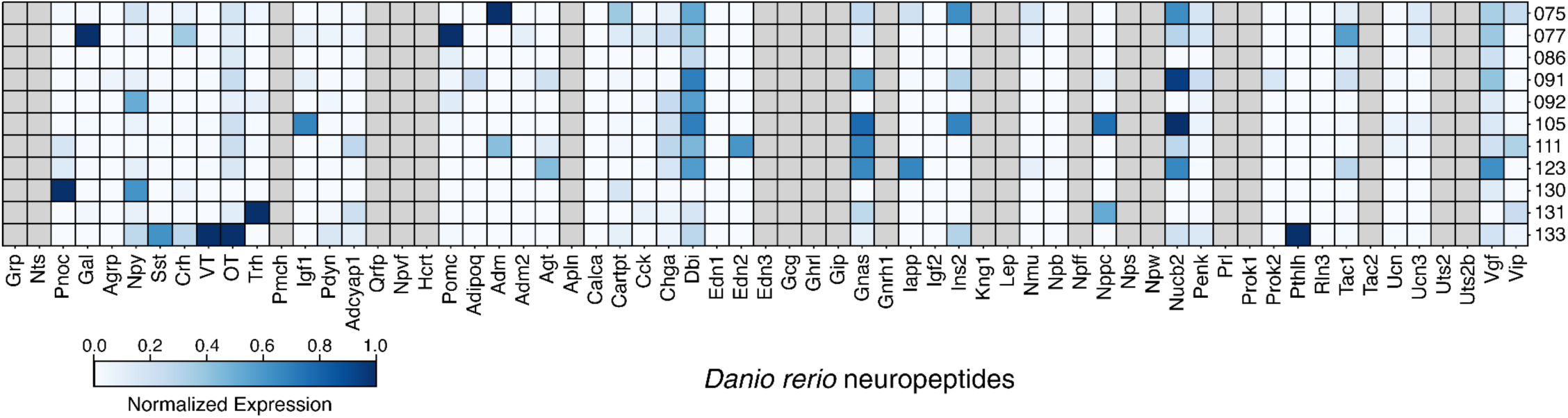
Neuropeptide expression in *Danio rerio* subclasses. Expression of neuropeptides with orthologs in 4 of 6 species in mapped subclasses. Bottom, names of selected neuropeptides. Right, IDs for all mapped hypothalamic subclasses. Color represents min-max normalized expression across hypothalamic subclasses in the corresponding subclass (scale at the left). Grey indicates that the gene is either not expressed, or no ortholog is found in the genome.

**Fig. S23.**
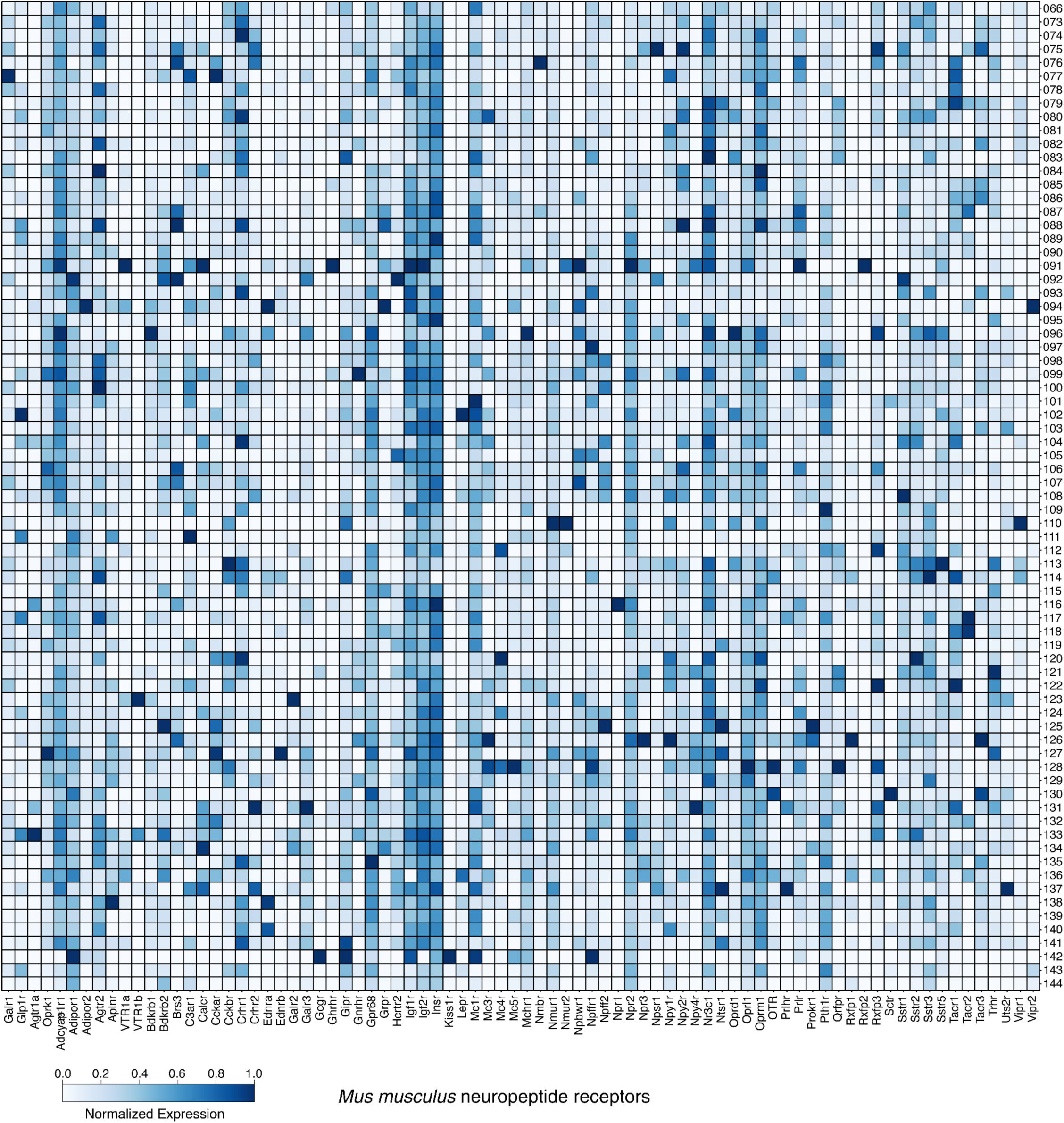
Neuropeptide receptor expression in *Mus musculus* subclasses. Expression of neuropeptide receptors with orthologs in 4 of 6 species in mapped subclasses. Bottom, names of selected neuropeptide receptors. Right, IDs for all mapped hypothalamic subclasses. Color represents min-max normalized expression across hypothalamic subclasses in the corresponding subclass (scale at the left). Grey indicates that the gene is either not expressed with any counts, or no ortholog is found in the genome.

**Fig. S24.**
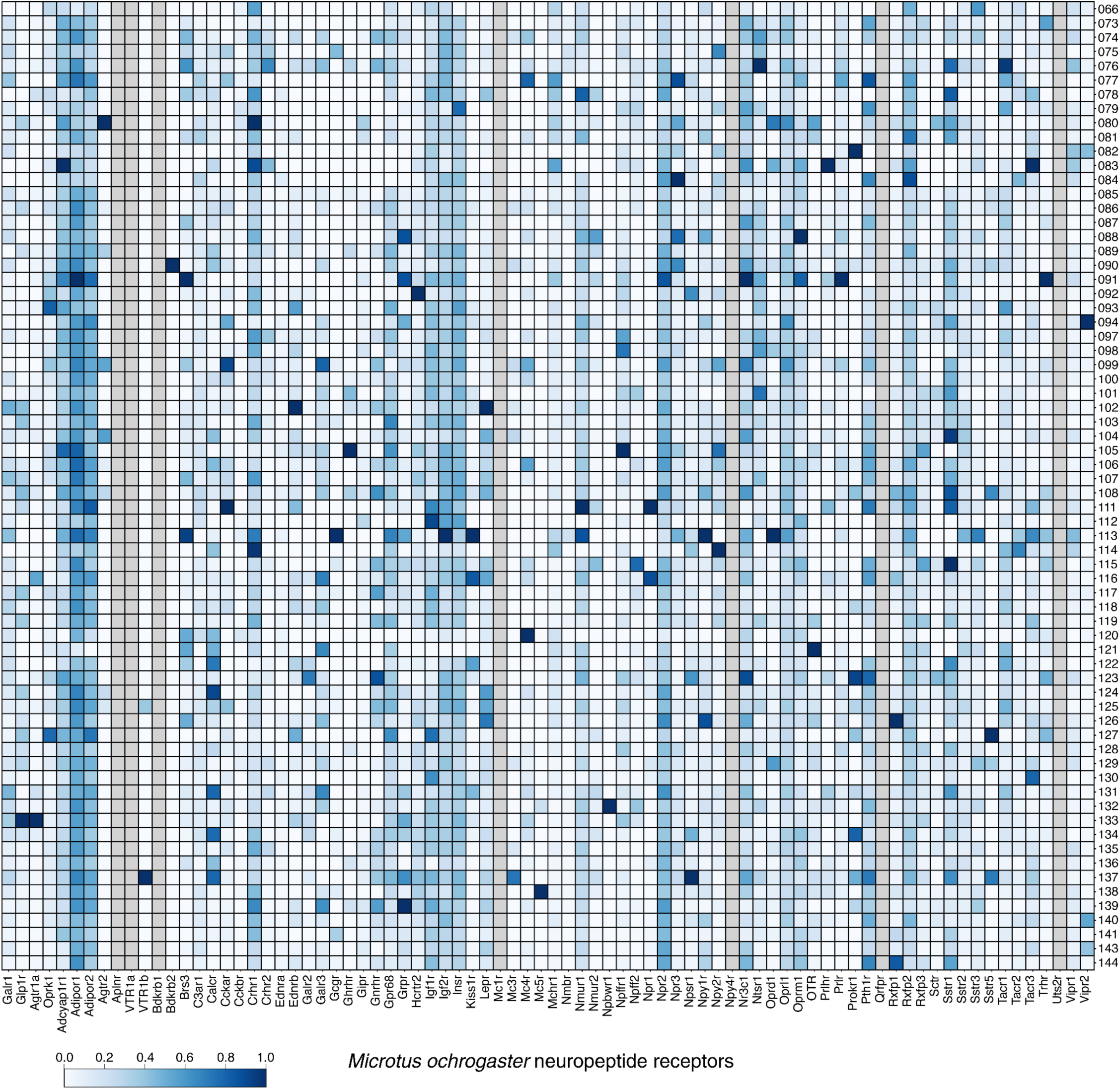
Neuropeptide receptor expression in *Microtus ochrogaster* subclasses. Expression of neuropeptide receptors with orthologs in 4 of 6 species in mapped subclasses. Bottom, names of selected neuropeptide receptors. Right, IDs for all mapped hypothalamic subclasses. Color represents min-max normalized expression across hypothalamic subclasses in the corresponding subclass (scale at the left). Grey indicates that the gene is either not expressed, or no ortholog is found in the genome.

**Fig. S25.**
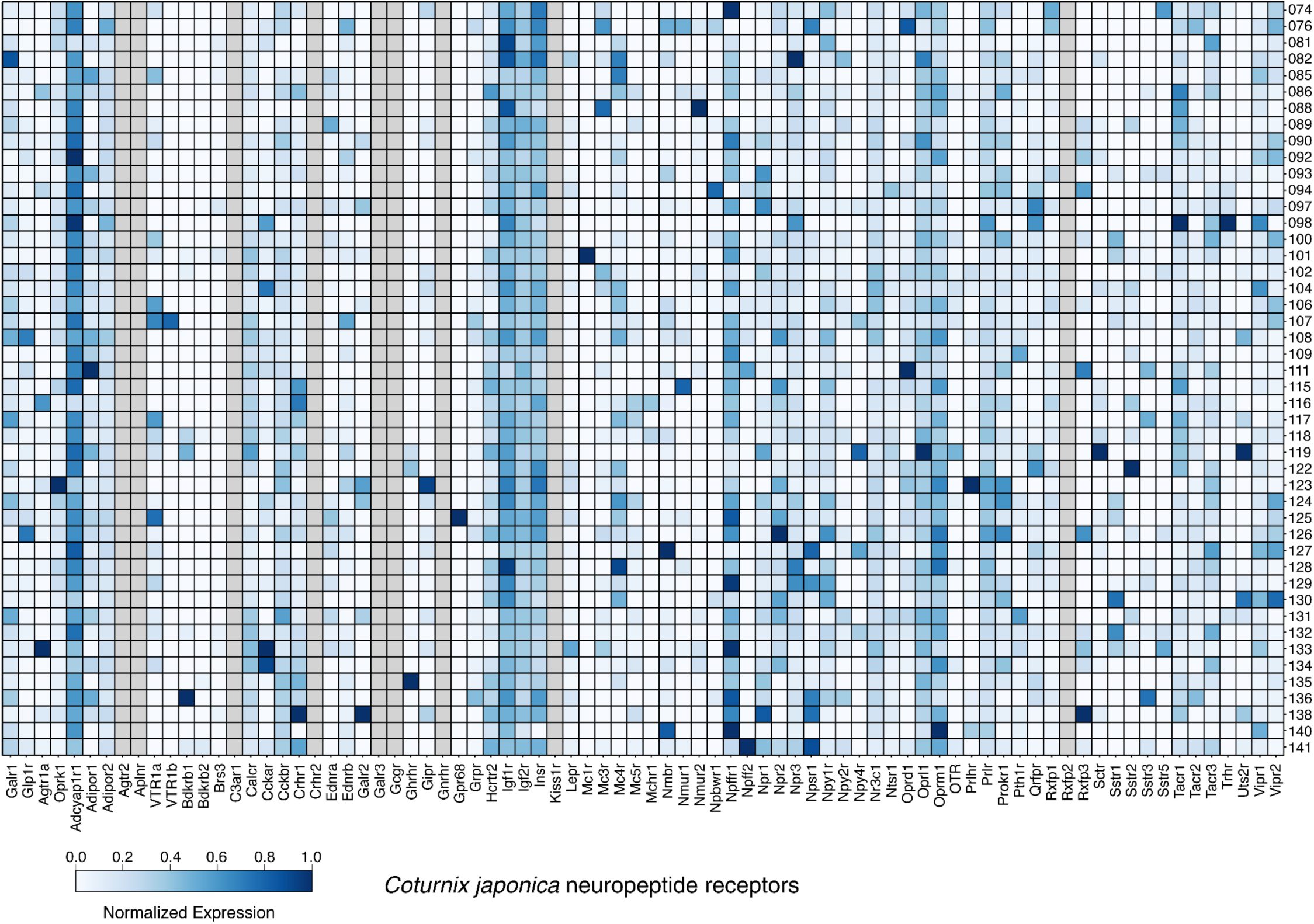
Neuropeptide receptor expression in *Coturnix japonica* subclasses. Expression of neuropeptide receptors with orthologs in 4 of 6 species in mapped subclasses. Bottom, names of selected neuropeptide receptors. Right, IDs for all mapped hypothalamic subclasses. Color represents min-max normalized expression across hypothalamic subclasses in the corresponding cell type (scale at the left). Grey indicates that the gene is either not expressed, or no ortholog is found in the genome.

**Fig. S26.**
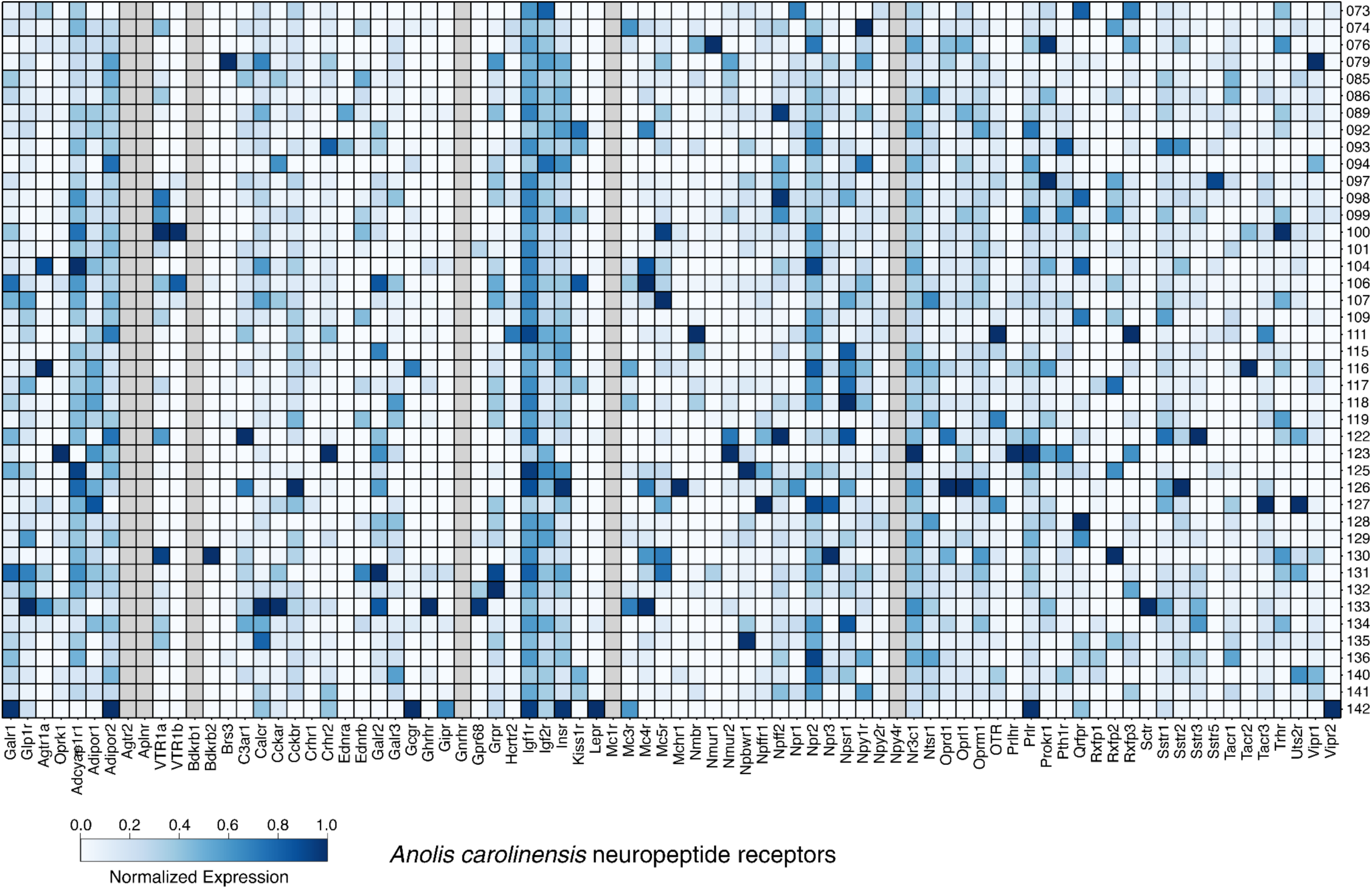
Neuropeptide receptor expression in *Anolis carolinensis* subclasses. Expression of neuropeptide receptors with orthologs in 4 of 6 species in mapped subclasses. Bottom, names of selected neuropeptide receptors. Right, IDs for all mapped hypothalamic subclasses. Color represents min-max normalized expression across hypothalamic subclasses in the corresponding cell type (scale at the left). Grey indicates that the gene is either not expressed, or no ortholog is found in the genome.

**Fig. S27.**
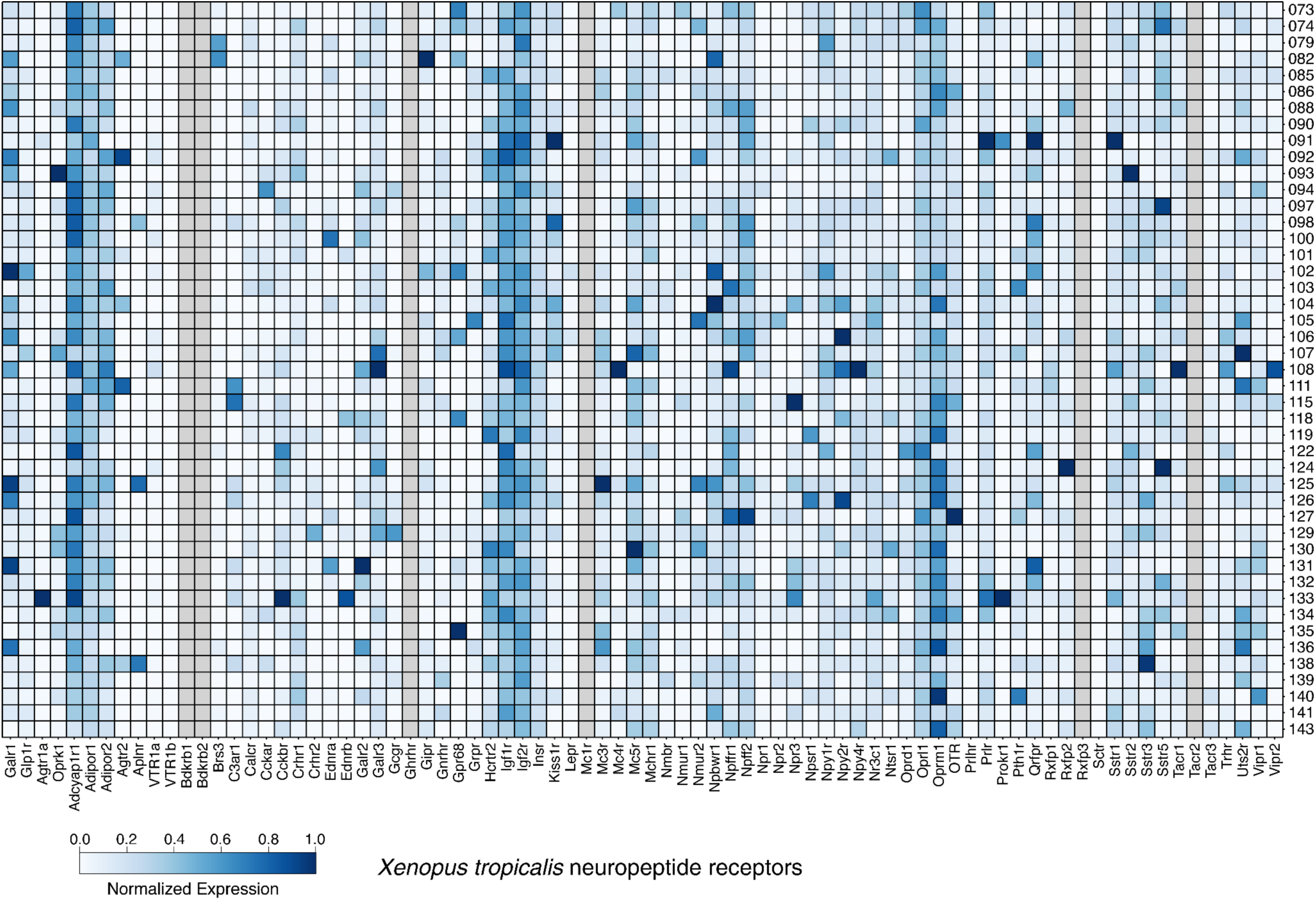
Neuropeptide receptor expression in *Xenopus tropicalis* subclasses. Expression of neuropeptide receptors with orthologs in 4 of 6 species in mapped subclasses. Bottom, names of selected neuropeptide receptors. Right, IDs for all mapped hypothalamic subclasses. Color represents min-max normalized expression across hypothalamic subclasses in the corresponding cell type (scale at the left). Grey indicates that the gene is either not expressed, or no ortholog is found in the genome.

**Fig. S28.**
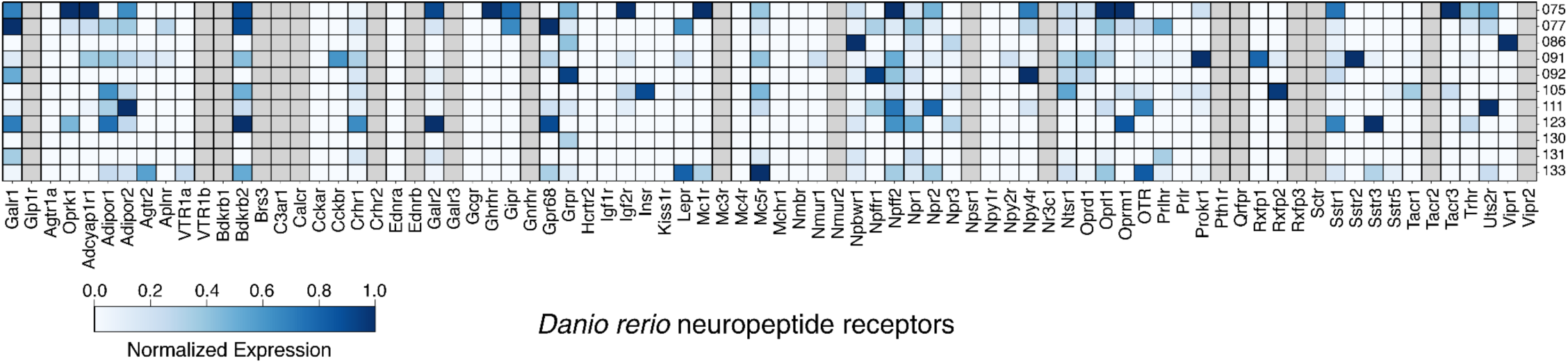
Neuropeptide receptor expression in *Danio rerio* subclasses. Expression of neuropeptide receptors with orthologs in 4 of 6 species in mapped subclasses. Bottom, names of selected neuropeptide receptors. Right, IDs for all mapped hypothalamic subclasses. Color represents min-max normalized expression across hypothalamic subclasses in the corresponding cell type (scale at the left). Grey indicates that the gene is either not expressed, or no ortholog is found in the genome.

**Fig. S29.**
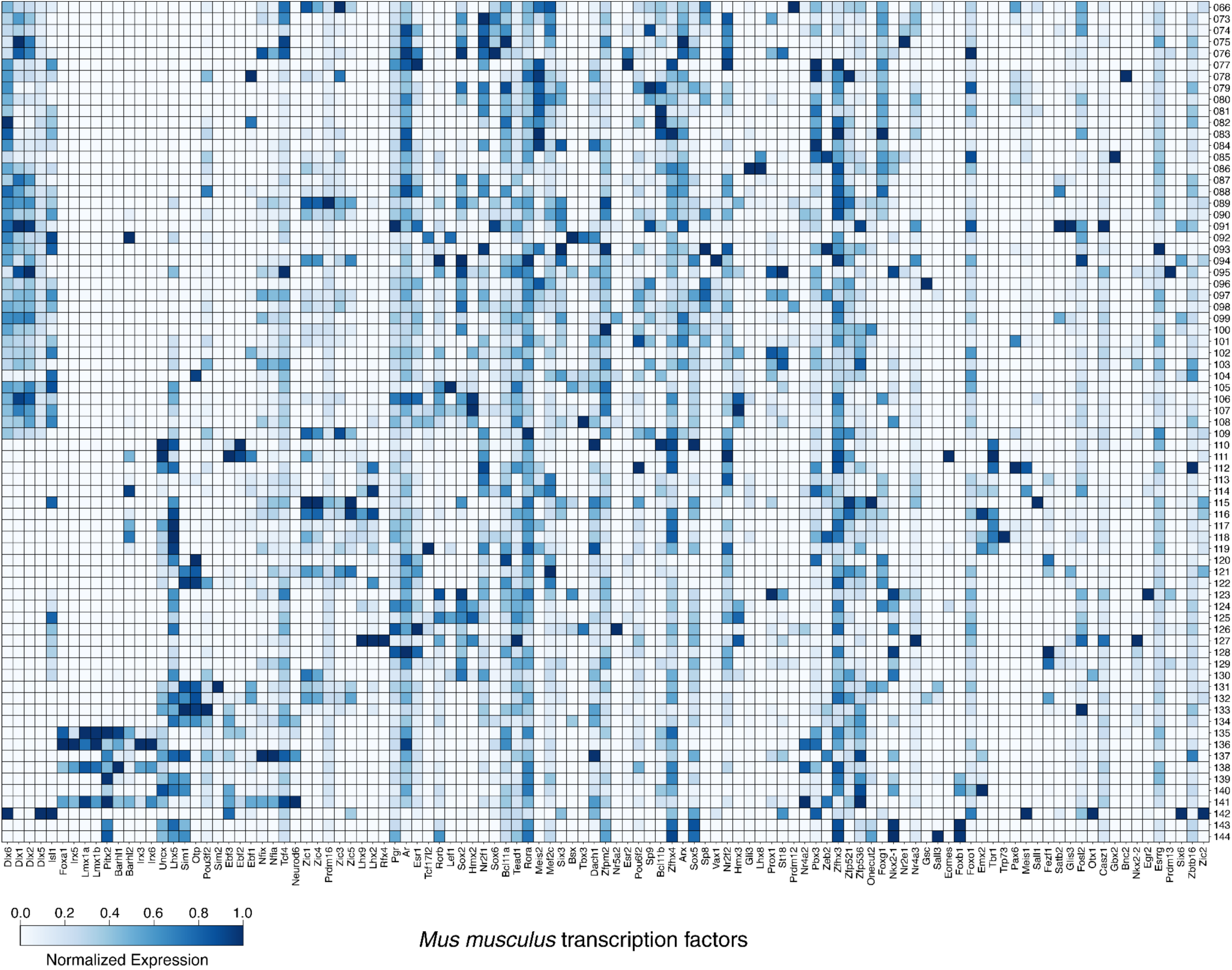
Transcription factor expression in *Mus musculus* subclasses. Expression of TFs that are enriched in a subclass across at least 4 species in mapped subclasses. Bottom, names of selected transcription factors. Right, IDs for all mapped hypothalamic subclasses. Color represents the min-max normalized expression across hypothalamic subclasses in the corresponding subclass (scale at the bottom). Grey indicates that the gene is either not expressed, or no ortholog is found in the genome.

**Fig. S30.**
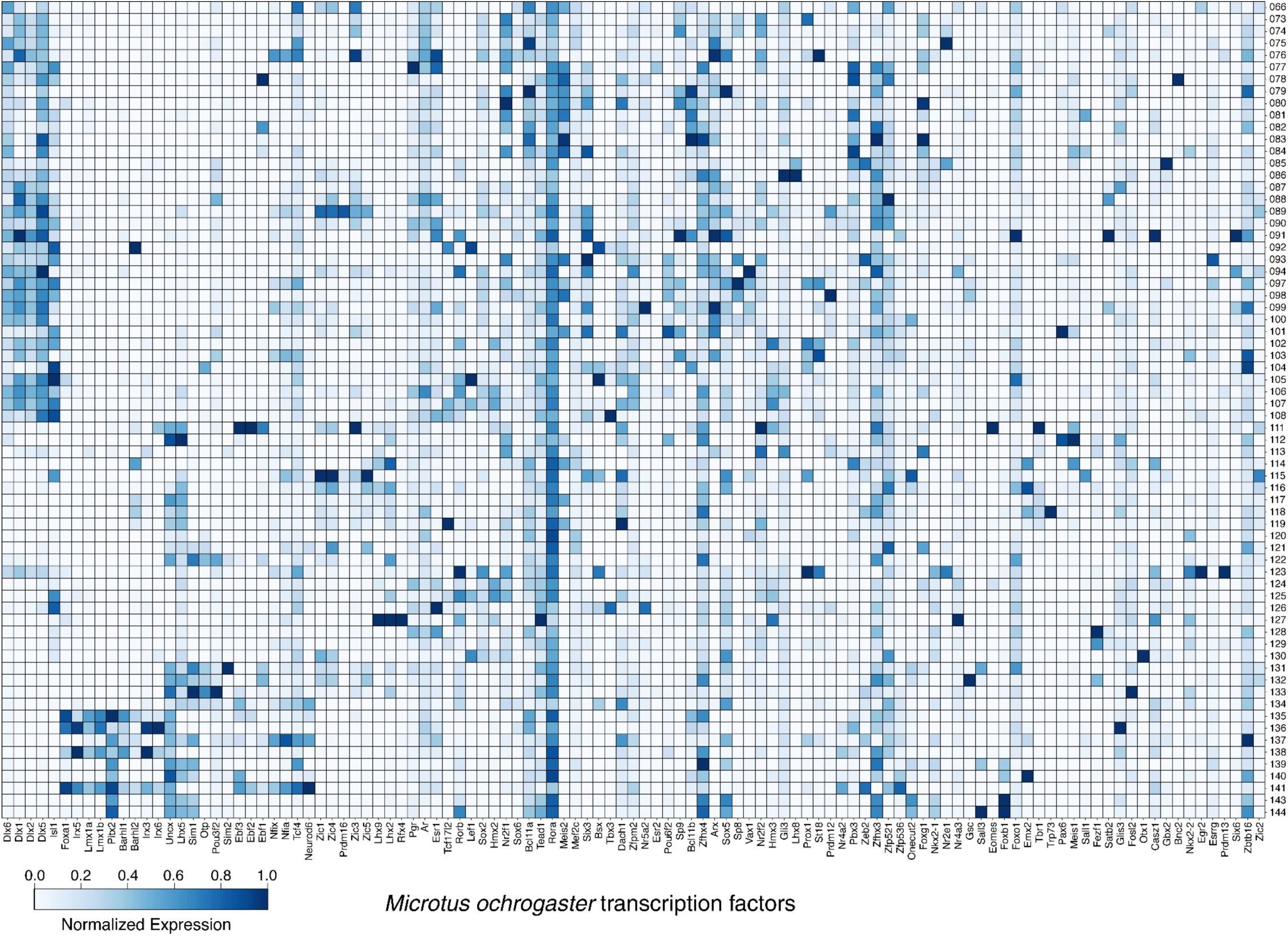
Transcription factor expression in *Microtus ochrogaster* subclasses. Expression of TFs that are enriched in a subclass across at least 4 species in mapped subclasses. Bottom, names of selected transcription factors. Right, IDs for all mapped hypothalamic subclasses. Color represents the min-max normalized expression across hypothalamic subclasses in the corresponding subclass (scale at the bottom). Grey indicates that the gene is either not expressed, or no ortholog is found in the genome.

**Fig. S31.**
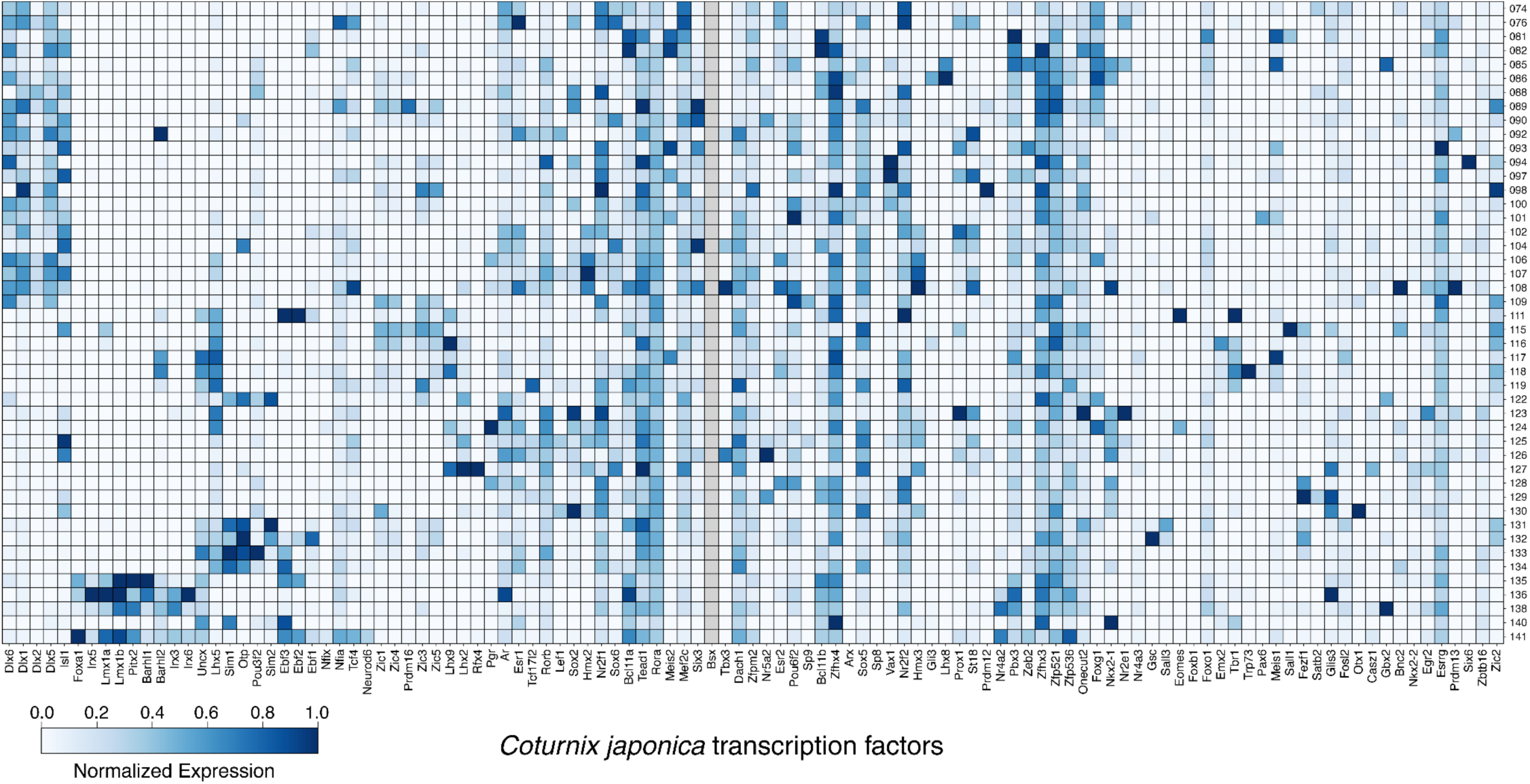
Transcription factor expression in *Coturnix japonica* subclasses. Expression of TFs that are enriched in a subclass across at least 4 species in mapped subclasses. Bottom, names of selected transcription factors. Right, IDs for all mapped hypothalamic subclasses. Color represents the min-max normalized expression across hypothalamic subclasses in the corresponding subclass (scale at the bottom). Grey indicates that the gene is either not expressed, or no ortholog is found in the genome.

**Fig. S32.**
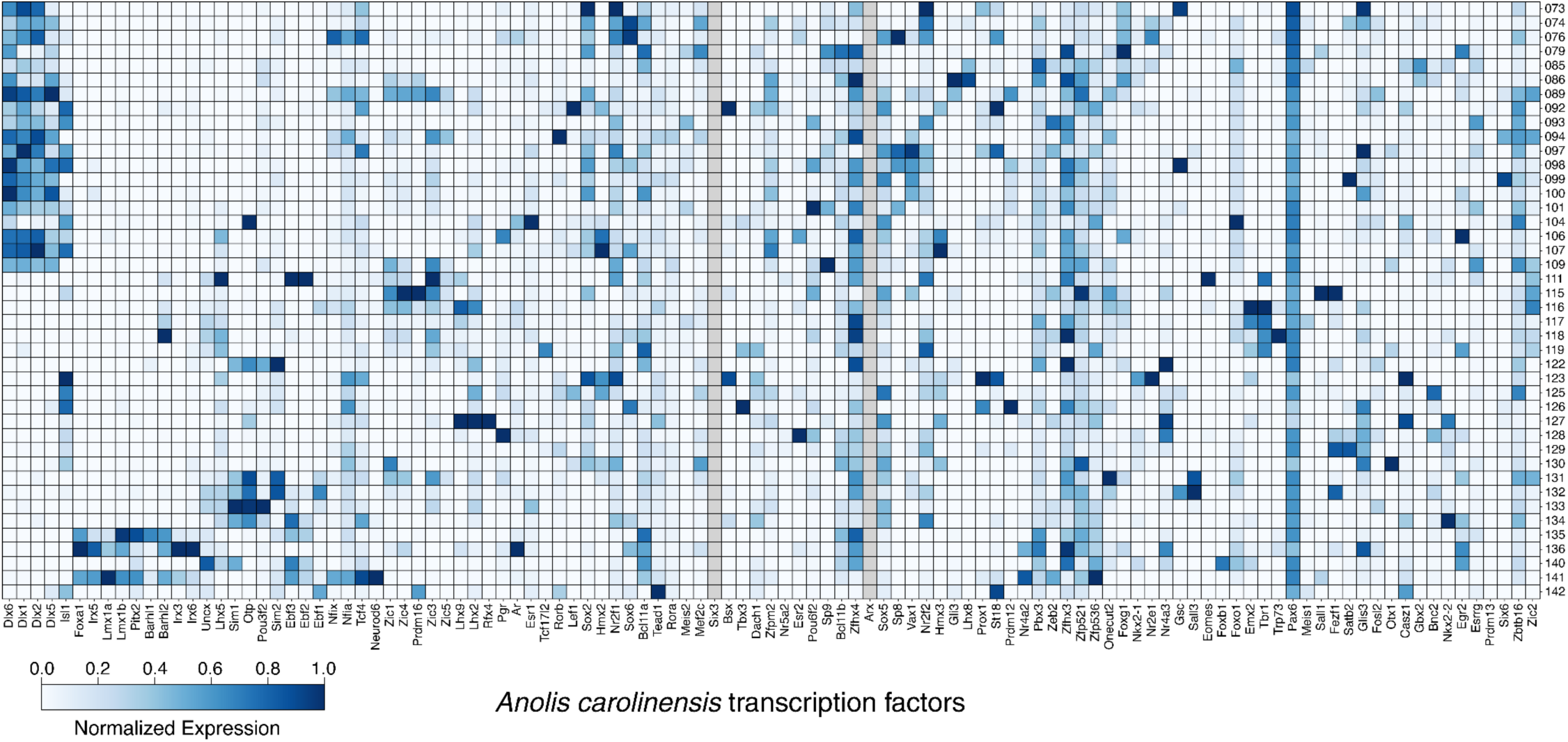
Transcription factor expression in *Anolis carolinensis* subclasses. Expression of TFs that are enriched in a subclass across at least 4 species in mapped subclasses. Bottom, names of selected transcription factors. Right, IDs for all mapped hypothalamic subclasses. Color represents min-max normalized expression across hypothalamic subclasses in the corresponding subclass (scale at the bottom). Grey indicates that the gene is either not expressed, or no ortholog is found in the genome.

**Fig. S33.**
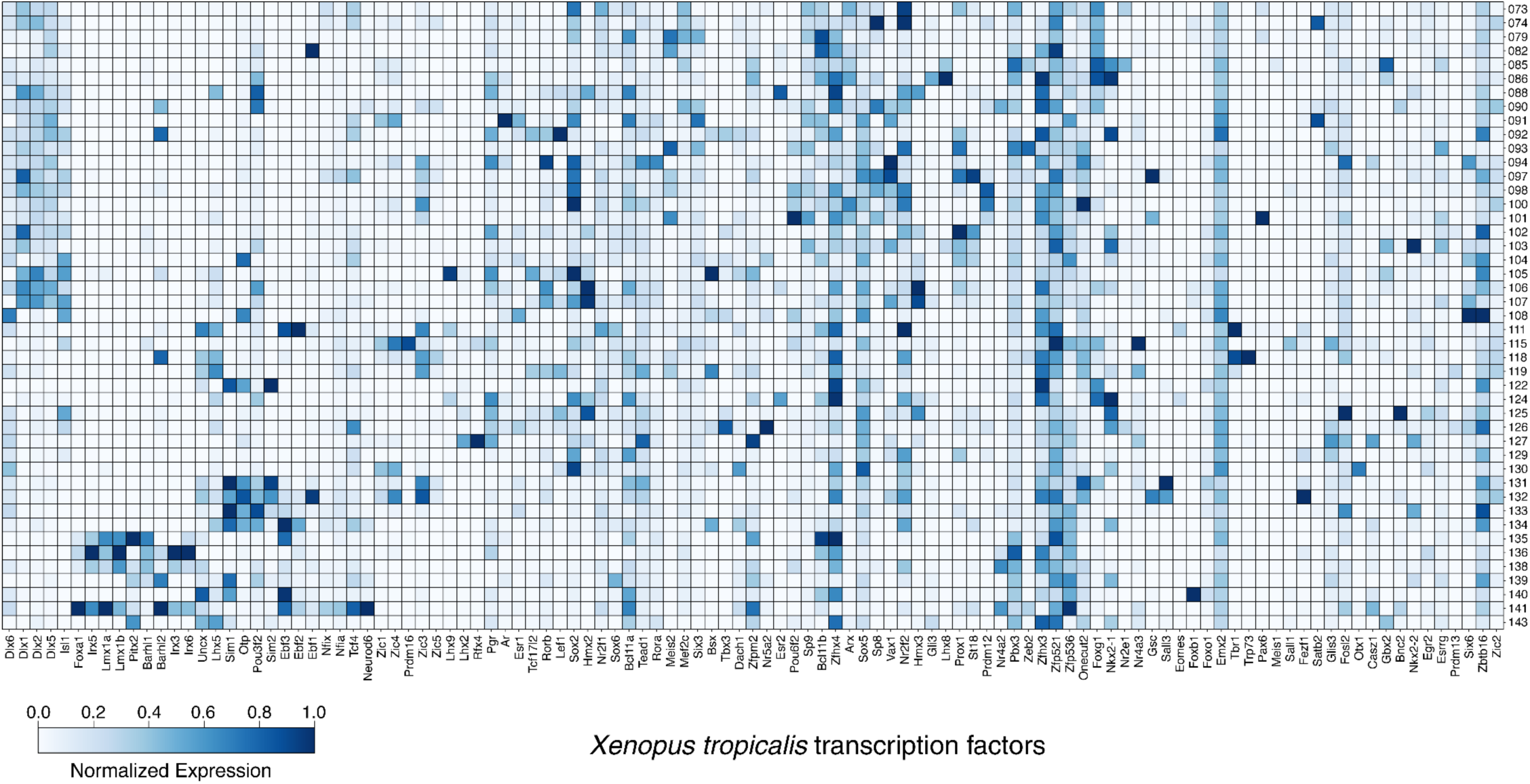
Transcription factor expression in *Xenopus tropicalis* subclasses. Expression of TFs that are enriched in a subclass across at least 4 species in mapped subclasses. Bottom, names of selected transcription factors. Right, IDs for all mapped hypothalamic subclasses. Color represents the min-max normalized expression across hypothalamic subclasses in the corresponding subclass (scale at the bottom). Grey indicates that the gene is either not expressed, or no ortholog is found in the genome.

**Fig. S34.**
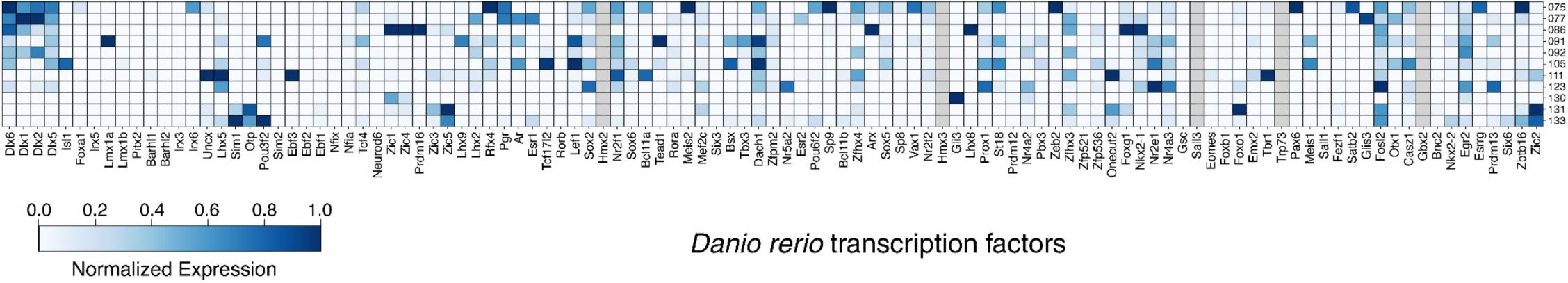
Transcription factor expression in *Danio rerio* subclasses. Expression of TFs that are enriched in a subclass across at least 4 species in mapped subclasses. Bottom, names of selected transcription factors. Right, IDs for all mapped hypothalamic subclasses. Color represents the min-max normalized expression across hypothalamic subclasses in the corresponding subclass (scale at the bottom). Grey indicates that the gene is either not expressed, or no ortholog is found in the genome.

**Fig. S35.**
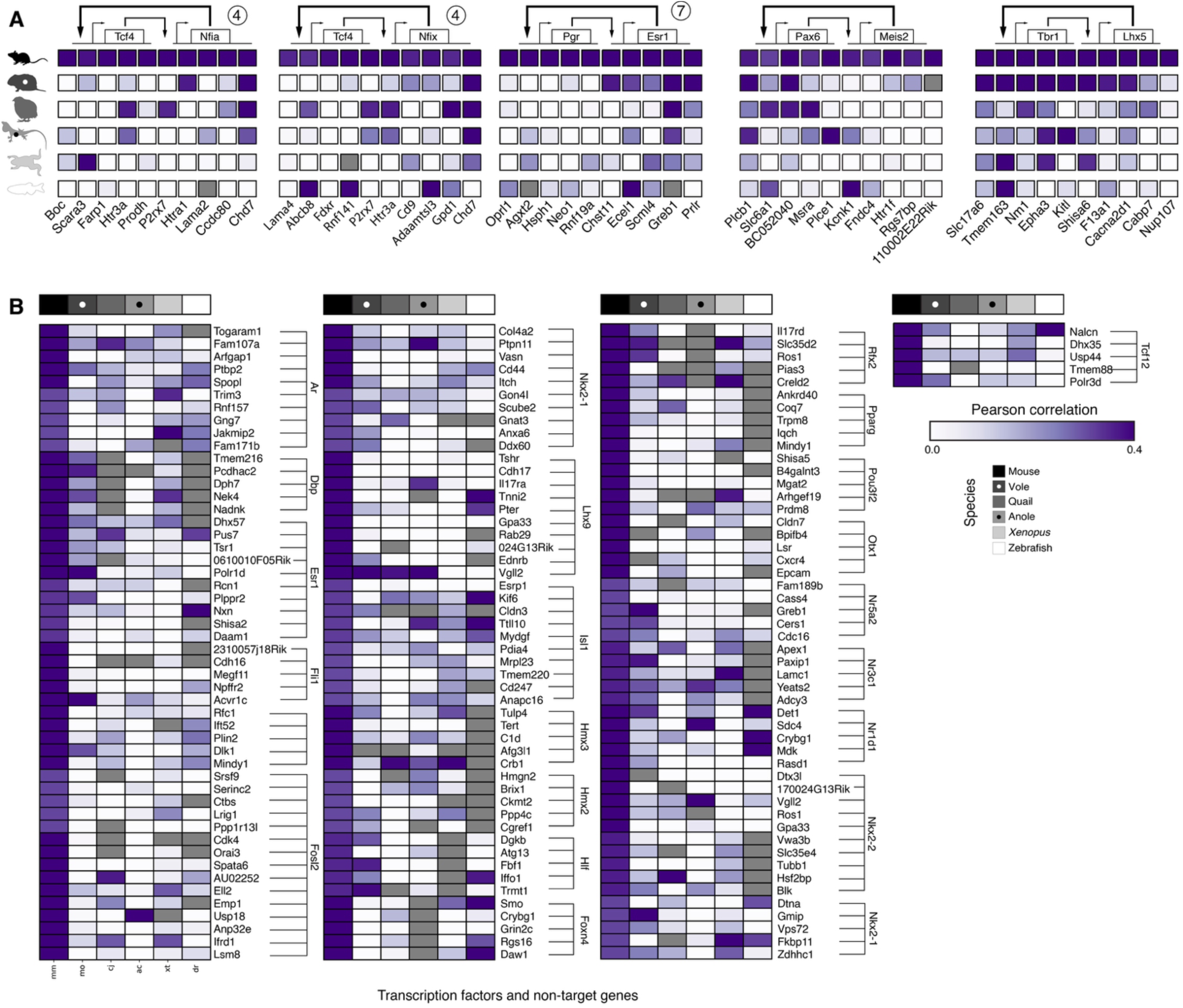
Correlation profile of target and non-target genes co-expressed with TFs in Fig 3E. and Fig. 3G. **(A)** Cross-species correlation for 8 conserved co-regulatory TF pairs and downstream genes identified using SCENIC+. Boxes are gray where the gene has no ortholog or no expression in the corresponding species. The remaining 8 conserved co-regulatory TF pairs are each composed of two paralogs (e.g., Dlx1–Dlx2), making their regulatory relationships difficult to disentangle. We therefore did not include them. Tbr1–Lhx5 and Pax6–Meis2 are not visualized in Fig. 3 due to only having 2 genes in the module, thus they have no module number. **(B)** Non-target genes are chosen because their correlation is comparable to the correlation between TFs and the neuropeptides they regulate (**Fig. 3G**; **Methods**). Grey indicates that either the TF or its nontarget gene is not expressed, or that no ortholog could be found in the species’ genome.

**Fig. S36.**
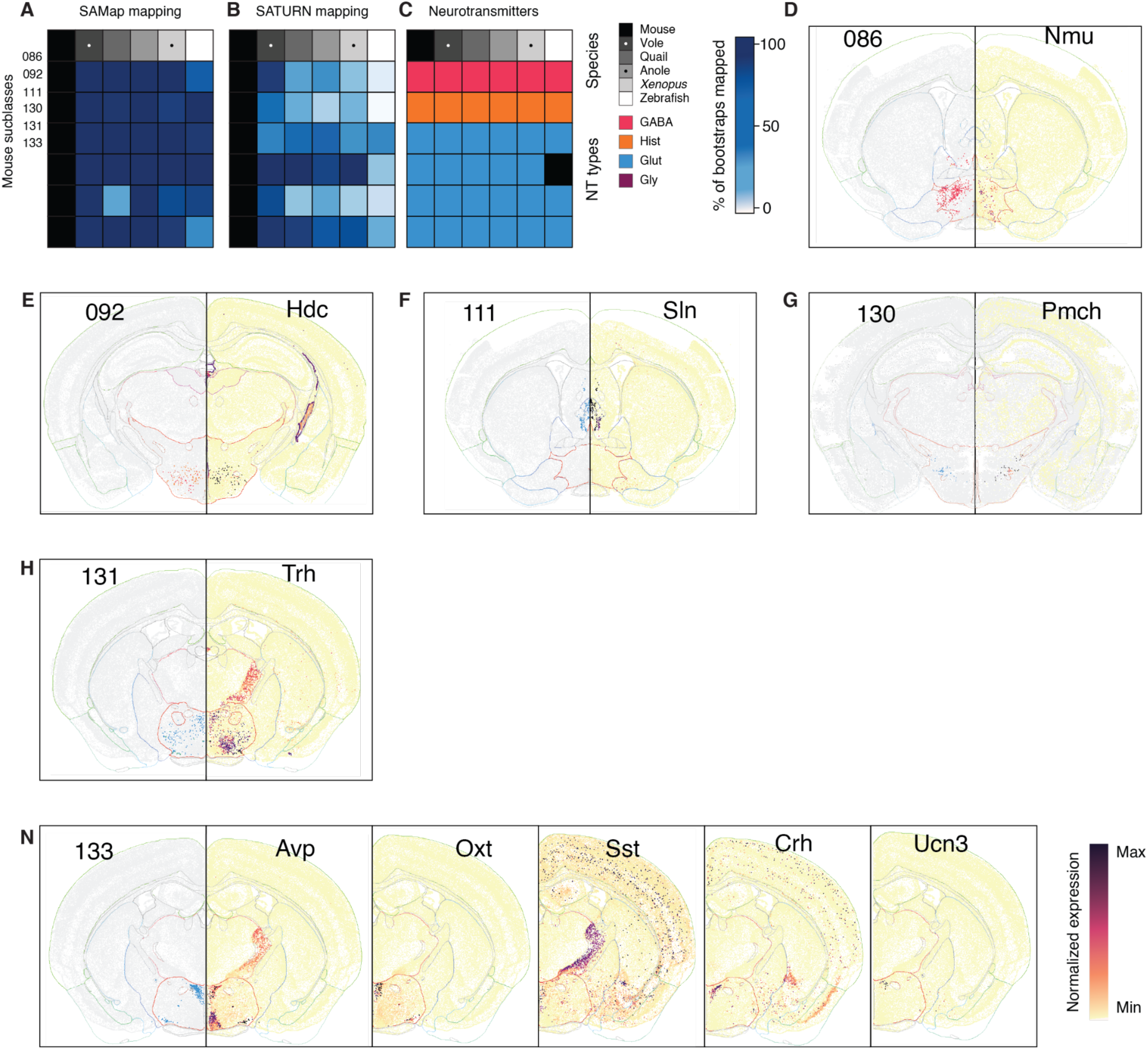
Highly conserved eleven hypothalamic subclasses across the vertebrate lineage. **(A–C)** SAMap (A) and SATURN (B) mapping and neurotransmitter types (C) of the six conserved hypothalamic subclasses across all six species. **(D–N)** Location (left panels) and marker gene expression (right panels) of the six subclasses in mouse based on the Allen Brain Atlas (*21*). The color in the left panels indicates the neurotransmitter types (same color code as C). The color in the right panel indicates the normalized expression levels. The range of the color intensity was normalized to the minimal and maximal value of the gene expression levels within the same image. The hypothalamic regions were outlined in red.

**Fig. S37.**
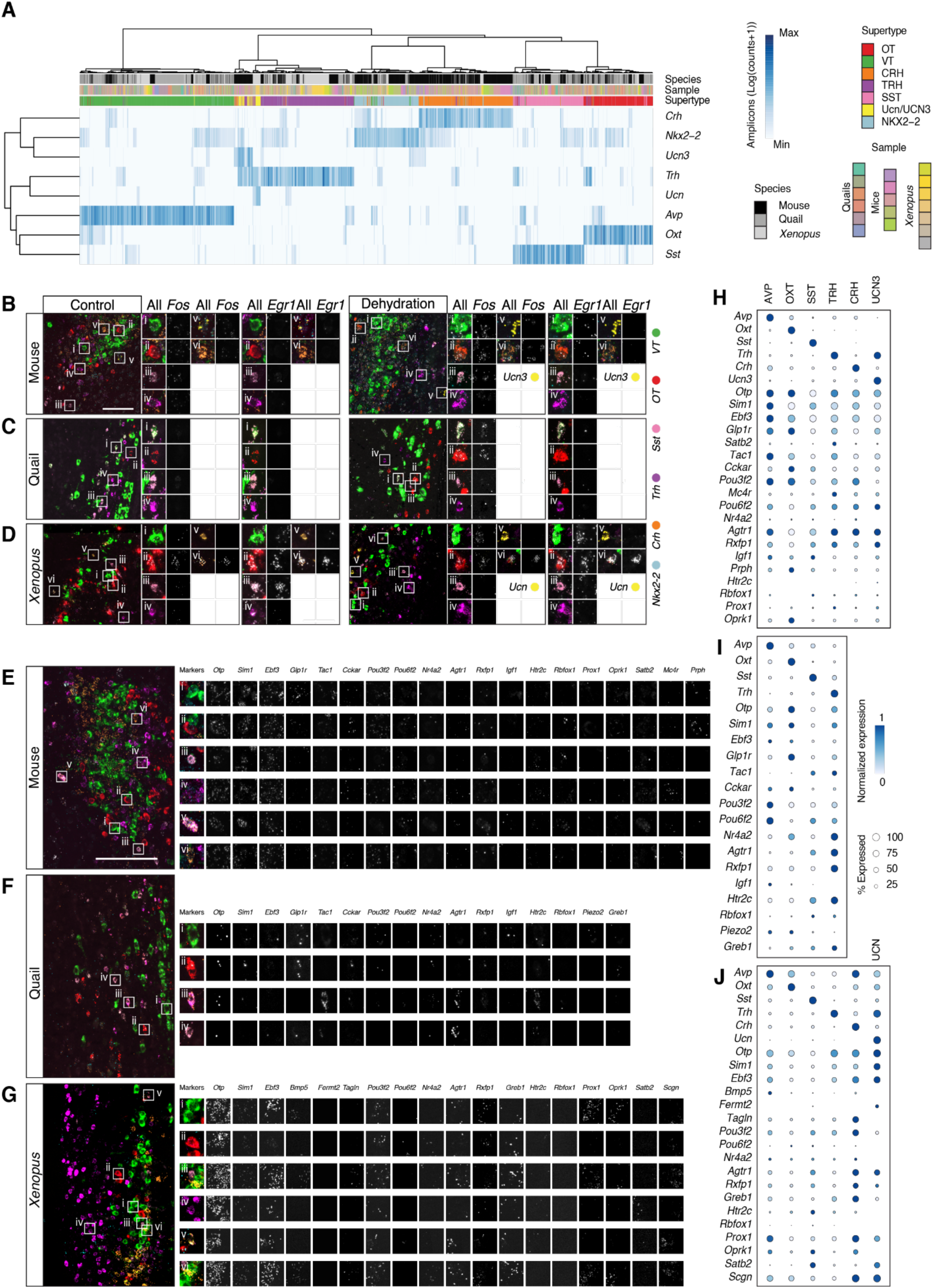
Additional analysis of STARmap data in Fig. 4. **(A)** Heatmap showing the expression of marker genes across mouse, quail, and *Xenopus* and across annotated supertypes. Each column indicates one cell. Color intensity indicates the log normalized amplicon counts. The first row indicates the species. The second row indicates individual samples. The third row indicates the annotated supertype. **(B–D)** Representative images showing *Egr1* expression in the same samples in **Fig. 4G–I**. Dots represent amplicons. Inset, representative cells expressing supertype markers, *Fos*, and *Egr1*. Scale bar, 100 µm. **(E–G)** Representative images showing additional genes detected from the STARmap libraries in PVH sections of mouse (E), quail (F), and xenopus (G). Scale bar, 100 µm. (**H–J**) Dotplot showing the expression level of the genes detected by the STARmap from the samples in panels E–G. Columns represent supertypes. Rows represent genes. Color intensity represents normalized amplicon count values. Darker blue indicates higher counts. Dot size represents the percentage of cells expressing the gene.

**Fig. S38.**
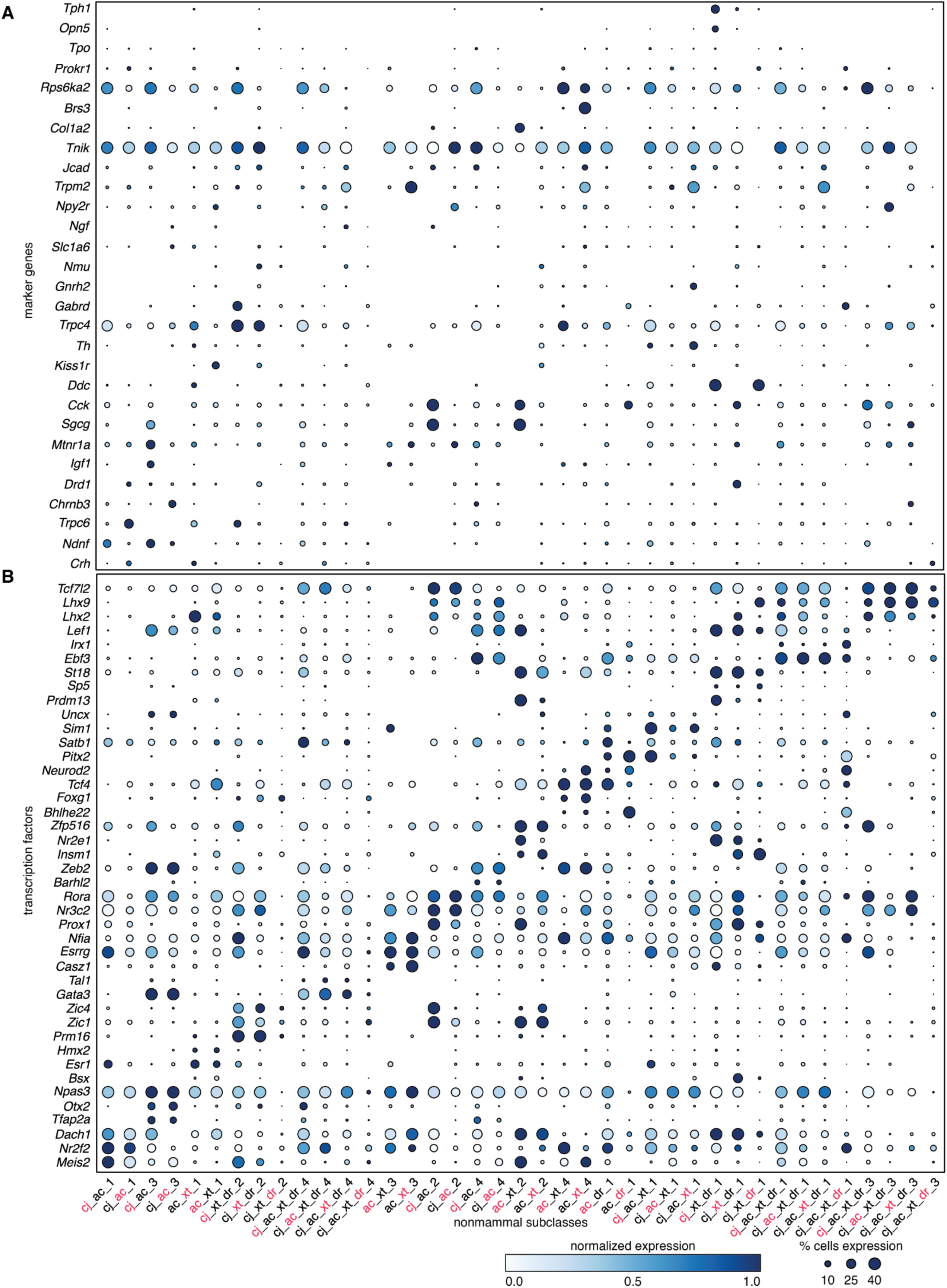
Additional analysis of non-mammalian subclasses. Min-max normalized expression of effectors (**A**) and TFs (**B**) in non-mammalian subclasses. The list of TFs is the same as in **Fig. 5c**. Effectors were manually collected and organized. Bottom, names of the non-mammalian subclasses. Red, the species for which the gene expression is plotted. Left, gene names.

**Fig. S39.**
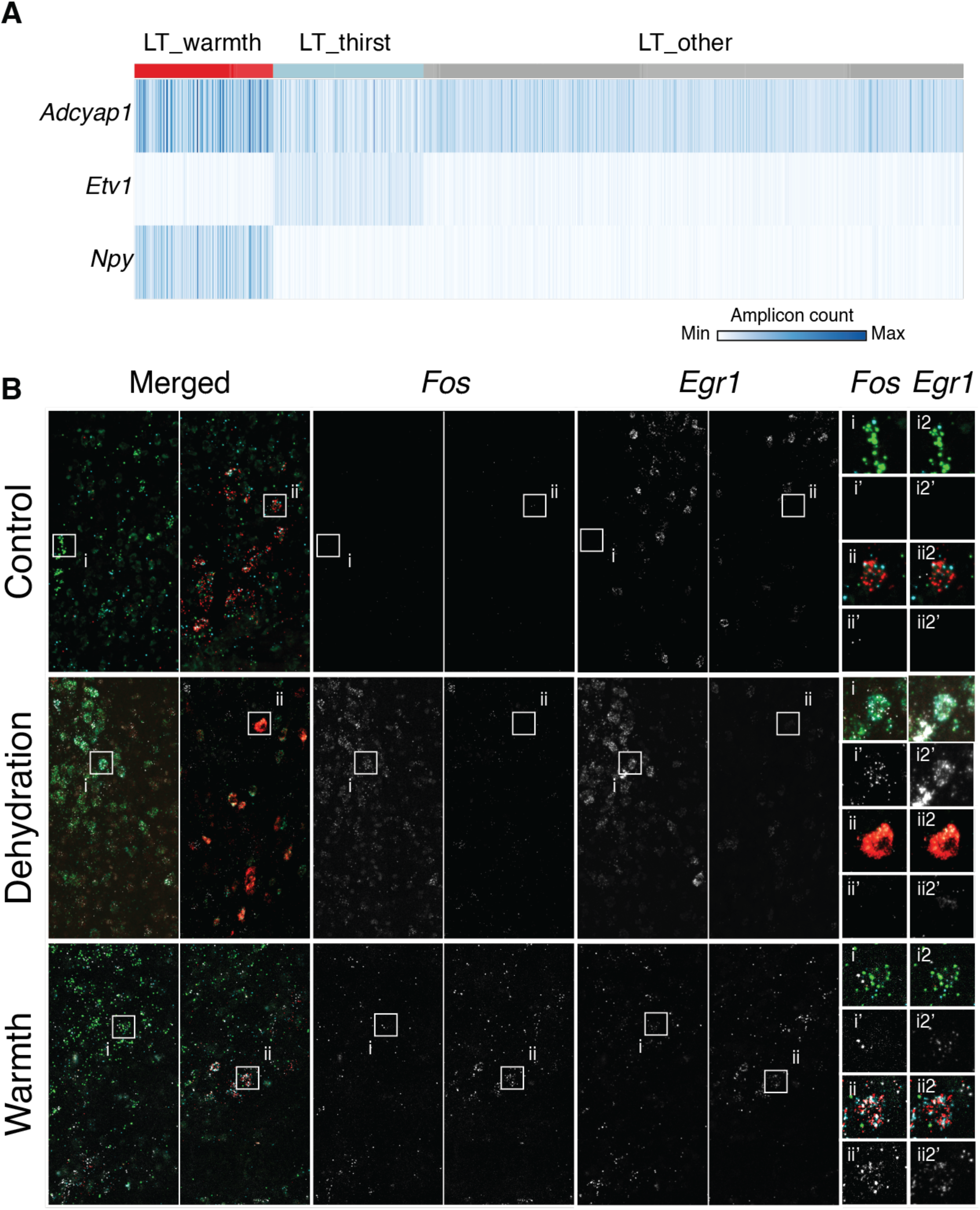
Additional analysis of STARmap data in Figure 6. **(A)** Amplicon counts for *Adcyap1*, *Etv1*, and *Npy* in the annotated LT supertypes. **(B)** *Egr1* amplicons in the same sample as in **Fig. 6E–G**.

**Table S1.**
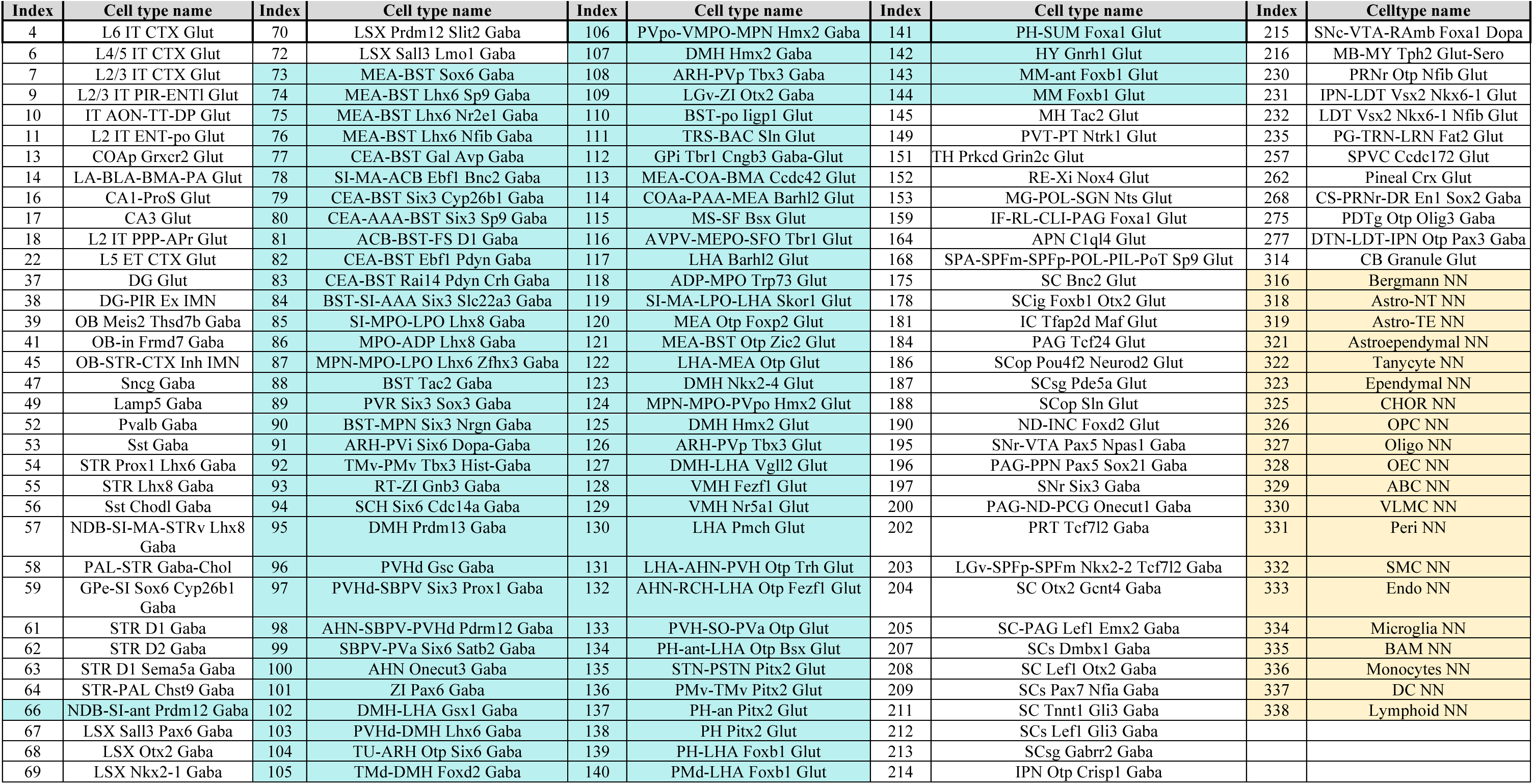
Allen Institute subclass names and the numerical IDs. Nomenclature is from (*21*) and the Allen Institute. Subclasses corresponding to hypothalamic and related extended amygdala neurons are highlighted in cyan. The other listed subclasses correspond to mapped cells from at least one species in our study, including glial cells (highlighted in yellow).

**Table S2.** TF correlation across thresholds. TF pairs are considered to be conserved if their correlation is higher than the threshold in 5 of the 6 species (**Methods**). Control genes are collected as described in **Methods**. Number of conserved co-expressed TFs significantly lower than conserved co-expressed control genes at a low correlation threshold (0.25), not significantly different at a medium threshold (0.30/0.35), and significantly higher at high correlation thresholds (0.40).

| Pearson correlation threshold | # Conserved co-expressed TFs | # Conserved co-expressed control | # Co-expressed TFs (any species) | % Conserved TFs |
| --- | --- | --- | --- | --- |
| 0.25 | 642 | 1,023 (95% C.I.: 763–1,283) | 54,196 | 1.18 |
| 0.30 | 267 | 346 (95% C.I.: 245–456) | 43,625 | 0.61 |
| 0.35 | 109 | 97 (95% C.I.: 59–142) | 34,070 | 0.32 |
| 0.40 | 55 | 22 (95% C.I.: 10–39) | 25,775 | 0.21 |

| Spearman correlation threshold | # Conserved co-expressed TF | # Conserved co-expressed control | # Co-expressed TFs (any species) | % Conserved TFs |
| --- | --- | --- | --- | --- |
| 0.25 | 871 | 1,219 (95% C.I.: 925– 1,497) | 65,540 | 1.33 |
| 0.30 | 298 | 300 (95% C.I.: 201– 406) | 49,595 | 0.60 |
| 0.35 | 96 | 56 (95% C.I.: 30– 86) | 35,413 | 0.27 |
| 0.40 | 43 | 8 (95% C.I.: 2 – 17) | 24,289 | 0.18 |

**Table S3.**
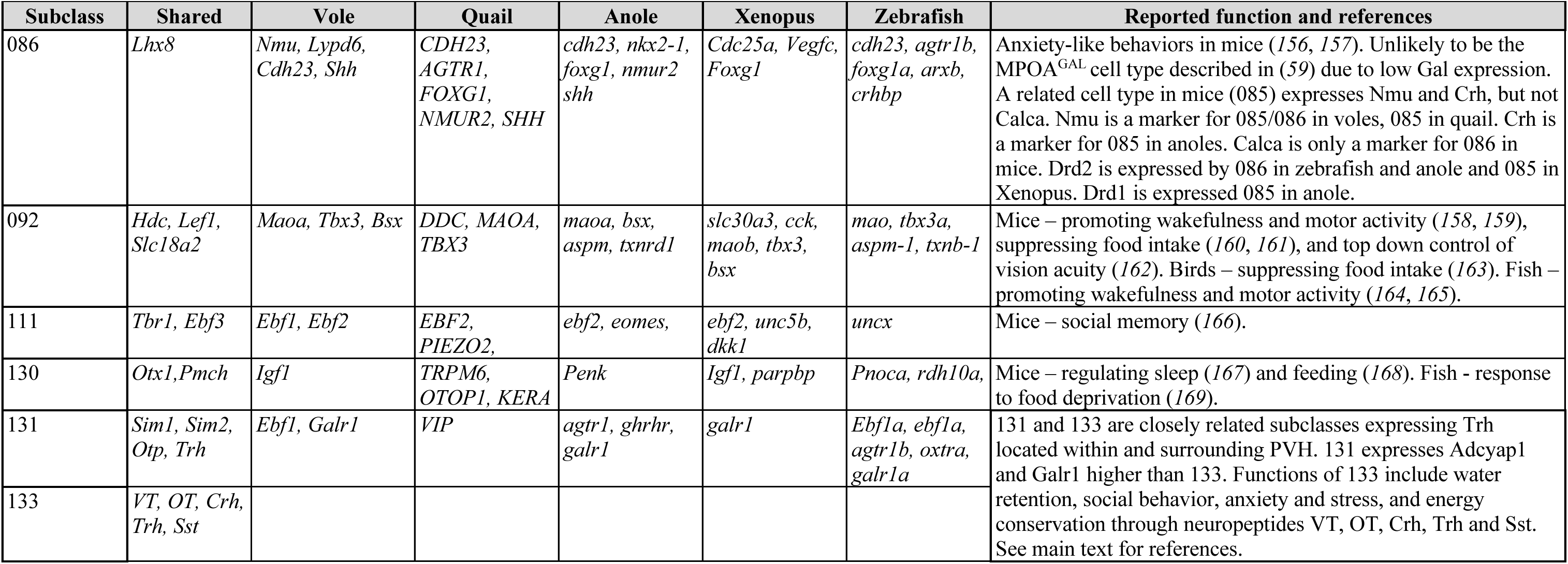
Additional marker genes and literatures of the highly conserved subclasses across vertebrates in Table 1.

**Table S4.** Genomes used in this study.

| Species | Repository | Genome | Annotation |
| --- | --- | --- | --- |
| *Microtus ochrogaster* | ENSEMBL | GCA_000317375.1 | Microtus_ochrogaster.MicOch1.0.112 |
| *Coturnix Japonica* | ENSEMBL | GCA_001577835.1 | Coturnix_japonica.Coturnix_japonica_2.0.105 |
| *Anolis Carolinesis* | NCBI | GCA_035594765.1 | GCF_035594765.1-RS_2024_02 |
| *Xenopus Tropicalis* | ENSEMBL | GCA_000004195.4 | Xenopus_tropicalis.UCB_Xtro_10.0.112 |
| *Ranitomeya Imitator*<br>*(draft)Ranitomeya Imitator (draft)* | Zenodo | doi/10.5281/zenodo.21312279 | Ranitomeya imitator dovetail assembly and annotation |
| *Ranitomeya Imitator (final)* | NCBI | GCA_032444005.1 | GCF_032444005.1-RS_2024_10 |
| *Danio Rerio* | NCBI | GCA_000002035.4 | GCF_000002035.6-RS_2024_08 |

**Table S6.**
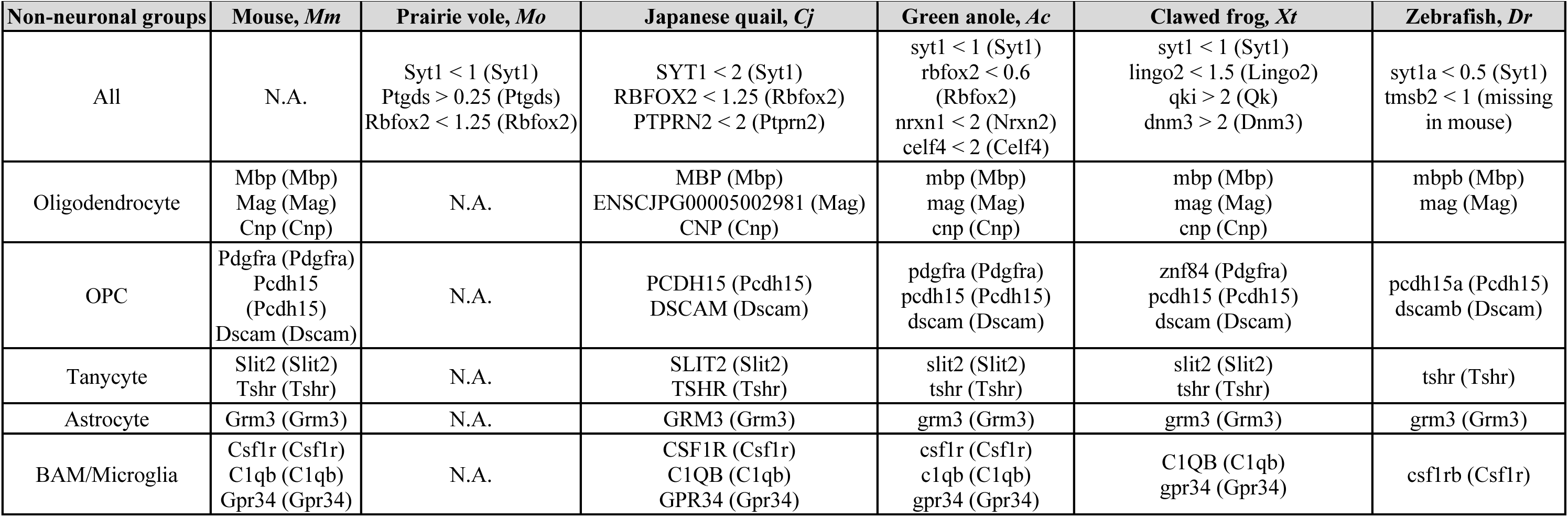
Non-neuronal thresholds and markers per species. Top row: Markers and thresholds for separating non-neuronal cells and neurons. Non-neuronal cells were defined as subclasses that did not pass two of the species’ specific thresholds, except in anole where the subclass needed to pass 3 of the thresholds. Markers were chosen based on genes described previously with mouse homolog in parentheses. Bottom rows: markers for each of the five non-neuronal populations found in **Fig. 2E**. Markers were selected manually.

**Table S7.**
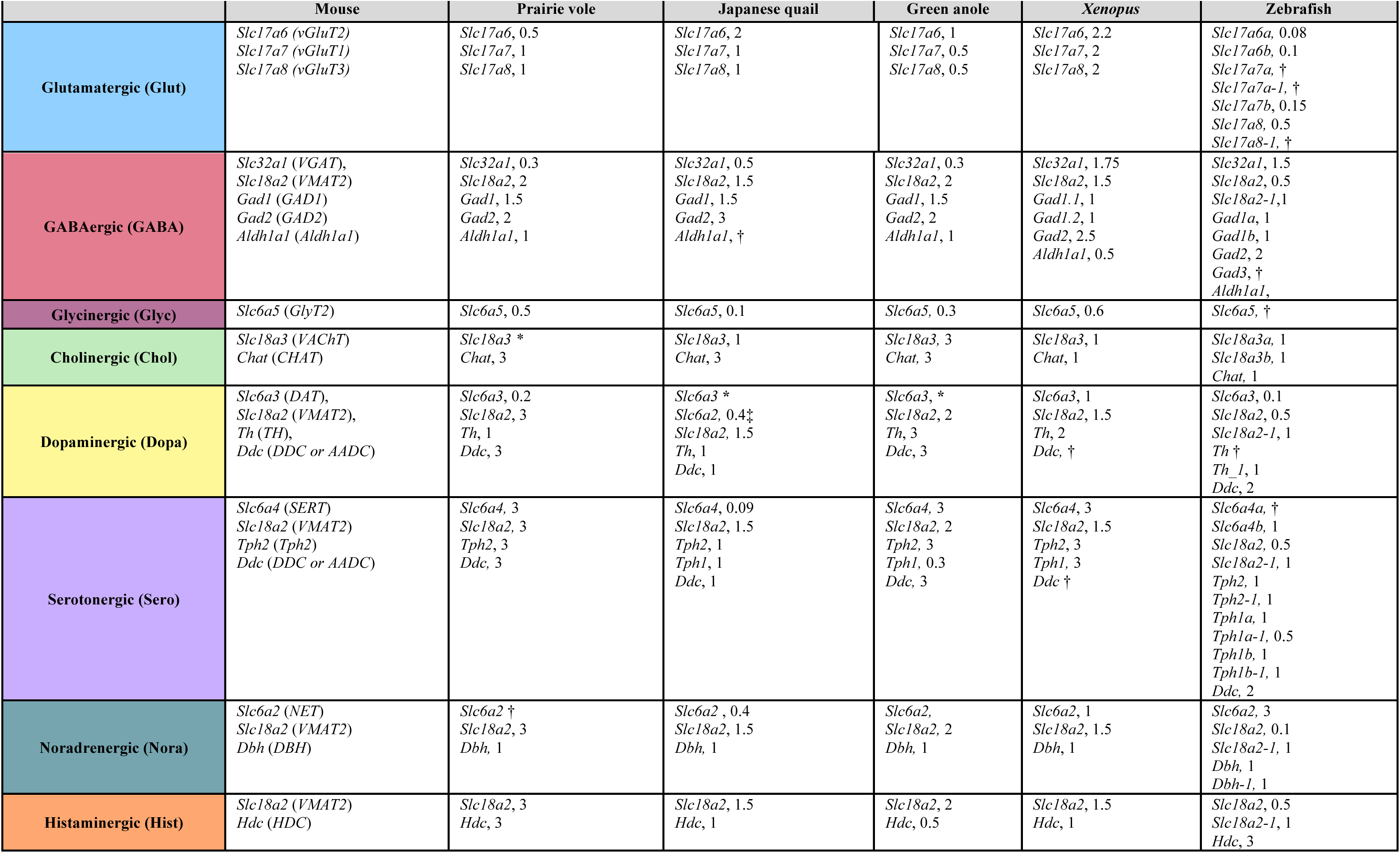
Genes and thresholds used to determine neurotransmitter types. Common or alternative names are shown in parentheses under the mouse column. Values are shown in ln(CPM+1). A subclass could be labeled as double positive for glutamatergic and GABAergic because it contained both glutamatergic and GABAergic supertypes while the individual cell was only glutamatergic or GABAergic but not both. One example of individual cells expressing both *Slc32a1* and *Slc17a6* was mouse subclass 112 shown in **Fig. 2**. * indicates that the gene was missing from the species’ genome annotation. † indicates that the gene was not detected. ‡ Dopaminergic cells in the quail expressed *Slc6a2*. There was no annotated *Slc6a3* in the quail genome annotation.

**Titles for additional supplementary tables (separate Excel sheets)**

**Table S5. Preprocessing thresholds (separate Excel sheet).** Thresholds used for preprocessing of raw snRNA-seq data are compiled, including minimum genes and UMIs per nucleus, ambient RNA cutoffs, and species-specific UMI thresholds where applicable.

**Table S8. Gene names used for plotting in this study (separate Excel sheet).** Assigned orthologs across all species for all genes described in the study. When more than one orthologs were found, one with the most similar expression pattern was used for plotting purposes. (see procedures described in **Methods)**. Empty cells represent orthologs not found. All genes in figures were labeled with mouse gene names unless otherwise specified.

**Table S9. List of neuropeptides and neuropeptide receptors used in this study (separate Excel sheet).** Neuropeptides and neuropeptide receptors are named with their mouse gene name. The spreadsheet is organized by neuropeptide with the receptors for the neuropeptide in the same row.

**Table S10. List of transcription factors (separate Excel sheet).** List of TFs used throughout the study. TFs are named with their mouse gene names. Annotation of TF family included for genes shown in **Fig. 3A**.

**Table S11. List of all TF modules, based on Pearson and Spearman correlation (separate Excel sheet).** All 272 unique TF modules are groups of TFs with a Pearson correlation above 0.3 in 5 or more species.

**Table S12. STARmap oligo sequences (separate Excel sheet).** This file includes the sequences of the oligos used in sequential visualization, detecting genes for determining neurotransmitter types in *Xenopus* (**Fig. S10**); detecting genes expressed in quail arcuate (**Fig. 3**); detecting genes expressed in quail, mouse, and *Xenopus* PVH (**Fig. 4**); and detecting genes expressed in quail LT (**Fig. 6**).

